# Dependence of diffusion in *Escherichia coli* cytoplasm on protein size, environmental conditions and cell growth

**DOI:** 10.1101/2022.02.17.480843

**Authors:** N. Bellotto, J. Agudo-Canalejo, R. Colin, R. Golestanian, G. Malengo, V. Sourjik

## Abstract

Inside prokaryotic cells, passive translational diffusion typically limits the rates with which cytoplasmic proteins can reach their locations. Diffusion is thus fundamental to most cellular processes, but the understanding of protein mobility in the highly crowded and non-homogeneous environment of a bacterial cell is still limited. Here we investigated the mobility of a large set of proteins in the cytoplasm of *Escherichia coli*, by employing fluorescence correlation spectroscopy (FCS) combined with simulations and theoretical modeling. We conclude that cytoplasmic protein mobility could be well described by Brownian diffusion in the confined geometry of the bacterial cell and at the high viscosity imposed by macromolecular crowding. We observed similar size dependence of protein diffusion for the majority of tested proteins, whether native or foreign to *E. coli*, and, for the faster-diffusing proteins, it is well consistent with the Stokes-Einstein relation once taking into account the specific dumbbell shape of protein fusions. Pronounced subdiffusion and hindered mobility are only observed for proteins with extensive interactions within the cytoplasm. Finally, while protein diffusion becomes markedly faster in actively growing cells, at high temperature, or upon treatment with rifampicin, and slower at high osmolarity, all of these perturbations affect proteins of different sizes in the same proportions, which could thus be described as changes of a well-defined cytoplasmic viscosity.

## Introduction

Diffusion of molecules is important for the function of any cellular system, setting the upper limit for the mobility of proteins and other (macro)molecules and for the rates of many biochemical reactions that rely on random encounters between molecules (Schavemaker, Boersma, and Poolman 2018). Although the fundamental physics of diffusion in dilute aqueous solutions is well understood and mathematically described (Einstein 1906; Langevin 1908; Perrin 1909), diffusion in a cellular environment may be quite different (Schavemaker, Boersma, and Poolman 2018; Mika and Poolman 2011). The concentration of macromolecules in the bacterial cytoplasm, primarily proteins but also ribonucleic acids (RNAs), a phenomenon known as macromolecular crowding, is extremely high. For *Escherichia coli*, it is around 300 mg/mL, which corresponds to a volume fraction of 25-30% (Cayley et al. 1991; Zimmerman and Trach 1991). Such macromolecular crowding could hinder free diffusion and influence kinetics of protein association and of gene expression (Klumpp et al. 2013; Tabaka et al. 2014; van den Berg, Boersma, and Poolman 2017). The effects of crowding on protein diffusion have been demonstrated both *in vitro* and *in vivo* (Dix and Verkman 2008; Rivas and Minton 2016). Compared to water, the diffusion of a free green fluorescent protein (GFP) was reported to be three to four times slower in the eukaryotic cytoplasm (Swaminathan, Hoang, and Verkman 1997) and up to ten times slower in the bacterial cytoplasm (Elowitz Michael et al. 1999; Nenninger, Mastroianni, and Mullineaux 2010; Mika et al. 2010; Kumar, Mommer, and Sourjik 2010).

In addition to the high density of macromolecules, the diversity in the size and chemical properties of the solutes makes the cytoplasmic environment highly inhomogeneous (Luby-Phelps 1999; Spitzer and Poolman 2013). How much the diffusion of a particular molecule is affected by macromolecular crowding might thus depend on the size (Muramatsu and Minton 1988) and shape of the molecule (Balbo et al. 2013) as well as on the nature of the crowders (Banks and Fradin 2005; Goins, Sanabria, and Waxham 2008). The effects of crowding observed in living cells appear to be even more complex, varying not only with the properties of the diffusing particle but also with the physiological state of the cell (Parry et al. 2014; Joyner et al. 2016) and the local cellular environment (Konopka et al. 2006; Persson, Ambati, and Brandman 2020). Moreover, non-trivial effects on diffusion arise due to reversible assembly and disassembly of the diffusing protein complexes (Agudo-Canalejo, Illien, and Golestanian 2020), and possibly also due to the active enhancement of enzyme diffusion by catalytic reactions (Golestanian 2015; Agudo-Canalejo et al. 2018; Zhang and Hess 2019).

The dependence of the diffusion coefficient (*D*) of a protein in the cytoplasm on its size might thus not necessarily follow the Stokes-Einstein (also called Stokes-Einstein-Sutherland-Smoluchowski) relation that is valid in dilute solutions, *D* ∝ *T*/(*ηR*) (Einstein 1906), where *T* is the absolute temperature in Kelvin, *η* the viscosity of the medium and *R* the hydrodynamic radius of the particle. For globular proteins, *R* is given by the radius of gyration (Tyn and Gusek 1990) and depends on the molecular mass (MM) as *R* ∝ MM^*β*^, where the exponent *β* would be 1/3 for perfectly compact and globular proteins but is in practice within the range of 0.35-0.43 for typical proteins, reflecting the fractal nature of the spatial distribution of protein mass (Smilgies and Folta-Stogniew 2015; Enright and Leitner 2005). Several studies of protein diffusion in the cytoplasm of *E. coli* have yielded different dependencies on the molecular mass, from ~0.33 (Nenninger, Mastroianni, and Mullineaux 2010) to ~2 (Kumar, Mommer, and Sourjik 2010), with an average *β* ~0.7 estimated based on the data pooled from multiple studies (Mika and Poolman 2011; Kalwarczyk, Tabaka, and Holyst 2012), and thus substantially steeper than predicted by the Stokes-Einstein relation. Similar exponent of ~0.7 was observed for limited sets of differently-sized proteins (Mika et al. 2010; Stracy et al. 2021). However, neither of these studies took explicitly into account the non-globularity of the used fluorescent constructs, where two or more proteins are typically connected by flexible linkers. For such multidomain proteins, shape fluctuations and hydrodynamic interactions between the different domains can have a sizeable effect on the effective diffusion coefficient of the whole protein (Agudo-Canalejo and Golestanian 2020), and they might thus be important to consider when interpreting deviations from the Stokes-Einstein relation.

Besides macromolecular crowding, the translational diffusion of cytoplasmic proteins is also influenced by intracellular structures, such as cytoskeletal filaments (Sabri et al. 2020), and by (transient) binding to other macromolecules (Saxton 2007; Guigas and Weiss 2008; von Bülow et al. 2019). Both these factors can not only reduce protein mobility but also lead to the anomalous subdiffusive behavior, where the mean square displacement (MSD) of diffusing particles does not scale linearly with time, as for Brownian diffusion in dilute solutions, but rather follows MSD ∝ *t^α^* with the anomalous diffusion exponent *α* being <1 (Saxton 1996; Etoc et al. 2018). Subdiffusion is commonly observed in eukaryotes, particularly at longer spatial scales, primarily due to the obstruction by the cytoskeletal filaments to the diffusion of proteins and larger particles (Di Rienzo et al. 2014; Sabri et al. 2020). The mobility of larger nucleoprotein (Golding and Cox 2004; Lampo et al. 2017) and multiprotein particles (Yu et al. 2018) in the bacterial cytoplasm is also subdiffusive, while the diffusion of several tested small proteins was apparently Brownian (Bakshi, Bratton, and Weisshaar 2011; English et al. 2011).

Even for the same protein, e.g. GFP or its spectral variants, estimates of the diffusion coefficient in the cytoplasm obtained in different studies vary widely (Schavemaker, Boersma, and Poolman 2018), which could be in part due to differences in methodologies. Most early studies in bacteria relied on fluorescence recovery after photobleaching (FRAP), where diffusion is quantified from the recovery of fluorescence in a region of the cell bleached by a high intensity laser (Loren et al. 2015). These measurements provided values of diffusion coefficient for GFP ranging from 3 μm^2^s^−1^ to 14 μm^2^s^−1^ (Elowitz Michael et al. 1999; Mullineaux et al. 2006; Konopka et al. 2009; Kumar, Mommer, and Sourjik 2010; Mika et al. 2010; Nenninger, Mastroianni, and Mullineaux 2010; Schavemaker, Smigiel, and Poolman 2017). More recently, single-particle tracking (SPT), where diffusion is measured by following the trajectories of single fluorescent molecules over time (Kapanidis, Uphoff, and Stracy 2018), became increasingly used. Lastly, diffusion can also be studied *in vivo* using fluorescence correlation spectroscopy (FCS) (Cluzel, Surette, and Leibler 2000), which measures the time required by a fluorescent molecule to cross the observation volume of a confocal microscope (Elson 2011). SPT and FCS measure protein mobility locally within the cell, with FCS having also a significantly better temporal resolution than FRAP and SPT. Both methods provided higher but still varying values of *D_GFP_*, from 8 μm^2^s^−1^ up to 18 μm^2^s^−1^ (Meacci et al. 2006; English et al. 2011; Sanamrad et al. 2014; Diepold et al. 2017; Rocha et al. 2019).

Protein mobility also depends on the environmental and cellular conditions that affect the structure of the bacterial cytoplasm (Schavemaker, Boersma, and Poolman 2018). Diffusion of large cytoplasmic particles, measured by SPT, was shown to be sensitive to the antibiotics-induced changes in the cytoplasmic crowding (Wlodarski et al. 2020) and to the energy-dependent fluidization of the cytoplasm (Parry et al. 2014). Protein diffusion is also affected by high osmolarity that increases macromolecular crowding and might create barriers to diffusion (Konopka et al. 2006; Konopka et al. 2009; Liu et al. 2019). Furthermore, the surface charge of cytoplasmic proteins has been shown to have a dramatic effect on their mobility (Schavemaker, Smigiel, and Poolman 2017).

Variations between values of diffusion coefficients observed even for the same model organism in different studies, each investigating only a limited number of protein probes, using different strains, growth conditions and measurement techniques, hampered drawing general conclusions about the effective viscosity of bacterial cytoplasm and its dependence on the protein size. Furthermore, while the impact of several physiological perturbations on protein diffusion has been established, most of these previous studies used either large particles or free GFP, and how these perturbations affect properties of the cytoplasm over the entire physiological range of protein sizes remained unknown.

Here, we address these limitations by systematically analyzing the mobility of a large number of differently-sized cytoplasmic fluorescent protein constructs under standardized conditions by FCS. We further combined experiments with Brownian dynamics simulations and theoretical modeling of diffusion to correct for effects of confined cell geometry. Our work establishes general methodology to analyze FCS measurements of protein mobility in a confined space, which could be broadly applicable to cellular systems.

For the majority of studied constructs, we observe consistent dependence of the diffusion coefficient on the protein size, with a pronounced upper limit on diffusion at a given molecular mass. When corrected for the confinement due to the bacterial cell geometry, the diffusion of these constructs was nearly Brownian. Moreover, part of the deviation of the mass-dependence of their diffusion coefficients from the Stokes-Einstein relation might be explained by the specific shape of the fusion proteins. The slower and more anomalous diffusion of several protein constructs was apparently due to their strong interactions with other cellular proteins and protein complexes, and disruption of these interactions restored a Brownian diffusion close to the upper limit expected for their mass. Proteins that are not native to *E. coli* were observed to diffuse very similarly to their *E. coli* counterparts, except for their motion being slightly subdiffusive. Under the same experimental conditions FCS and FRAP measurements yield similar values of diffusion coefficients, suggesting that no pronounced dependence of protein mobility on spatial scale could be observed in the bacterial cytoplasm. Finally, we investigated the effects of environmental osmolarity and temperature, of exposure to antibiotics and of cell growth on the mobility of proteins of different size, demonstrating that the effects of all these perturbations, including cell growth, on protein diffusion could be simply explained by changes in a unique cytoplasmic viscosity.

## Results

### Dependence of cytoplasmic protein mobility on molecular mass measured by FCS

For our analysis of cytoplasmic protein mobility, we generated a plasmid-encoded library of 31 cytoplasmic proteins (Figure 1 – supplementary table 1 and supplementary table 2) of *E. coli* fused to superfolder GFP (sfGFP) (Pédelacq et al. 2006). We selected proteins that, according to the available information, are not known to bind DNA or to form homomultimers, although we did not exclude *a priori* proteins that interact with other proteins. The structure of all selected proteins is known and roughly globular, avoiding effects of the irregular protein shape on mobility. These proteins belong to different cellular pathways. The expected size and stability of each construct was verified by gel electrophoresis and immunoblotting (Figure 1 – figure supplement 1). Only one of the constructs, ThpR-sfGFP, showed >20% degradation to free sfGFP, and it was therefore excluded from further analyses. This was also the sole construct with an atypically high isoelectric point (pI), and all remaining constructs have pI ranging from 5.1 to 6.2, as common for cytoplasmic proteins (Schwartz, Ting, and King 2001). We further imaged the distribution of fusion proteins in the cytoplasm. Except for RihA-sfGFP and NagD-sfGFP that were subsequently excluded, all other constructs showed uniform localization (Figure 1A). Expression of most fusion proteins used for the measurements of diffusion had little effect on *E. coli* growth (Figure 1 – figure supplement 2), and even for several proteins that delayed the onset of the exponential growth, the growth rate around the mid-log phase when cultures were harvested for the analysis was similar.

**Figure 1.**
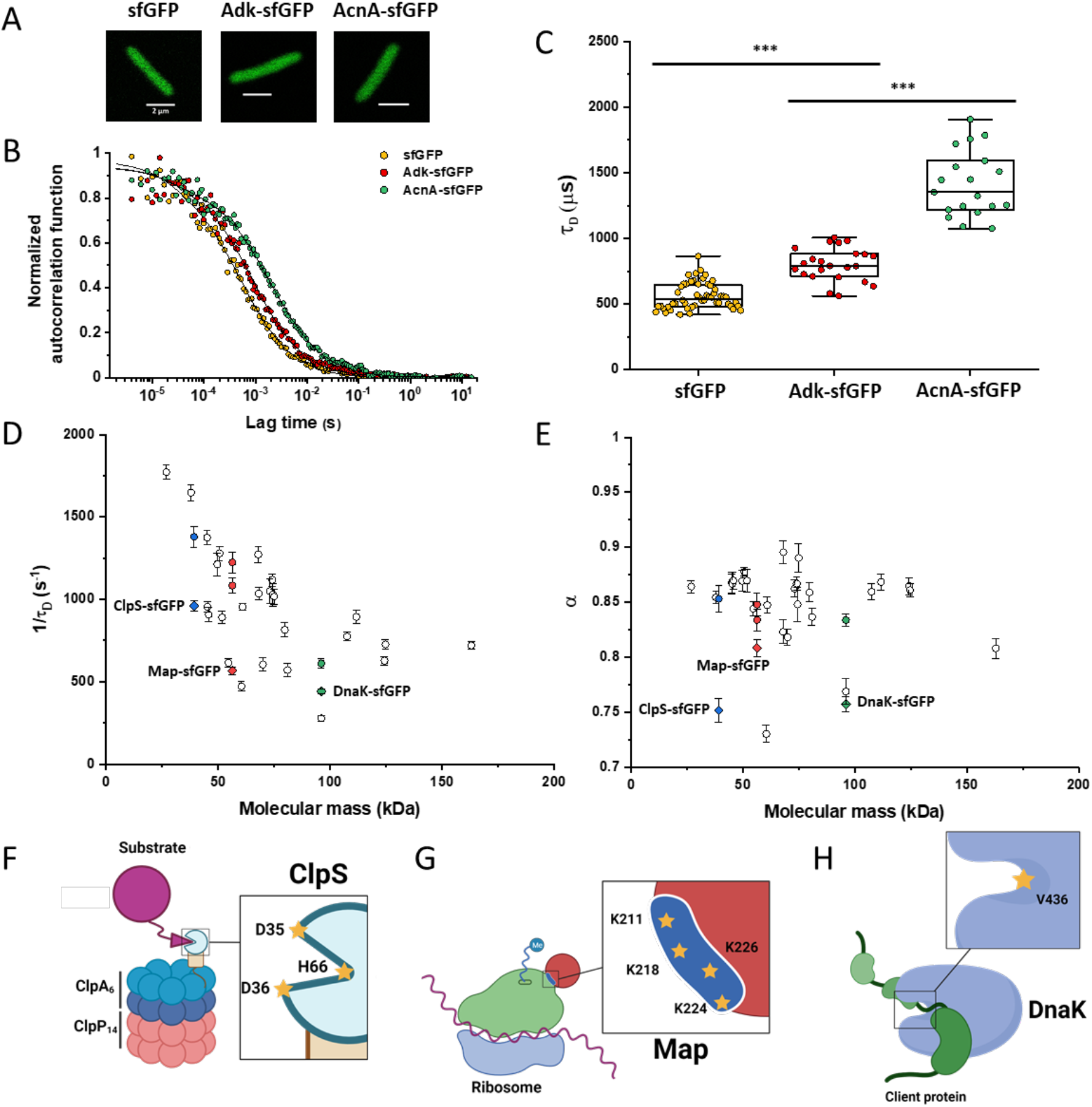
Dependence of protein mobility in bacterial cytoplasm on molecular mass and cellular interactions. (A) Examples of fluorescence microscopy images of *E. coli* cells expressing either sfGFP or the indicated sfGFP-tagged cytoplasmic proteins. Scale bar is 2 μm. (B) Representative autocorrelation functions (ACFs) measured by FCS for the indicated protein constructs. Data were fitted using the anomalous diffusion model (solid lines). All ACF curves were normalized to their respective maximal values to facilitate comparison. (C) Diffusion times (*τ_D_*) measured for the indicated protein constructs. Each dot in the box plot represents the value for one individual cell, averaged over six consecutive acquisitions (Figure 1 – figure supplement 3) *** p<0.0001 in a two-tailed heteroscedastistic *t*-test. Exact p-values for all significance analyses performed in this study can be found in Appendix 4. (D, E). Dependence of protein mobility (1/*τ_D_*; D) and apparent anomaly of diffusion (*α*; E) on molecular mass. Each symbol represents the average value for all individual cells that have been measured for that particular construct and the error bars represent the standard error of the mean. Individual values are shown in Figure 1 – figure supplement 5 and the numbers of measured cells for all the experiments performed in this study are shown in Appendix 5. Protein constructs with low mobility for which effects of specific interactions were further investigated are highlighted in color and labeled. The values of 1/*τ_D_* and *α* for both the original constructs (diamonds) and the constructs where mutations were introduced to disrupt interactions (circles) are shown. For Map, two alternative amino acid substitutions that disrupt its interaction with the ribosome are shown (see Figure 1 – figure supplement 10). (F-H) Cartoons illustrating the cellular interactions that could affect mobility of ClpS (F), Map (G), and DnaK (H). ClpS engages with the ClpAP protease and with substrates, cartoon adapted from (Roman-Hernandez et al. 2011). Map interacts with the actively translating ribosomes, cartoon adapted from (Sandikci et al. 2013). DnaK interacts with unfolded client protein. Amino acidic residues that were mutated to disrupt interactions are highlighted (see text for details). **Figure 1 – source data** Individual *τD* measurements from Figure 1C Individual mean and standard errors of the mean of 1/*τ_D_* values from Figure 1D Individual mean and standard errors of the mean of *α* values from Figure 1E

The mobility of the remaining 28 fusion constructs and of free sfGFP was investigated in living *E. coli* cells by FCS (see Materials and Methods and Appendix 1). In order to reduce the impact of photobleaching on FCS measurements, cell length was moderately (approximately twofold) increased by treatment with the cell-division inhibitor cephalexin for 45 min, yielding an average cell length of ~5 μm (Figure 1A). The resulting larger cell volume indeed reduces the rate of photobleaching. During each FCS measurement, the laser focus was positioned close to the polar region in the cell cytoplasm, in order to keep the confocal volume possibly away from both the cell membrane and the nucleoid, and the fluorescence intensity in the confocal volume was measured over time (Figure 1 – figure supplement 3). For each individual cell, six subsequent acquisitions of 20 s each were performed at the same position. The autocorrelation function (ACF) of the fluorescence intensity fluctuations was independently calculated for each time interval and fitted to extract the mobility parameters of the fluorescent proteins. Although we initially considered both the Brownian diffusion and the anomalous diffusion models, the latter model proved to be considerably better in fitting the experimental data (Figure 1 – figure supplement 4). The anomalous diffusion model was therefore used to determine the diffusion (or residence) time (*τ_D_*) of a fluorescent molecule in the confocal volume and the anomalous diffusion exponent *α* for all ACFs (Figure 1B and Figure 1 – supplementary table 1 and Figure 1 - figure supplement 3). The averaged values of *τ_D_* and *α* for each individual cell were then calculated from these six individual acquisitions (Figure 1C and Figure 1 – figure supplement 5). Although, as mentioned above, all finally used protein constructs showed no or little degradation, we tested a possible impact of the fraction of free sfGFP for the construct that displayed the strongest (~15%) degradation, DsdA-sfGFP. To this end, we fitted the FCS data using a model of two-components anomalous diffusion, where the weight of the fast component was fixed to 15% and its values of *τ_D_* and *α* to the average values obtained for sfGFP (Figure 1 – figure supplement 6). The average value of *τ_D_* for the slow component was only ~7% lower compared to our regular fit using the one-component model, and the value of *α* remained unchanged, suggesting that the impact of an even smaller fraction of free GFP for other constructs could also be neglected.

Despite their substantial intercellular variability, the obtained mean values of the diffusion time were clearly different between protein constructs (Figure 1C and Figure 1 – supplementary table 1). We next plotted the mean values of 1/*τ_D_*, which reflect protein mobility, against the molecular mass of protein constructs (Figure 1D). This dependence revealed a clear trend, where mobility of more than half of the constructs decreased uniformly with their molecular mass, while some exhibited much lower mobility than the other constructs of similar mass. In contrast, the anomalous diffusion exponent *α* showed no apparent dependence on the protein size, ranging from 0.8 to 0.86 for most of the constructs (Figure 1E). Notably, the few protein constructs with *α* of ~ 0.8 or lower were also among the ones with low mobility (Figure 1D and 1E, colored symbols). There was no significant correlation between the values of 1/*τ_D_* or *α* and the length or the width of individual cells, although a weak trend of *α* to increase with cell width might exist (Figure 1 – figure supplement 7). Furthermore, no significant effect of the treatment with cephalexin for cells of similar length (Figure 1 – figure supplement 8) could be observed.

### Macromolecular interactions reduce protein mobility

We reasoned that the main group of constructs that exhibit mobility close to the apparent mass-dependent upper limit represents proteins whose diffusion is only limited by macromolecular crowding, and that the lower 1/*τ_D_* and *α* of other constructs might be due to their specific interactions with other cellular proteins or protein complexes. Indeed, for three of these proteins (ClpS, Map, DnaK) such interactions are well characterized and can be specifically disrupted. ClpS is the adaptor protein that delivers degradation substrates to the protease ClpAP (Roman-Hernandez et al. 2011). The substrate-binding site of ClpS is constituted by three amino acid residues (D35, D36, H66) that interact with the N-terminal degron of target proteins (Figure 1F). If these residues are mutated into alanine, substrate binding *in vitro* is substantially reduced (Roman-Hernandez et al. 2011; Humbard et al. 2013). Additionally, ClpS directly docks to the hexameric ClpA. Consistently, we observed that while the stability of the mutant construct ClpS^D35A_D36A_H66A^-sfGFP was not affected (Figure 1 – figure supplement 9), its mobility in a *ΔclpA* strain became significantly higher and less anomalous, with both 1/*τ_D_* and *α* reaching levels similar to those of other proteins of similar mass (Figure 1D and 1E and Figure 1 – figure supplement 10).

Similar results were obtained for the other two constructs. Map is the methionine aminopeptidase that cleaves the N-terminal methionine from nascent polypeptide chains (Solbiati et al. 1999). Map interacts with the negatively charged backbone of ribosomes through four positively charged lysine residues (K211, 218, 224, 226) located in a loop (Figure 1G). If these residues are mutated into alanine, the *in vitro* affinity of Map for the ribosomes is reduced (Sandikci et al. 2013). The mobility of Map-sfGFP was indeed much increased by alanine substitutions at all four lysine sites (Figure 1D and 1E and Figure 1 – figure supplement 9 and figure supplement 10). Interestingly, charge inversion of lysines to glutamic acid did not further increase Map-sfGFP mobility as was expected based on *in vitro* experiments (Sandikci et al. 2013).

DnaK is the major bacterial chaperone that binds to short hydrophobic polypeptide sequences, which become exposed during protein synthesis, membrane translocation or protein unfolding (Genevaux, Georgopoulos, and Kelley 2007). DnaK accommodates its substrate peptides inside a hydrophobic pocket (Figure 1H). The substitution of the valine residue 436 with bulkier phenylalanine creates steric hindrance that markedly decreases substrate binding to DnaK *in vitro* (Mayer, Rüdiger, and Bukau 2000), and both the 1/*τ_D_* and *α* of DnaK^V436F^-sfGFP were significantly higher than for the correspondent wild-type construct (Figure 1D and 1E and Figure 1 – figure supplement 9 and figure supplement 10). Nevertheless, in this case the 1/*τD* did not reach the levels of other proteins of similar molecular mass, which is likely explained by multiple interactions of DnaK with other components of the cellular protein quality control machinery besides its binding to substrates (Kumar and Sourjik 2012).

### Apparent anomaly of diffusion could be largely explained by confinement

When FCS measurements are performed in a confined space with dimensions comparable to those of the observation volume, such confinement may affect the apparent mobility of fluorescent molecules (Gennerich and Schild 2000; Jiang et al. 2020). To investigate the effect of confinement on our FCS measurements, we performed Brownian dynamics simulations of FCS experiments with particles undergoing three-dimensional, purely Brownian diffusion inside a bacterial cell-like volume (Figure 2A *Inset*; see Materials and Methods). For the values of cell diameter commonly observed under our growth conditions, 0.8-0.9 μm, and over a wide range of particle diffusion coefficients, simulated autocorrelation functions could be indeed successfully fitted with the anomalous diffusion model, yielding an anomalous diffusion exponent of around 0.8-0.9 (Figure 2A and 2B). This made us hypothesize that the relatively small apparent deviation from Brownian diffusion in the fit, with *α* above 0.8 common to most constructs, may primarily reflect a confinement-induced effect rather than proper subdiffusion.

**Figure 2.**
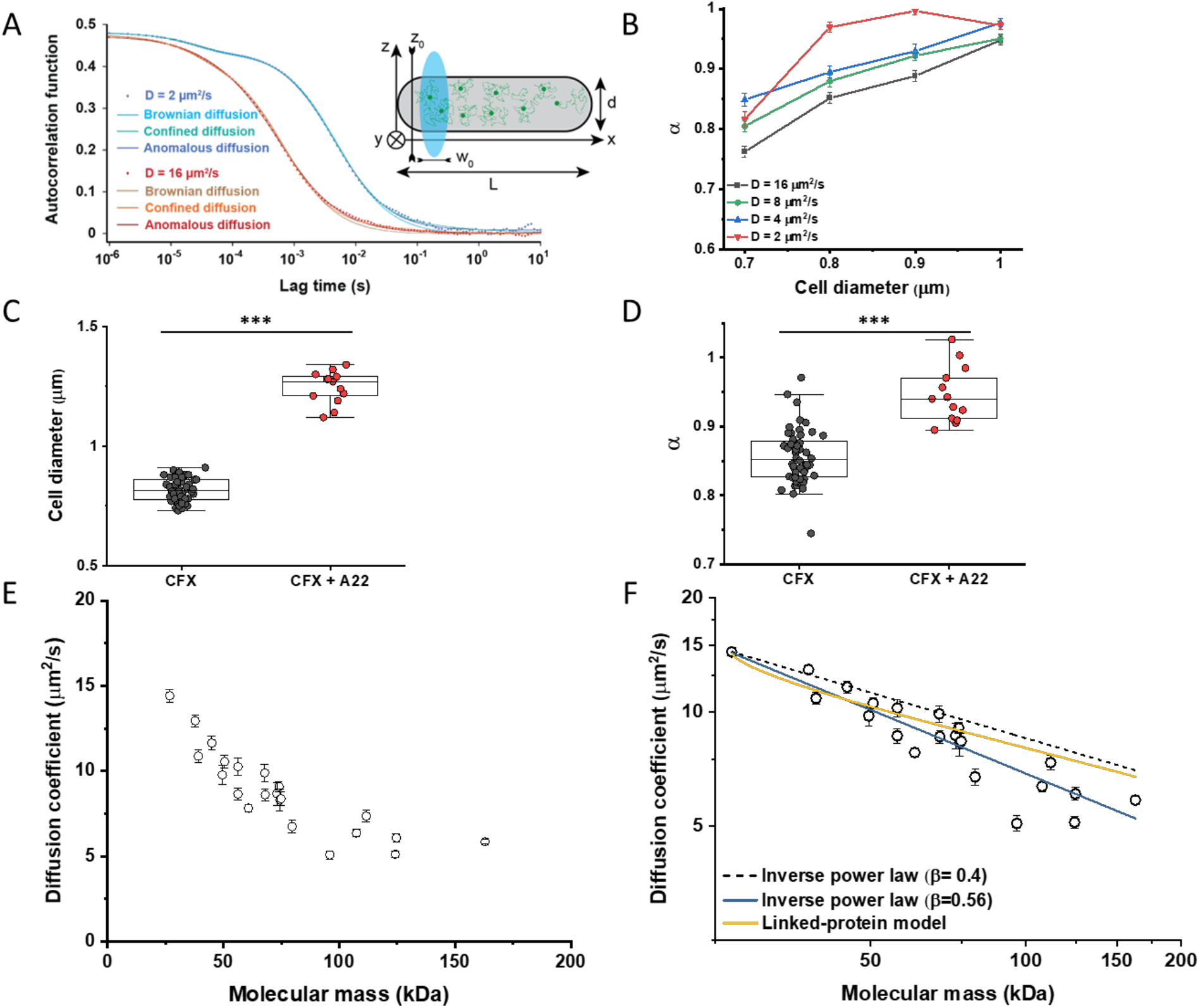
Protein diffusion in bacterial cytoplasm corrected for confinement. (A) Representative ACFs of simulated fluorescence intensity fluctuations. Simulations were performed in a confined geometry of a cell with indicated length *L* and diameter *d*, and dimensions of the measurement volume *ω_0_* and *z_0_*, representing an experimental FCS measurement (*Inset*; see Materials and Methods) for two different values of the ansatz diffusion coefficient. Solid lines are fits by the models of unconfined Brownian diffusion, anomalous diffusion and by the Ornstein-Uhlenbeck (OU) model of Brownian diffusion under confinement, as indicated. (B) The exponent *α* extracted from the fit of the anomalous diffusion model to the ACFs data that were simulated at different values of the cell diameter. Corresponding values of the diffusion coefficient are shown in Figure 2 – figure supplement 5. (C, D) *E. coli* cells treated with cephalexin alone or with cephalexin and 1 μg/mL of A22 (see Materials and methods), show A22-dependent increase in the measured cell diameter (C) and higher values of the exponent *α* extracted from the fit to the ACF measurements (D). *** p<0.0001 in a two-tailed heteroscedastistic *t*-test. (E) Dependence of the diffusion coefficient calculated from fitting the experimental ACFs with the OU model of confined diffusion. Only the subset of apparently freely diffusing constructs from Figure 1D has been analyzed with the OU model (see also Figure 1 – supplementary table 1). Each circle represents the average value for all individual cells that have been measured for that particular construct (Appendix 5), and the error bars represent the standard error of the mean. Error bars that are not visible are smaller than the symbol size. (F) Fit of the mass dependence with an inverse power law (solid blue line, exponent *β* = 0.56 ± 0.05), and predictions of the Stokes-Einstein relation (black dashed line) and of the model describing diffusion of two linked globular proteins (solid yellow line), both with exponent *β* = 0.4. **Figure 2 – source data** Average and error from each simulation in Figure 2B Individual measurements of cell diameters from Figure 2C Individual measurements of *α* from Figure 2D Individual mean and standard error of the mean of diffusion coefficient values from Figure 2E and 2F

In apparent agreement with these simulation results, when *E. coli* cell width was increased by treatment with the inhibitor of bacterial cell wall biosynthesis A22 (Ouzounov et al. 2016) (Figure 2C), in addition to the standard cephalexin-induced elongation, the anomalous diffusion exponent of sfGFP (Figure 2D) also significantly increased. A small, but significant increase in protein mobility is also observed (Figure 2 – figure supplement 1). Since it was previously reported that treatment with A22 can reduce dry-mass density of *E. coli* cells (Oldewurtel Enno, Kitahara, and van Teeffelen 2021), we further performed a cell sedimentation assay (Figure 2 – figure supplement 2A-C). While the treatment with cephalexin slightly (by <0.1%) decreased the density of *E. coli* cells in this assay, the additional treatment with A22, in our growth conditions, had only minor and not significant impact once the effect of the volume increase on sedimentation was accounted for (Figure 2 – figure supplement 2H). We thus conclude that the influence of A22 on the anomaly of protein diffusion is most likely due to its effect on cell width and not on the cytoplasmic density.

To additionally test our conclusion that the reduced value of *α* is due to confinement by the cell width, we performed FCS measurements for sfGFP and AcnA-sfGFP on a smaller confocal volume, thus limiting the analysis to fluorophores diffusing at a distance from the cell boundary, by reducing the pinhole size to a less optimal but smaller value of 0.66 Airy units. Indeed, the value of *α* derived from these measurements was significantly higher for both proteins, >0.9 (Figure 2 - figure supplement 3A), consistent with our prediction. As expected, the residence time (*τ_D_*) of proteins in such a smaller confocal volume is also smaller (Figure 2 - figure supplement 3B)

We next derived an Ornstein-Uhlenbeck (OU) model for fitting FCS data, where the confinement of Brownian diffusing fluorescent particles within the width of the cell is approximated by trapping in a harmonic potential of the same width (Appendix 2). The two models fit the ACF of the Brownian dynamic simulations comparably well and better than the model of unconfined Brownian diffusion (Figure 2A and Figure 2 - figure supplement 4), with the OU model having one less free parameter than the anomalous diffusion model. The OU model directly estimates the ansatz diffusion coefficient with ± 5% accuracy for the typical cell widths observed in our experiments (Figure 2 – figure supplement 5).

Since the OU model proved accurate in fitting the experimental data, comparably to the anomalous diffusion model (Figure 2 – figure supplement 6), we used it to re-fit the ACF data for all faster diffusing constructs and to estimate their Brownian diffusion coefficients (Figure 2E and Figure 1 – supplementary table 1). The dependence of *D* on molecular mass for this set of constructs was scaling as (MM)^−*β*^ with *β* = 0.56 ± 0.05 (Figure 2F, solid blue line), less steep compared to the previous estimates (Kumar, Mommer, and Sourjik 2010; Mika et al. 2010; Stracy et al. 2021) but still steeper than expected from the Stokes-Einstein relation, even when assuming *β* = 0.4 for not perfectly globular proteins (Figure 2F, black dashed line) (Enright and Leitner 2005; Smilgies and Folta-Stogniew 2015). In order to elucidate whether part of this residual deviation may be accounted for by the specific shape of fusion constructs, where sfGFP is fused to the differently-sized target proteins by a short flexible linker, we further applied a previously derived model describing diffusion of such linked proteins (Appendix 3) (Agudo-Canalejo and Golestanian 2020). The dependence of *D* on molecular mass predicted by this linked-protein model seems indeed to better recapitulate our experimental data, particularly for smaller protein fusions (Figure 2F, solid yellow line), although it moderately overestimates *D* for several of the largest protein fusions (>100 kDa). Thus, we conclude that the size dependence of diffusion for the majority of cytoplasmic proteins follows the Stokes-Einstein relation, once the shape of the sfGFP-tagged protein constructs is taken into account.

### Protein diffusion coefficients measured using FRAP or FCS are consistent

Since many previous measurements of protein diffusion in bacteria were performed using FRAP, we aimed to directly compare the results of FRAP and FCS measurements for a set of constructs of different mass. Importantly, we used the same growth conditions and microscopy sample preparation protocols as for the FCS experiments. The cells were photobleached in a region close to the pole, similar to the position that was used for the FCS experiment. The recovery of fluorescence was then followed for 11 seconds with the time resolution of 18 ms (Figure 3A). The diffusion coefficients were computed from the time course of recovery with the plugin for ImageJ, simFRAP (Blumenthal et al. 2015), which utilizes a simulation-based approach (Figure 3B). We observed very good correlation between both values of diffusion coefficients, although for most constructs the diffusion coefficients determined by FRAP were 5 to 30% lower than those obtained from the FCS data (Figure 3C and Figure 1 – supplementary table 1).

**Figure 3.**
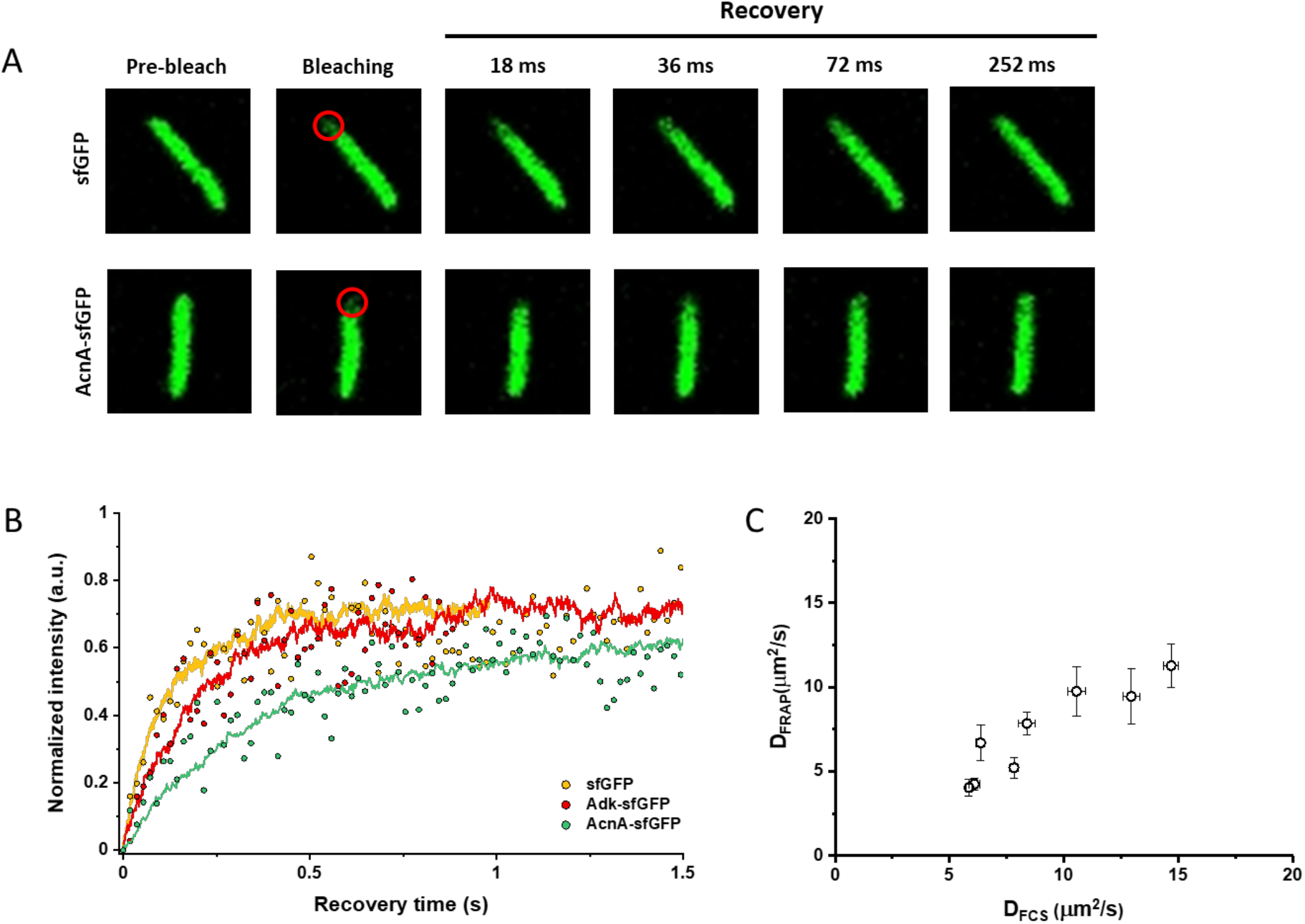
Comparison between protein diffusion coefficients measured by FCS and FRAP. (A) Examples of FRAP measurements for two different constructs, sfGFP and AcnA-sfGFP. A 3×3 pixels area close to one cell pole (red circle) was photobleached with a high intensity laser illumination for 48 ms and the recovery of fluorescence in the bleached area was monitored for 11 s with the time resolution of 18 ms. (B) Representative curves of fluorescence recovery in FRAP experiments and their fitting using simFRAP. The experimental data (colored dots) are used by the simFRAP algorithm to simulate the underlying diffusional process (colored lines). The simulation is then used to compute the diffusion coefficient. The simulation proceeds until the recovery curve reaches a plateau, therefore it is interrupted at a different time for each curve. (C) Correlation between the diffusion coefficients measured in FCS experiments (*D_FCS_*, fitting with the OU model; data from Figure 2E) and in FRAP experiment (*D_FRAP_*, fitting with simFRAP). Error bars represent the standard error of the mean. Error bars that are not visible are smaller than the symbol size. **Figure 3 – source data** Individual mean and standard error of the mean of diffusion coefficient values from Figure 3C

### Diffusive properties of cytoplasmic proteins are largely conserved between bacterial species

We then investigated whether sfGFP fusions to non-native proteins, originating from other bacteria, may show different diffusive properties in *E. coli* cytoplasm than their native counterparts. The existence of an organism-dependent “quinary” code of unspecific, short living interactions has been recently proposed in order to explain the reduced mobility of heterologous human proteins in *E. coli* cytoplasm (Mu et al. 2017). Thus, we investigated the mobility of proteins from other Gram-negative proteobacteria *Yersinia enterocolitica*, *Vibrio cholerae*, *Caulobacter crescentus* and *Myxococcus xanthus* and from the Gram-positive bacterium *Bacillus subtilis* that are homologous to several analyzed freely-diffusing *E. coli* protein constructs. Within this set of constructs, we observed no significant differences of their 1/*τ_D_* values from *E. coli* homologues. An exception was AcnA from *M. xanthus* (Figure 4A and Figure 4 – figure supplement 1A), whose lower mobility might be a sign of its multimerization, although cellular distribution of this construct was uniform. In contrast, all constructs showed slightly but mostly significantly increased anomaly of diffusion compared to *E. coli* proteins (Figure 4B and Figure 4 – figure supplement 4B), which might reflect the weakly increased propensity of non-native proteins to engage in unspecific interactions in *E. coli* cytoplasm.

**Figure 4.**
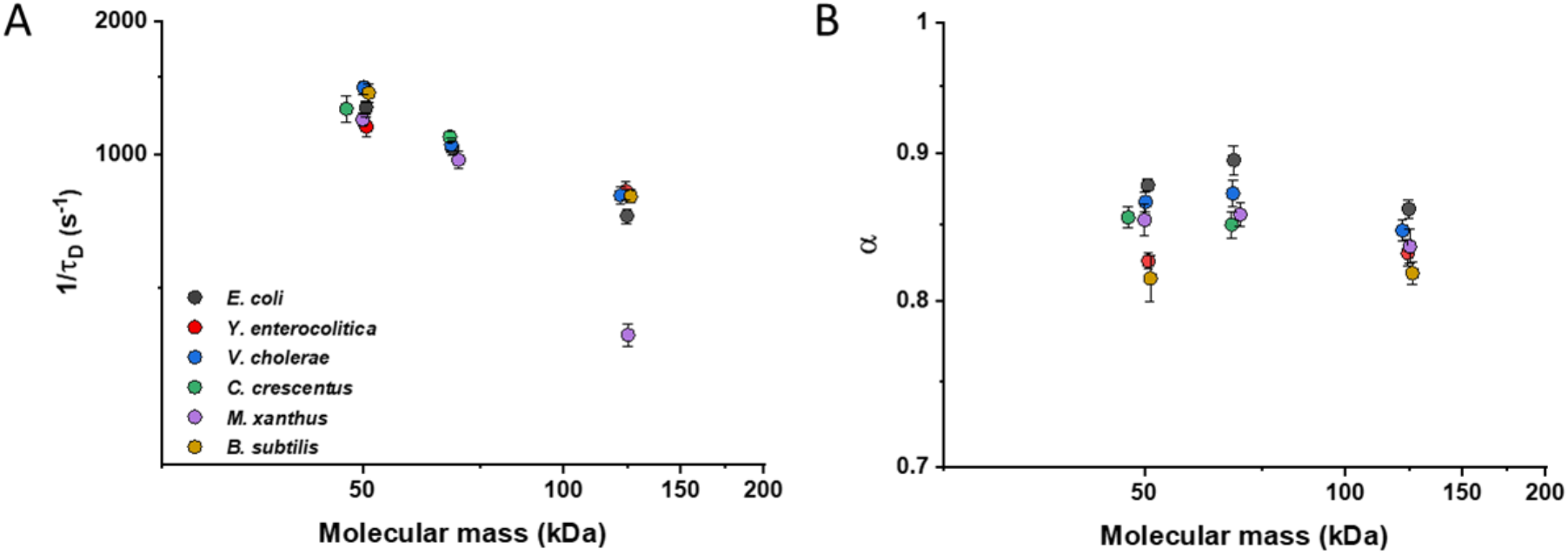
Mobility of homologous proteins from other bacterial species in *E. coli*. Mass dependence of protein mobility (1/*τ_D_*; A) and anomaly of diffusion (*α*; B) of sfGFP fusions to homologues of Adk, Pgk and AcnA from indicated bacterial species (*E.c*. = *Escherichia coli; Y.e*. = *Yersinia enterocolitica; V.c*. = *Vibrio cholerae; C.c*. = *Caulobacter crescentus; M.x*. = *Myxococcus xanthus; B.s*. = *Bacillus subtilis*) compared with that of their counterpart from *E. coli*. Each symbol represents the average value for all individual cells that have been measured for that construct (Appendix 5) and the error bars represent the standard error of the mean. Error bars that are not visible are smaller than the symbol size. **Figure 4 – source data** Individual mean and standard error of the mean of 1/*τ_D_* values from Figure 4A Individual mean and standard error of the mean of *α* values from Figure 4B

### Effects of osmolarity, temperature, antibiotics and cell growth on mobility of differently-sized proteins

We further characterized the impact of several environmental and cellular perturbations of the bacterial cytoplasm on protein mobility, using apparently freely diffusing protein fusions of different sizes as probes. We started with the previously characterized decrease in mobility of GFP and large protein complexes or aggregates upon osmotic upshift (Konopka et al. 2006; Konopka et al. 2009; Mika et al. 2010; Liu et al. 2019; Wlodarski et al. 2020). *E. coli* cells exposed to increased ionic strength by addition of 100 mM NaCl showed decrease in cell length and width (Figure 5 – figure supplement 1A, B) and an increase in cell density in the sedimentation assay (Figure 2 – figure supplement 2D), consistent with a previous report (Wlodarski et al. 2020). Higher ionic strength also significantly decreased the mobility of sfGFP (Figure 5A and Figure 5 – figure supplement 2A), comparably to previously measured values (Konopka et al. 2009; Mika et al. 2010). Importantly, the mobility of all other tested constructs decreased proportionally (Figure 5A), meaning that – in this range of molecular sizes – the effect of a moderate osmotic upshift can be interpreted as a simple increase in cytoplasmic viscosity due to higher molecular crowding, which is in contrast to the different effects of high osmolarity on small molecules and on GFP (Mika et al. 2010). No effect is observed on the anomaly of diffusion for any protein construct (Figure 5 – figure supplement 3A).

**Figure 5.**
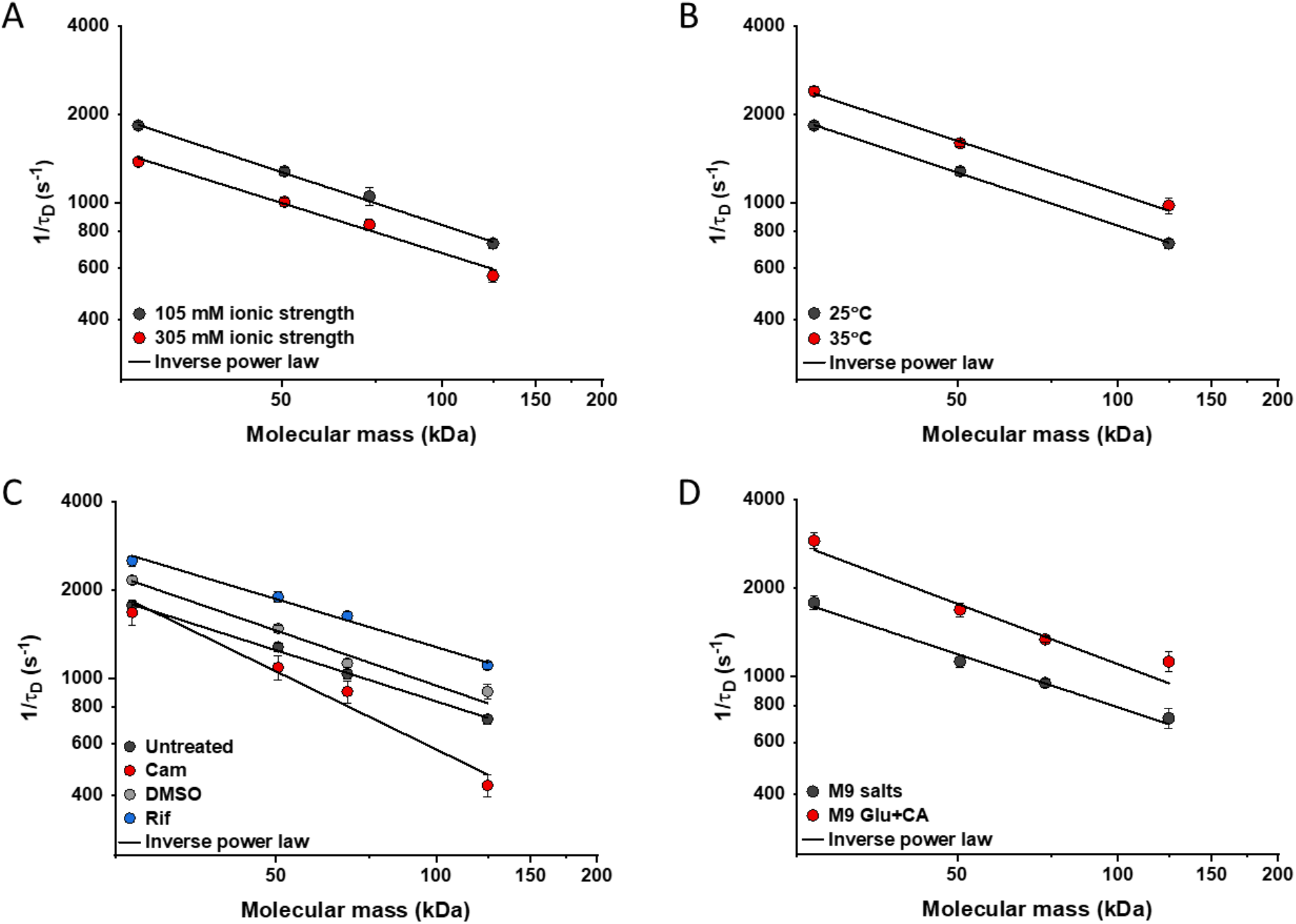
Effects of physicochemical perturbations and cell growth on mobility of differently-sized proteins. Each dot represents the average value of protein mobility (1/*τ_D_*) of all the cells measured for the construct of the indicated molecular mass (Appendix 5). Error bars represent the standard error. Error bars that are not visible are smaller than the symbol size. The solid black lines are the fit with an inverse power law to extract the size dependence of protein mobility (*β*) in that condition. (A) Protein mobility measured in cells that were resuspended in either tethering buffer (ionic strength of 105 mM; *β* = 0.60 ± 0.01) or in the same buffer but supplemented with additional 100 mM NaCl (total ionic strength of 305 mM; *β* = 0.57 ± 0.05). The measurements were performed in agarose pads prepared at the same ionic strength. (B) Protein mobility at different environmental temperatures. As for the other experiments, *E. coli* cultures were grown at 37°C and bacterial cells during the measurements were incubated at 25°C (*β* = 0.60 ± 0.01) or at 35°C (*β* = 0.60 ± 0.05), as indicated. (C) Protein mobility in control cells (*β* = 0.58 ± 0.02) and after treatment with chloramphenicol (Cam; 200 μg/mL; *β* = 0.88 ± 0.11), rifampicin (Rif; 200 μg/mL, in 0.1% v/v DMSO; *β* = 0.54 ± 0.04), or DMSO control (0.1% v/v; *β* = 0.62 ± 0.07) as indicated. Antibiotics were added to growing *E. coli* culture 60 minutes prior to harvesting. (D) Protein mobility in non-growing cells incubated at 35°C on agarose pads containing only M9 salts (*β* = 0.60 ± 0.05) in comparison with growing cell incubated on pads with M9 salts supplemented with 20 mM glucose and 0.2% casamino acids (Glu + CA; *β* = 0.68 ± 0.10). **Figure 5 – source data** Individual mean and standard error of the mean of 1/*τ_D_* values from Figure 5A Individual mean and standard error of the mean of 1/*τ_D_* values from Figure 5B Individual mean and standard error of the mean of 1/*τ_D_* values from Figure 5C Individual mean and standard error of the mean of 1/*τ_D_* values from Figure 5D

Next, we studied the effect of environmental temperature on protein mobility in bacterial cytoplasm. According to the Stokes-Einstein equation, the diffusion of a particle directly depends on the system’s temperature in Kelvin and on the viscosity of the fluid, which itself changes with temperature. In the biologically relevant range, the temperature sensitivity of diffusion is primarily determined by the temperature dependence of water viscosity. The measured increase in mobility of sfGFP and two other constructs, by approximately 20-25% between 25°C and 35°C (Figure 5B and Figure 5 – figure supplement 2B), agrees well with the temperature-dependent decrease in water viscosity over 10°C (Huber et al. 2009). Expectedly, the effect of imaging temperature was not linked to any changes of the cell size (Figure 5 – figure supplement 1C, D). Of note, a weak, but consistent, increase in the anomaly of protein diffusion is also observed at higher environmental temperature (Figure 5 – figure supplement 3B). Instead, the growth temperature of the *E. coli* culture had no apparent effect on protein mobility (Figure 5 – figure supplement 4), suggesting that – at least in the tested temperature range – *E. coli* lacks the growth-temperature dependent regulation of cytoplasmic viscosity that has been recently reported in the budding yeast (Persson, Ambati, and Brandman 2020).

Antibiotics that inhibit transcription (e.g., rifampicin) or translation (e.g., chloramphenicol) are known to affect the spatial organization of bacterial chromosomes (Bakshi et al. 2014). The mobility of chromosomal loci and of large cytoplasmic aggregates was also shown to be affected by several antibiotics, in apparent correlation with changes in the cytoplasmic density (Wlodarski et al. 2020). We observed that chloramphenicol treatment caused a minor increase in cell width (Figure 5 – figure supplement 1F) and a decrease in cell density (Figure 2 – figure supplement 2G). However, protein mobility rather decreased in chloramphenicol-treated cells, opposite to what could be expected based alone on the chloramphenicol-induced reduction of cell density (Figure 5C and Figure 5 – figure supplement 2C). The reduced protein mobility could neither be simply explained by compaction of the nucleoid in cells treated with chloramphenicol, since it was only marginally lower inside than outside of the nucleoid (Figure 5 – figure supplement 5A, B). It should be noted that no significant difference in the anomaly of diffusion (Figure 5 – figure supplement 5C) was observed inside or outside of the nucleoid.

In contrast, inhibition of RNA transcription by rifampicin treatment led to a marked increase in protein mobility (Figure 5C and Figure 5 – figure supplement 2C). Such higher protein mobility is consistent with the previously observed rifampicin-induced reduction of macromolecular crowding in bacterial cytoplasm (Wlodarski et al. 2020), although only a minor decrease in cell density was observed in our sedimentation assay (Figure 2 – figure supplement 2F) beyond the effect of DMSO that was used as a solvent for rifampicin (Figure 2 – figure supplement 2E). Similar to the effects of osmolarity and temperature, the increase in protein mobility caused by the rifampicin treatment, and its decrease induced by chloramphenicol were similar for all tested proteins (Figure 5C), except for the AcnA-sfGFP construct that was disproportionally affected by chloramphenicol in both mobility and anomaly of diffusion (Figure 5 – figure supplement 3C).

Finally, we investigated whether protein mobility might be influenced by cell growth, comparing FCS measurements in cells incubated at 35°C on agarose pads containing either only M9 salts or M9 salts plus glucose and casamino acids. These conditions had only minor impact on the cell shape (Figure 5 – figure supplement 1G, H). Although at this high temperature residual growth was also observed for cells on M9 salt pads, cell growth in presence of nutrients was expectedly much more pronounced. The observed protein mobility was also much higher in the presence of nutrients, and this increase was again similar for the four tested differently-sized constructs (Figure 5D and Figure 5 – figure supplement 2D), while no consistent trend was observed in the anomaly of protein diffusion across these conditions (Figure 5 – figure supplement 3D). To further distinguish the respective contributions of metabolic activity and of biosynthesis and resulting cell growth, we incubated cells in presence of both nutrients and chloramphenicol on the agarose pad. Similar to our previous experiments where chloramphenicol was added to the batch culture, its addition had no or little effect on the mobility of sfGFP or the AcnA-sfGFP construct in absence of nutrients (Figure 5 – figure supplement 6). In contrast, protein mobility in presence of nutrients was strongly affected by chloramphenicol treatment. Thus, the enhanced protein mobility in presence of nutrients appears to be primarily due to active protein production and cell growth. Nevertheless, even chloramphenicol-treated cells exhibited a moderate increase in protein mobility in presence of nutrients, indicating that the metabolic activity contributes to the overall effect of growth on diffusion. It is possible that the contribution of the metabolic activity might be even larger than it appears in this experiment, since the inhibition of protein translation might reduce metabolic activity. In any case, the impact of growth on diffusion of individual proteins cannot be simply explained by the energy state of the cell, since lowering it by the inhibition of respiration-dependent ATP synthesis using treatment with dinitrophenol (DNP) did not reduce protein mobility, at either 25°C or 35°C. This is contrary to the effect of the DNP treatment on large cytoplasmic particles (Parry et al. 2014) (Figure 5– figure supplement 7). An interesting exception was the mobility of Adk-sfGFP, which was indeed reduced by the DNP treatment at high temperature. This, however, might be a specific effect related to the enzymatic activity or conformation of Adk that binds ATP as a substrate.

## Discussion

Bacteria rely on translational diffusion to deliver proteins and other macromolecules to their cellular destinations, including their reaction partners, and the diffusional properties of bacterial cytoplasm are therefore fundamental to the understanding of bacterial cell biology. Consequently, a number of studies have investigated protein mobility in bacteria, all showing strong effects of macromolecular crowding in the bacterial cytoplasm on diffusion (Konopka et al. 2006; Mullineaux et al. 2006; Konopka et al. 2009; Kumar, Mommer, and Sourjik 2010; Mika et al. 2010; Nenninger, Mastroianni, and Mullineaux 2010). Nevertheless, the relatively small number of proteins investigated in each of these previous studies, and the differences between strains, growth conditions and between methodologies, limited general conclusions about protein mobility, even in the most-studied environment of *E. coli* cytoplasm. For example, combining data from different studies to determine the relation between the size of a protein and its cytoplasmic diffusion coefficient yielded only uncertain estimates (Mika and Poolman 2011; Schavemaker, Boersma, and Poolman 2018). Such variability between different studies might be further compounded by potentially profound effects on diffusion of size-independent protein properties such as surface charge (Schavemaker, Smigiel, and Poolman 2017) or weak interactions with other proteins and other cellular components (von Bülow et al. 2019). Similarly, it remains unclear whether a typical protein in the cytoplasm exhibits Brownian diffusion, as has been shown in few examples (Bakshi, Bratton, and Weisshaar 2011; English et al. 2011), or rather a subdiffusive behavior as common in eukaryotic cells (Di Rienzo et al. 2014; Sabri et al. 2020) and for large proteins and nucleoprotein particles in bacteria (Golding and Cox 2004; Lampo et al. 2017; Yu et al. 2018). Additionally, while in eukaryotic cells anomalous diffusion is primarily associated with hindrance by intracellular structures, the possible causes of anomalous diffusion in bacteria are still unclear.

Here, we addressed these questions by systematically investigating the diffusive behavior of a large set of fluorescent protein fusions to differently-sized cytoplasmic proteins of *E. coli*. We demonstrate that the majority of studied proteins exhibit a rather uniform relation between their molecular mass and cytoplasmic mobility, with a clear upper bound on protein mobility at a given molecular mass. This bound likely reflects the fundamental size-specific physical limit on protein diffusion in *E. coli* cytoplasm, with lower mobility of individual proteins being due to their interactions with other cellular components.

Furthermore, our simulations suggest that the apparent weak anomaly of diffusion observed in the FCS data analysis, with *α* ~ 0.85-0.9 for most proteins, could be accounted for by confinement of the otherwise purely Brownian diffusing particles in the small volume of a bacterial cell. Similar conclusions have been drawn by previous SPT studies for several individual proteins (Bakshi, Bratton, and Weisshaar 2011; English et al. 2011). This interpretation is consistent with our measurements of diffusion in A22-treated *E. coli* with an increased cell. Although the interpretation of these experiments might be complicated by the reduced cytoplasmic density of A22-treated bacteria (Oldewurtel Enno, Kitahara, and van Teeffelen 2021), under our conditions the effect of A22 on cell density seems to be negligible. It is also supported by the FCS measurements using smaller confocal volume, which yielded higher value of *α*, as could be expected from protein diffusion away from the cell boundary.

We therefore used a model of purely Brownian diffusion under confinement (Ornstein-Uhlenbeck model) to determine diffusion coefficients by directly fitting the autocorrelation functions of our FCS measurements. The obtained overall dependence of diffusion coefficients on the molecular mass of the fusion protein showed the exponent *β* = 0.56, steeper than predicted by the Stokes-Einstein relation, with *β* = 0.33 for fully compact proteins or *β* = 0.4 for the more realistic case where proteins are assumed to be not entirely compact (Enright and Leitner 2005; Smilgies and Folta-Stogniew 2015). Nevertheless, at least for smaller constructs, the observed dependence of the diffusion coefficient on the molecular mass could be well reproduced once the specific shape of fusion constructs, where two roughly globular proteins are fused by a short linker, was taken into account along with their imperfect globularity (Agudo-Canalejo and Golestanian 2020). Only largest proteins in our set (above 100 kDa) showed mobility that was slower than predicted by this model, possibly because diffusion of larger proteins is more strongly impacted by weak interactions with other macromolecules (von Bülow et al. 2019).

Our analysis thus suggests that, despite the high crowdedness of the bacterial cytoplasm, the diffusion of typical cytoplasmic proteins in bacteria is mostly Brownian and can be well described by treating the cytoplasm as a viscous fluid, with only a moderate dependence of the effective viscosity on the size of diffusing proteins. Given the diffusion coefficient determined in our study for free sfGFP, ~ 14 μm^2^s^−1^, for small proteins this effective viscosity of bacterial cytoplasm is only approximately six times higher than in dilute solution (Potma et al. 2001). This diffusion coefficient for GFP is substantially larger than the values reported in the early studies that used FRAP (Elowitz Michael et al. 1999; Konopka et al. 2006; Mullineaux et al. 2006; Konopka et al. 2009; Kumar, Mommer, and Sourjik 2010; Mika et al. 2010; Nenninger, Mastroianni, and Mullineaux 2010), although it is consistent with other FCS and SPT studies (Meacci et al. 2006; English et al. 2011; Sanamrad et al. 2014; Diepold et al. 2017; Rocha et al. 2019). These differences are apparently due to the limitations of early FRAP analyses that generally underestimated protein mobility, rather than due to different spatial and temporal scales assessed by the two techniques, since our direct comparison between FCS and FRAP measurements yielded similar values of diffusion coefficients. Indeed, a more recent FRAP study also reported higher diffusion coefficients for GFP (Schavemaker, Smigiel, and Poolman 2017).

Several proteins in our set showed much lower mobility than expected from their size, and in some cases also clearly subdiffusive behavior. For three selected examples, this deviation could be explained by specific association with other proteins or multiprotein complexes, since disrupting these interactions both increased protein mobility and reduced subdiffusion. This is consistent with theoretical studies suggesting that binding of diffusing molecules to crowders can lead to subdiffusion (Saxton 2007; Guigas and Weiss 2008). Thus, protein-protein interactions may be the main cause of protein subdiffusion in bacterial cytoplasm, although other explanations might hold for subdiffusion of large cytoplasmic particles (Golding and Cox 2004; Lampo et al. 2017; Yu et al. 2018).

Unspecific transient interactions might also explain the slightly subdiffusive behavior of sfGFP fusions to proteins from other bacteria in *E. coli* cytoplasm. However, this anomaly was weak and there was overall only little difference between the mobility of these non-native proteins and their similarly-sized *E. coli* homologues, which is in contrast to pronounced differences observed between bacterial and mammalian proteins (Mu et al. 2017). Thus, there is apparently little organism-specific adaptation of freely diffusing proteins to their “bacterial host”, with a possible exception of bacteria with extreme pH or ionic strength of the cytoplasm (Schavemaker, Smigiel, and Poolman 2017). This might facilitate horizontal gene transfer among bacteria, by ensuring that their surface properties do not hinder accommodation of proteins in a new host.

We further probed how the effective viscous properties of bacterial cytoplasm changed under different physicochemical perturbations, using a subset of proteins that showed fastest diffusion for their molecular mass as reporters of unhindered diffusion. Consistent with the importance of macromolecular crowding and in agreement with previous results (Konopka et al. 2009), protein mobility decreased upon osmotic upshift as cytoplasmic crowding increases. In contrast, the effective cytoplasmic viscosity decreases significantly (~20%) upon treatment with rifampicin that inhibits transcription and thereby reduces the overall macromolecular crowding. This observation is consistent with recent SPT measurements on large cytoplasmic particles (Wlodarski et al. 2020; Rotter et al. 2021), and it agrees well with the relative contribution of RNA to the macromolecular composition of an *E. coli* cell (Cayley et al. 1991) and with the reduction of molecular crowding in rifampicin-treated cells (Wlodarski et al. 2020).

Despite multiple effects of environmental temperature on cellular processes, such as the active (nonthermal) stirring of the cytoplasm at higher temperature (Weber, Spakowitz, and Theriot 2012), the temperature dependence of the cytoplasmic viscosity in the tested range was similar to that of water and consistent with the Stokes-Einstein relation, decreasing by 20-30% for a temperature increase of 10°C (Huber et al. 2009). Furthermore, the same temperature dependence of protein mobility was observed upon treatment with the protonophore DNP that deenergizes cells by dissipating proton gradient, arguing against general active stirring of cytoplasm in *E. coli* under our experimental conditions. We further observed no dependence of the effective cytoplasmic viscosity on growth temperature, in contrast to the homeostatic adaptation of bacterial membrane fluidity (Sinensky 1974) and of bacterial signaling (Oleksiuk et al. 2011; Almblad et al. 2021) to the growth temperature. Since growth-temperature dependent adaptation of the cytosolic viscosity was recently reported for budding yeast (Persson, Ambati, and Brandman 2020), it is surprising that such compensation apparently does not exist in *E. coli*. One possible explanation for this difference might be a broader range of growth temperatures for budding yeast *Saccharomyces cerevisiae* compared to *E. coli*, and a stronger temperature effect on protein diffusion in the yeast cytosol. Of note, here we did not explore protein diffusion in thermally stressed *E. coli* cells, which might have more profound effects on the properties of bacterial cytoplasm as recently shown for *Listeria monocytogenes* (Tran et al. 2021).

Finally, we observed that protein mobility was significantly higher in rapidly growing cells. This “fluidizing” effect of growth seems to be primarily due to the biosynthetic processes, likely protein translation, as evidenced by the reduced mobility upon chloramphenicol treatment, or to cell growth itself. The contribution of metabolic activity in presence of nutrients was also significant but weaker, although it might be underestimated since inhibition of protein biosynthesis by chloramphenicol could possibly indirectly reduce metabolic activity. Thus, the observed phenomenon may be different from previously characterized ATP-dependent fluidization of the bacterial cytoplasm that enables mobility of large multiprotein complexes but apparently does not affect free GFP (Montero Llopis et al. 2012; Parry et al. 2014), as also observed for sfGFP and other constructs in our experiments. The interplay between these energy-, metabolism- and growth-dependent effects on diffusional properties of bacterial cytoplasm remains to be investigated.

Importantly, we observed that these perturbations to the cytoplasmic protein mobility, including cell growth and changes to the macromolecular crowding and temperature, have proportional effects on differently-sized proteins. These results suggest that - within the tested size range - protein diffusion in *E. coli* cytoplasm remains Brownian under all tested conditions, including growing cells, and effects of these perturbations on protein mobility can be simply accounted for by changes in the cytoplasmic viscosity. We hypothesize that such proportional changes in diffusion of differently-sized proteins might be important to maintain balanced rates of diffusion-limited cellular processes under various environmental conditions.

## Materials and methods

### Bacterial strains, plasmids and media

All strains and plasmids used in this study are listed in Figure 1 – supplementary table 2. All experiments were performed in the *Escherichia coli* strain W3110 (Serra et al. 2013). Genes of interest were amplified by PCR using Q5 polymerase (New England Biosciences) and cloned in frame with sfGFP using Gibson assembly (Gibson et al. 2009) into pTrc99A vector (Amann, Ochs, and Abel 1988), under control of the *trc* promoter inducible by isopropyl ß-D-1-thiogalactopyranoside (IPTG). In all cases sfGFP was fused at the C-terminus of the protein of interest with a GGGGS linker. The stability of the fusion constructs was verified by gel electrophoresis and immunoblotting using an anti-GFP primary antibody (JL-8 monoclonal, Takara). Point mutations were introduced by site-directed mutagenesis (New England Biosciences). The *ΔclpA* strain was generated by transferring the kanamycin resistance cassette from the corresponding mutant in the Keio collection (Baba et al. 2006) by P1 transduction. The cassette was further removed by FLP recombinase carried on the temperature-sensitive plasmid pCP20 (Cherepanov and Wackernagel 1995).

*E. coli* cultures were grown in M9 minimal medium (48 mM Na_2_HPO_4_, 22 mM KH_2_PO_4_, 8.4 mM NaCl, 18.6 mM NH_4_Cl, 2 mM MgSO_4_, 0.1 mM CaCl_2_) supplemented with 0.2% casamino acids, 20 mM glucose and 100 μg/mL ampicillin for selection. The overnight cultures were diluted to OD_600_ = 0.035 and grown for 3.5 hours at 37°C and 200 rpm shaking. Cultures were treated for additional 45 minutes, under the same temperature and shaking conditions, with 100 μg/mL cephalexin and with 0 - 15 μM IPTG (Figure 1 – supplementary table 1), to induce expression of the fluorescent protein constructs. Where indicated, cultures were further incubated with 200 μg/mL rifampicin, DMSO as a mock treatment, 200 μg/mL chloramphenicol or 2 mM DNP for 1 hour or with 1 μg/mL A22 for 4 hours under the same temperature and shaking conditions.

### Growth curves

Measurements of bacterial growth were performed using 96-well plates (Cellstar transparent flat-bottom, Greiner). Overnight cultures were inoculated at an initial OD_600_ of 0.01 in the same medium as used for growth in other experiments. Each well contained 150 μl of culture and the plate was covered with the plastic cover provided by the producer and further sealed with parafilm that prevents evaporation but allows air exchange. Plates were incubated at 37°C with continuous shaking, alternating between 150 s orbital and 150 s linear, in a Tecan Infinite^®^ 200 PRO plate reader.

### FCS data acquisition

Cells were harvested by centrifugation at 7000g for 3 minutes and washed 3 times in tethering buffer (10 mM K_2_HPO_4_, 10 mM KH_2_PO_4_, 1 μM methionine, 10 mM sodium lactate, buffered with NaOH to pH 7). When indicated, 1 mL of chloramphenicol-treated cells were stained for 15 minutes with 300 nM SYTOX Orange Nucleic Acid Stain (Invitrogen). The excess of SYTOX Orange was washed in tethering buffer before proceeding with FCS experiments. 2.5 μL of bacterial cells were then spread on a small 1% agarose pad prepared in tethering buffer salts (10 mM K_2_HPO_4_, 10 mM KH_2_PO_4_ buffered with NaOH to pH 7), unless differently stated. Imaging was performed on Ibidi 2-well μ-Slides (#1.5H, 170 μm ± 5 μm). After the 45 minutes treatment with cephalexin, length of most bacterial cells was in a range of 4-8 μm.

FCS measurements were performed on a LSM 880 confocal laser scanning microscope (Carl Zeiss Microscopy) using a C-Apochromat 40×/1.2 water immersion objective selected for FCS. sfGFP was excited with a 488 nm Argon laser (25 mW) and fluorescence emission was collected from 490 to 580 nm. SYTOX Orange was excited with a 543 nm laser and fluorescence emission was collected from 553 to 615 nm. In order to avoid partial spectral overlap between the emission spectra of sfGFP and SYTOX Orange, fluorescence emission of sfGFP in the co-staining experiments was collected from 490 to 535 nm. Each sample was equilibrated for at least 20 minutes at 25°C (or 35°C when specified), on the stage of the microscope and measurements were taken at the same temperature. FCS measurements were acquired within 60 minutes from the sample preparation. The pinhole was aligned on a daily basis, by maximizing the fluorescence intensity count rate of an Alexa488 (Invitrogen) solution (35 nM) in phosphate-buffered saline (PBS; 137 mM NaCl, 2.7 mM KCl, 8 mM Na_2_HPO_4_, 1.8 mM KH_2_PO_4_, pH 7.4). Unless differently stated, all measurements were performed with a pinhole size correspondent to 1 Airy unit, to ensure the optimal gathering of fluorescence signal. The coverslip collar adjustment ring of the water immersion objective was also adjusted daily, maximizing the fluorescence intensity signal and the brightness of Alexa 488. The laser power was adjusted in order to obtain molecular brightness (i.e. photon counts per second per molecule, cpsm) of 10 kcpsm for Alexa 488, using the ZEN software (Carl Zeiss Microscopy). The brightness of Alexa 488 was used as a daily reference to ensure constant laser power and adjusting it using the software-provided laser power percentage whenever necessary (range over the entire set of measurements was 0.11 – 0.18%). Before each measurement session, we acquired three sequential FCS measurements of Alexa488 in PBS, to verify the reproducibility of the confocal volume shape and size. The ratio between axial and lateral beam waist 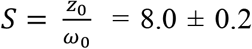 (Avg. ± SEM) was obtained from a Brownian fit of the Alexa 488 autocorrelation curves using the ZEN software. For the lateral beam waist, we obtained *ω_0_* = 0.186 ± 0.001 μm (Avg. ± SEM), calculated from the diffusion time *τ_D_* = 20.9 ± 0.11 μs (Avg. ± SEM) obtained from the Brownian fit, being

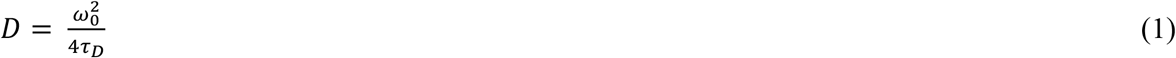

and being *D_Alexa488_* = 414 μm^2^/s at 25°C (Petrov et al. 2006).

For the FCS measurements *in vivo*, the laser was positioned at the center of the short length axis and typically 0.8-1 μm from one of the cell poles along the long axis. For each cell, six sequential fluorescence intensity acquisitions of 20 seconds each were performed on the same spot (Figure 1 – figure supplement 3). The laser power used for measurements *in vivo* was fixed to a value about 7 times lower than for Alexa488 in PBS, in order to reduce photobleaching. Confocal images of the selected cell were routinely acquired before and after the FCS measurement to verify focal (*z*) and positioning (*xy*) stability (see Appendix 1 for additional information on the FCS measurements).

### FCS data analysis

Due to the small size of bacterial cells, fluorescence intensity traces are affected by photobleaching (Appendix 1). The effect of photobleaching on autocorrelation curves was corrected by detrending the long-time fluorescence decrease of each of the six fluorescence intensity traces using an ImageJ plugin (Jay Unruh, https://research.stowers.org/imagejplugins/index.html, Stowers Institute for Medical Research, USA). The plugin calculates the autocorrelation function (ACF) from each fluorescence intensity trace, correcting it for the photobleaching effect by approximating the decreasing fluorescence intensity trend with a multi-segment line (the number of segments was fixed to two). We obtained almost identical ACFs correcting for photobleaching effects by local averaging (Appendix 1 - figure 3) using the FCS-dedicated software package Fluctuation Analyzer (Wachsmuth et al. 2015). In both cases, ACFs were calculated starting at 2 μs, since at times shorter than 2 μsec, ACFs can be significantly affected by the GaAsp photomultipliers afterpulsing.

For each FCS measurement, we fitted all the six ACFs, calculated using the multi-segment detrending method, with a three-dimensional anomalous diffusion model that includes one diffusive component and one blinking component due to the protonation-deprotonation of the chromophore of sfGFP, according to the Equation (2):

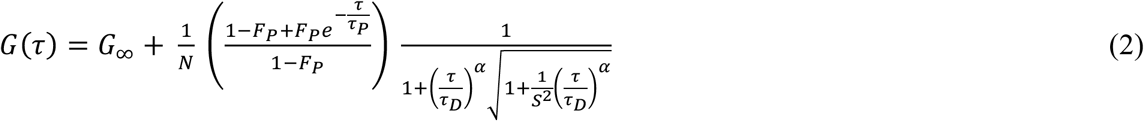

where *N* is the average number of particles in the confocal volume, *F_P_* is the fraction of particles in the non-fluorescent state, *τ_P_* is the protonation-deprotonation lifetime at pH 7.5, 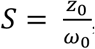, the aspect ratio of the confocal volume with *z_0_* and *ω_0_* being the axial and lateral beam waists, *τ_D_* is the diffusion time in the confocal volume, *α* is the anomalous diffusion exponent, and *G_∞_* is the offset of the autocorrelation function. The protonation-deprotonation lifetime (*τ_P_*) for sfGFP was fixed to 25 μs according to FCS measurements for sfGFP in PBS at pH 7.5 (Cotlet et al. 2006). The aspect ratio of the confocal volume was fixed to *S* = 8 in the fittings to be consistent with the experimental calibration (see above). All other parameters were left free. For each FCS measurement, we calculated the average diffusion time *τ_D_* and the average anomalous diffusion exponent *α* based on the autocorrelation curves of the six sequential fluorescence intensity traces. Importantly, no significant trend in *τ_D_* or *α* was apparent when comparing the six sequential autocorrelation functions acquired for a given bacterial cell (Appendix 1 - figure 4). Fitting to the anomalous diffusion model was performed using the Levenberg-Marquardt algorithm in the FCS analysis-dedicated software QuickFit 3.0 developed by Jan Wolfgang Krieger and Jörg Langowski (Deutsches Krebsforschungszentrum, Heidelberg, https://github.com/jkriege2/QuickFit3). Identical results were obtained when fitting the data with OriginPro.

Alternatively, the autocorrelation functions were fitted by the Ornstein-Uhlenbeck model (Appendix 2) according to Equation (3):

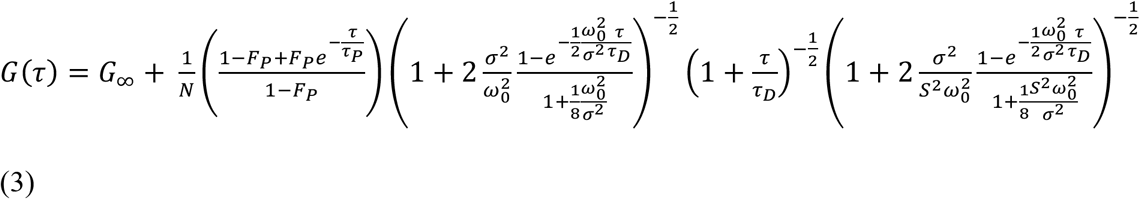

where *S* and *τ_P_* were fixed to the same values mentioned for equation (2), *ω_0_* was fixed to 0.19 and *σ* was fixed to *d*/2=0.42 μm, being *d* the typical diameter of an *E. coli* cell (see Ornstein-Uhlenbeck model validation paragraph). Fitting to the Ornstein-Uhlenbeck model was performed with OriginPro.

### FRAP data acquisition and analysis

Cells for FRAP experiments were grown and prepared for imaging following the same protocol as for the FCS measurements. Due to the higher sensitivity of FCS at low fluorophore concentrations, several fusion constructs required higher induction by IPTG (Figure 1 – supplementary table 1) to obtain fluorescence intensity suitable for FRAP. The same LSM 880 confocal microscope, including objective and light path was used for FRAP as for the FCS measurements. The bacterial cell was imaged at 40×40 pixels with 30× zoom (pixel size 0.177 μm) with a pixel dwell time of 3.15 μs. First, 15 pre-bleaching frames were acquired at 2% laser power, subsequently the photobleaching was performed on a 3×3 pixels area on one cell pole with 100% laser power for a total of 48 ms and 584 post-bleaching frames were acquired to monitor the fluorescence recovery.

We observed that the mobile fraction for all constructs was >0.9. FRAP measurements were analyzed using simFRAP (Blumenthal et al. 2015), an ImageJ plugin based on a simulation approach implemented in a fast algorithm, which bypasses the need of using analytical models to interpolate the data. The simFRAP algorithm simulates two-dimensional random walks in each pixel, using the first image acquired after bleaching to define initial and boundary conditions, and it resolves numerically the diffusion equation by iterative simulation. The frame time and pixel size were fixed respectively to 0.018 s and 0.177 μm, and the target cell and the bleached region were defined as ImageJ ROIs (regions of interest). Of note, we used the target cell itself as a reference to compensate for the gradual bleaching during the measurement, as done previously (Kumar, Mommer, and Sourjik 2010). This enabled us to achieve the highest possible temporal resolution, by reducing the acquisition area to a single *E. coli* cell. The FRAP derived diffusion coefficient *D_FRAP_* was directly obtained as output of the plugin.

### Cellular density measurements

Cell cultures were grown following the same protocol as for the FCS and FRAP measurements. Cultures were harvested at 4000g for 5 minutes, and the pellet was resuspended in motility buffer (MB) (10 mM KPO_4_, 0.1 mM EDTA, 67 mM NaCl, 0.01% Tween 80). Tween 80 is a surfactant that prevents cell-surface adhesion (Nielsen et al. 2016; Schwarz-Linek et al. 2016). Bacterial suspension was adjusted to a high cell density (OD_600_=15) by subsequent centrifugation (4000g, 5 minutes) and resuspension in a medium containing 20% iodixanol to match the density of MB with that of *E. coli* cell (1.11 g/ml) (Martínez-Salas, Martín, and Vicente 1981). Each sample was then loaded in the chamber of a previously fabricated poly-di-methylsiloxane (PDMS) microfluidic device. The chamber consists of an inlet connected to an outlet by a straight channel of 50 μm height, 1 mm width and 1 cm length. The channel was then sealed with grease to prevent fluid flows. After letting the mixtures reach the steady state in the microfluidic device for 40 minutes, cell sedimentation was visualized by acquiring Z-stack images of the whole microfluidic channel using the same microscopy setup as for the FCS and FRAP measurements (1px = 0.2 μm in X and Y, 1px = 1 μm in Z; field of view = 303.64 × 303.64 × 70 μm^3^, 0.35 μs/px exposure). The number of cells in each Z plane was quantified by the connected components labeling algorithm for ImageJ (Legland, Arganda-Carreras, and Andrey 2016). Each experiment was conducted in three technical replicates. Because the height and the tilt of the microfluidic channels slightly varies from sample to sample, the Z position was binned and the mean of the cell fraction over the bins was calculated.

The vertical density profiles were fitted to the theoretical expectation for diffusing particles in a buoyant fluid, 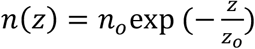, in the range *z* = [0.25, 0.8] × 50 μm to avoid effects of sample boundaries. The fitted decay length is expected to obey 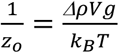, with *Δρ* the difference in density between the cells and the suspending fluid, *V* the average volume of the cells, *g* = 9.81 m^2^/s the acceleration of gravity and *k_B_T* = 4.11 pN×nm the thermal energy at 25°C. To compute the buoyancy-corrected cell density 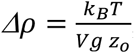, the cell volume was estimated assuming the cells are cylinders closed by hemispherical caps, *V* = *πd*^3^/6 + (*L* − *d*)*πd*^2^/4. For all conditions, the cell diameter *d* was evaluated on confocal images taken prior to FCS measurement (see Figure 1 – figure supplement 7, Figure 2C and Figure 5 – figure supplement 1), and so was the length of cephalexin treated cells (L = 5.5 ± 0.1 (SEM) μm), cephalexin + A22 treated cells (L = 5.8 ± 0.1 (SEM) μm), and untreated cells (L=2.8 ± 0.2 μm). Cell length for 100 mM NaCl, DMSO, rifampicin and chloramphenicol treated cells was kept equal to the one of untreated cells, because cephalexin was not used during culture growth for sedimentation assay for these conditions.

### Brownian dynamics simulations

We performed Brownian dynamics simulations of uncorrelated point particles under confinement. The *N* = 50 fluorescent particles performed a random walk with steps taken from a Gaussian distribution of width 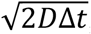, with *D* the free diffusion coefficient and Δ*t* = 10^6^ *s* the simulation step. Confinement was imposed by redrawing the random steps that moved out of the confinement volume. Imposing elastic reflections on the walls yielded identical results. The confinement volume was assumed to be a cylinder of diameter *d* and length (*L-d*) closed at both ends by hemispheric caps of diameter *d*, idealizing the shape of *E. coli*. The cell length was fixed to *L* = 5 μm. The diameter was varied in the range *d* = [0.7, 1] μm. The collected fluorescence intensity was computed at each time step assuming a Gaussian intensity profile of the laser beam, 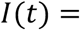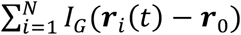 with ***r***_*i*_(*t*) the position of particle *i*, ***r***_0_ the center of the confocal volume and 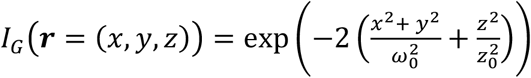, with *ω*_0_ = 200 nm and *z*_0_ = 800 nm the lateral and axial widths of the confocal volume. The normalized intensity autocorrelation 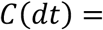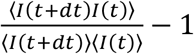 is computed for logarithmically spaced lag times *dt*, to reflect experimental practices. The center of the confocal volume was chosen in the center of the cell along the *y* and *z* axes and 1 μm away from the edge of the cell along the longitudinal *x* axis of the cell, similarly to experimental conditions. The intensity autocorrelation function was finally multiplied by an exponential decay, 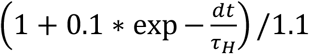 with *τ_H_* = 25 μs, to mimic the blinking component due to the protonation-deprotonation process of sfGFP, before fitting with the different models of diffusion. The code used for this simulation is available in GitHub (https://github.com/croelmiyn/Simulation_FCS_in_Bacteria) and via DOI: 10.5281/zenodo.5940484.

### Validation of fitting by the Ornstein-Uhlenbeck model

We first estimated the relation between the width *σ* of the potential well and the diameter *d* of the bacteria by fitting the ACF of the Brownian simulations with the OU model, fixing all parameters except *σ* to their ansatz values. The best fit was obtained for *σ* ≃ *d*/2 over the whole range of tested parameters. To mimic the fit procedure of experimental data and evaluate the accuracy of the diffusion coefficient estimation by the OU model (Figure 2 – figure supplement 3), we then fixed *σ* = *d*/2, and *ω*_0_ and *z*_0_ to their ansatz values, since they are measured independently in experiments, whereas the diffusion coefficient, number of particles *N* in the confocal volume, fraction of triplet excitation and background noise were taken as free parameters.

## Acknowledgements

We thank Lotte Søgaard-Andersen, Martin Thanbichler, Knut Drescher and Andreas Diepold for providing bacterial genomic DNA. We thank Silvia Espada Burriel for the assistance with the cellular density measurements and data analysis. This work was supported by the Max Planck Society.

## Competing Interests

The authors declare no competing interests.

## Notes

### Competing Interest Statement

The authors have declared no competing interest.

### Summary of Updates

Discussion revised to better highlight the novelty of results compared to existing literature; Figures 1 and 2 revised; Figure 4 added; Figure 5 (former Figure 4) revised.

