## Supplementary Material for "Dependence of diffusion in *Escherichia coli* cytoplasm on protein size, environmental conditions and cell growth"

This article contains the following supplementary files:

Figure 1 – supplementary table 1

Figure 1 – supplementary table 2

Figure 1 – figure supplement 1

Figure 1 – figure supplement 2

Figure 1 – figure supplement 3

Figure 1 – figure supplement 4

Figure 1 – figure supplement 5

Figure 1 – figure supplement 6

Figure 1 – figure supplement 7

Figure 1 – figure supplement 8

Figure 1 – figure supplement 9

Figure 1 – figure supplement 10

Figure 2 – figure supplement 1

Figure 2 – figure supplement 2

Figure 2 – figure supplement 3

Figure 2 – figure supplement 4

Figure 2 – figure supplement 5

Figure 2 – figure supplement 6

Figure 4 – figure supplement 1

Figure 5 – figure supplement 1

Figure 5 – figure supplement 2

Figure 5 – figure supplement 3

Figure 5 – figure supplement 4

Figure 5 – figure supplement 5

Figure 5 – figure supplement 6

Figure 5 – figure supplement 7

Appendix 1: Notes on the acquisition and analysis protocols for FCS measurements in bacterial cells

Appendix 2: Ornstein-Uhlenbeck model for confinement effect in FCS measurements

Appendix 3: Effective diffusion coefficient of two linked proteins

Appendix 4: Exact p-values for each figure

Appendix 5: Numerosity of the constructs and conditions for each experiment

Supplementary file 1: Uncut figures of the western blots analysis.

Supplementary file 2: Source data files for all figures.

### Supplementary figures

**Figure 1 – supplementary table 1. sfGFP-tagged library of cytoplasmic proteins**

| Protein name | Molecular mass of sfGFP-construct | Biological function | IPTG concentration for FCS (for FRAP) | Number of cells (FCS) | $\tau_D$ (µs; avg. ± SE) | α (Avg. ± SE) | Diffusion coefficient, FCS (µm²/s, Avg. ± SE) | Number of cells (FRAP) | Diffusion coefficient, FRAP (µm²/s, Avg. ± SE) |
|---|---|---|---|---|---|---|---|---|---|
| *sfGFP* | 26.9 | - | 5 µM (15 µM) | 52 | 561 ± 14 | 0.86 ± 0.01 | 14.7 ± 0.3 | 11 | 11.3 ± 1.3 |
| *YggX* | 39.2 | Probable Fe (2+)-trafficking protein | 5 µM (5 µM) | 8 | 611 ± 19 | 0.85 ± 0.01 | 12.9 ± 0.4 | 10 | 9.4 ± 1.6 |
| *ClpS* | 39.2 | ATP-dependent Clp protease adapter protein | 0 µM | 11 | 1054 ± 33 | 0.75 ± 0.01 | | | |
| *FolK* | 45.1 | 2-amino-4-hydroxy-6-hydroxymethyldihydropteridine pyrophosphokinase | 0 µM | 8 | 734 ± 24 | 0.87 ± 0.01 | 11.6 ± 0.4 | | |
| *Crr* | 45.2 | Component of glucose-specific phosphotransferase enzyme IIA | 0 µM | 14 | 1065 ± 36 | 0.87 ± 0.01 | | | |
| *UbiC* | 45.7 | Chorismate pyruvate-lyase | 15 µM | 14 | 1140 ± 58 | 0.87 ± 0.01 | | | |
| *ThpR* | 46.9 | RNA 2',3'-cyclic phosphodiesterase | | | Discarded due to instability of sfGFP-construct | | | | |
| *CoaE* | 49.6 | Dephospho-CoA kinase | 0 µM | 11 | 854 ± 47 | 0.87 ± 0.01 | 9.8 ± 0.6 | | |
| *Adk* | 50.6 | Adenylate kinase | 5 µM (15 µM) | 23 | 802 ± 26 | 0.88 ± 0.00 | 10.6 ± 0.4 | 16 | 9.8 ± 1.5 |
| *Cmk* | 51.7 | Cytidylate kinase | 5 µM | 16 | 1163 ± 58 | 0.87 ± 0.01 | | | |
| *NagD* | 54.1 | Ribonucleotide monophosphatase | | | Discarded due to non-uniform protein localization | | | | |
| *KdsB* | 54.6 | 3-deoxy-manno-octulosonate cytidylyltransferase | 0 µM | 11 | 1659 ± 70 | 0.84 ± 0.01 | | | |
| *Map* | 56.3 | Methionine aminopeptidase | 0 µM | 20 | 1830 ± 78 | 0.81 ± 0.01 | | | |
| *MmuM* | 60.4 | Homocysteine S-methyltransferase | 5 µM | 14 | 2241 ± 138 | 0.73 ± 0.01 | | | |
| *RihA* | 60.8 | Pyrimidine-specific ribonucleoside hydrolase | | | Discarded due to non-uniform protein localization | | | | |
| *PanE* | 60.8 | 2-dehydropantoate 2-reductase | 0 µM (5 µM) | 18 | 1059 ± 26 | 0.85 ± 0.01 | 7.8 ± 0.2 | 11 | 5.2 ± 0.6 |
| *SolA* | 67.9 | N-methyl-L-tryptophan oxidase | 0 µM | 7 | 795 ± 31 | 0.82 ± 0.01 | 9.9 ± 0.5 | | |
| *Pgk* | 68.1 | Phosphoglycerate kinase | 0 µM | 16 | 991 ± 41 | 0.90 ± 0.01 | 8.6 ± .0.3 | | |
| *EntC* | 69.9 | Isochorismate synthase | 15 µM | 15 | 1777 ± 119 | 0.82 ± 0.01 | | | |
| *AroA* | 73.1 | 3-phosphoshikimate 1-carboxyvinyltransferase | 5 µM | 9 | 995 ± 69 | 0.86 ± 0.01 | 8.7 ± 0.7 | | |
| *ThrC* | 74.1 | Threonine synthase | 0 µM | 14 | 908 ± 28 | 0.87 ± 0.01 | 9.1 ± 0.3 | | |
| *MurF* | 74.4 | UDP-N-acetylmuramoyl-tripeptide--D-alanyl-D-alanine ligase | 0 µM | 7 | 1008 ± 76 | 0.85 ± 0.02 | 8.3 ± 0.7 | | |
| *DsdA* | 74.9 | D-serine dehydratase | 0 µM | 14 | 1017 ± 53 | 0.89 ± 0.01 | 8.4 ± 0.4 | 10 | 7.8 ± 0.7 |
| *HemN* | 79.7 | Oxygen-independent coproporphyrinogen III oxidase | 0 µM | 13 | 1262 ± 54 | 0.86 ± 0.01 | 6.7 ± 0.4 | | |
| *PrpD* | 80.9 | 2-methylcitrate dehydratase | 0 µM | 12 | 1866 ± 140 | 0.84 ± 0.01 | | | |
| *DnaK* | 96.0 | Molecular chaperone | 5 µM | 10 | 2296 ± 78 | 0.76 ± 0.01 | | | |
| *MalZ* | 96.0 | Maltodextrin glucosidase | 0 µM | 9 | 3725 ± 229 | 0.77 ± 0.01 | | | |
| *GlcB* | 107.5 | Malate synthase G | 5 µM (15 µM) | 16 | 1315 ± 45 | 0.86 ± 0.01 | 6.4 ± 0.2 | 10 | 6.7 ± 1.1 |
| *MetE* | 111.7 | 5-methyltetrahydropteroyltriglutamate--homocysteine methyltransferase | 5 µM | 8 | 1137 ± 53 | 0.87 ± 0.01 | 7.4 ± 0.3 | | |
| *LeuS* | 124.2 | Leucine--tRNA ligase | 0 µM | 14 | 1637 ± 75 | 0.86 ± 0.01 | 5.1 ± 0.2 | | |
| *AcnA* | 124.7 | Aconitate hydratase A | 5 µM (15 µM) | 19 | 1415 ± 56 | 0.86 ± 0.01 | 6.1 ± 0.2 | 10 | 4.3 ± 0.4 |
| *MetH* | 163.0 | Methionine synthase | 0 µM (5 µM) | 9 | 1402 ± 45 | 0.81± 0.01 | 5.8 ± 0.1 | 15 | 4.0 ± 0.5 |

**Figure 1 – supplementary table 2. Strains and plasmids used in this study**

| Strain or plasmid | Relevant genotype or phenotype | Reference or source |
|---|---|---|
| | ***Strains*** | |
| *E. coli* W3110 | W3110 derivative with functional RpoS [rpoS396(Am)] | (Serra Diego et al.) |
| NB63 | W3110 Δ*clpA* | This work |
| RC111 | W3110 *Δflu, fliC::KanR* | This work |
| | ***Plasmids*** | |
| pTrc99A | Amp$^r$; expression vector; pBR ori; *trc* promoter, IPTG inducible | (Amann, Ochs, and Abel 1988) |
| pCP20 | Amp$^r$, Cam$^r$; *flp* | (Cherepanov and Wackernagel 1995) |
| pNB1 | Amp$^r$; *sfGFP* in pTrc99A | This work |
| pNB3 | Amp$^r$; *Adk-sfGFP* in pTrc99A | This work |
| pNB4 | Amp$^r$; *CoaE-sfGFP* in pTrc99A | This work |
| pNB5 | Amp$^r$; *Cmk-sfGFP* in pTrc99A | This work |
| pNB6 | Amp$^r$; *Pgk-sfGFP* in pTrc99A | This work |
| pNB7 | Amp$^r$; *MmuM-sfGFP* in pTrc99A | This work |
| pNB8 | Amp$^r$; *PrpD-sfGFP* in pTrc99A | This work |
| pNB9 | Amp$^r$; *DsdA-sfGFP* in pTrc99A | This work |
| pNB11 | Amp$^r$; *GlcB-sfGFP* in pTrc99A | This work |
| pNB13 | Amp$^r$; *HemN-sfGFP* in pTrc99A | This work |
| pNB14 | Amp$^r$; *Map*$^{WT}$-*sfGFP* in pTrc99A | This work |
| pNB15 | Amp$^r$; *ThrC-sfGFP* in pTrc99A | This work |
| pNB16 | Amp$^r$; *MalZ-sfGFP* in pTrc99A | This work |
| pNB17 | Amp$^r$; *EntC-sfGFP* in pTrc99A | This work |
| pNB18 | Amp$^r$; *ThpR-sfGFP* in pTrc99A | This work |
| pNB19 | Amp$^r$; *AroA-sfGFP* in pTrc99A | This work |
| pNB20 | Amp$^r$; *ClpS*$^{WT}$-*sfGFP* in pTrc99A | This work |
| pNB21 | Amp$^r$; *Crr-sfGFP* in pTrc99A | This work |
| pNB22 | Amp$^r$; *KdsB-sfGFP* in pTrc99A | This work |
| pNB23 | Amp$^r$; *LeuS-sfGFP* in pTrc99A | This work |
| pNB24 | Amp$^r$; *MurF-sfGFP* in pTrc99A | This work |
| pNB25 | Amp$^r$; *NagD-sfGFP* in pTrc99A | This work |
| pNB26 | Amp$^r$; *RihA-sfGFP* in pTrc99A | This work |
| pNB27 | Amp$^r$; *SolA-sfGFP* in pTrc99A | This work |
| pNB28 | Amp$^r$; *UbiC-sfGFP* in pTrc99A | This work |
| pNB29 | Amp$^r$; *PanE-sfGFP* in pTrc99A | This work |
| pNB30 | Amp$^r$; *FolK-sfGFP* in pTrc99A | This work |
| pNB39 | Amp$^r$; *AcnA-sfGFP* in pTrc99A | This work |
| pNB40 | Amp$^r$; *MetE-sfGFP* in pTrc99A | This work |
| pNB42 | Amp$^r$; *MetH-sfGFP* in pTrc99A | This work |
| pNB44 | Amp$^r$; *YggX-sfGFP* in pTrc99A | This work |
| pNB45 | Amp$^r$; *Adk*$^{C.c.-}$*sfGFP* in pTrc99A | This work |
| pNB46 | Amp$^r$; *Adk*$^{V.c.}$-*sfGFP* in pTrc99A | This work |
| pNB47 | Amp$^r$; *Adk*$^{M.x.}$-*sfGFP* in pTrc99A | This work |
| pNB48 | Amp$^r$; *AcnA*$^{M.x.}$-*sfGFP* in pTrc99A | This work |
| pNB49 | Amp$^r$; *AcnA*$^{V.c.}$-*sfGFP* in pTrc99A | This work |
| pNB51 | Amp$^r$; *Adk*$^{B.s.}$-*sfGFP* in pTrc99A | This work |
| pNB52 | Amp$^r$; *AcnA*$^{B.s.}$-*sfGFP* in pTrc99A | This work |
| pNB54 | Amp$^r$; *Adk*$^{Y.e.}$-*sfGFP* in pTrc99A | This work |
| pNB56 | Amp$^r$; *Pgk*$^{C.c.}$-*sfGFP* in pTrc99A | This work |
| pNB58 | Amp$^r$; *Pgk*$^{V.c.}$-*sfGFP* in pTrc99A | This work |
| pNB59 | Amp$^r$; *Pgk*$^{M.x.}$*sfGFP* in pTrc99A | This work |
| pNB60 | Amp$^r$; *AcnA*$^{Y.e.}$-*sfGFP* in pTrc99A | This work |
| pNB61 | Amp$^r$; *DnaK*$^{WT}$-*sfGFP* in pTrc99A | This work |
| pNB62 | Amp$^r$; *Map*$^{K211E\_K218E\_K224E\_K226E}$-*sfGFP* in pTrc99A | This work |
| pNB63 | Amp$^r$; *Map*$^{K211A\_K218A\_K224A\_K226A}$-*sfGFP* in pTrc99A | This work |
| pNB64 | Amp$^r$; *DnaK*$^{V436F}$ -*sfGFP* in pTrc99A | This work |
| pNB66 | Amp$^r$; *ClpS*$^{D35A\_D36A\_H66A}$-*sfGFP* in pTrc99A | This work |

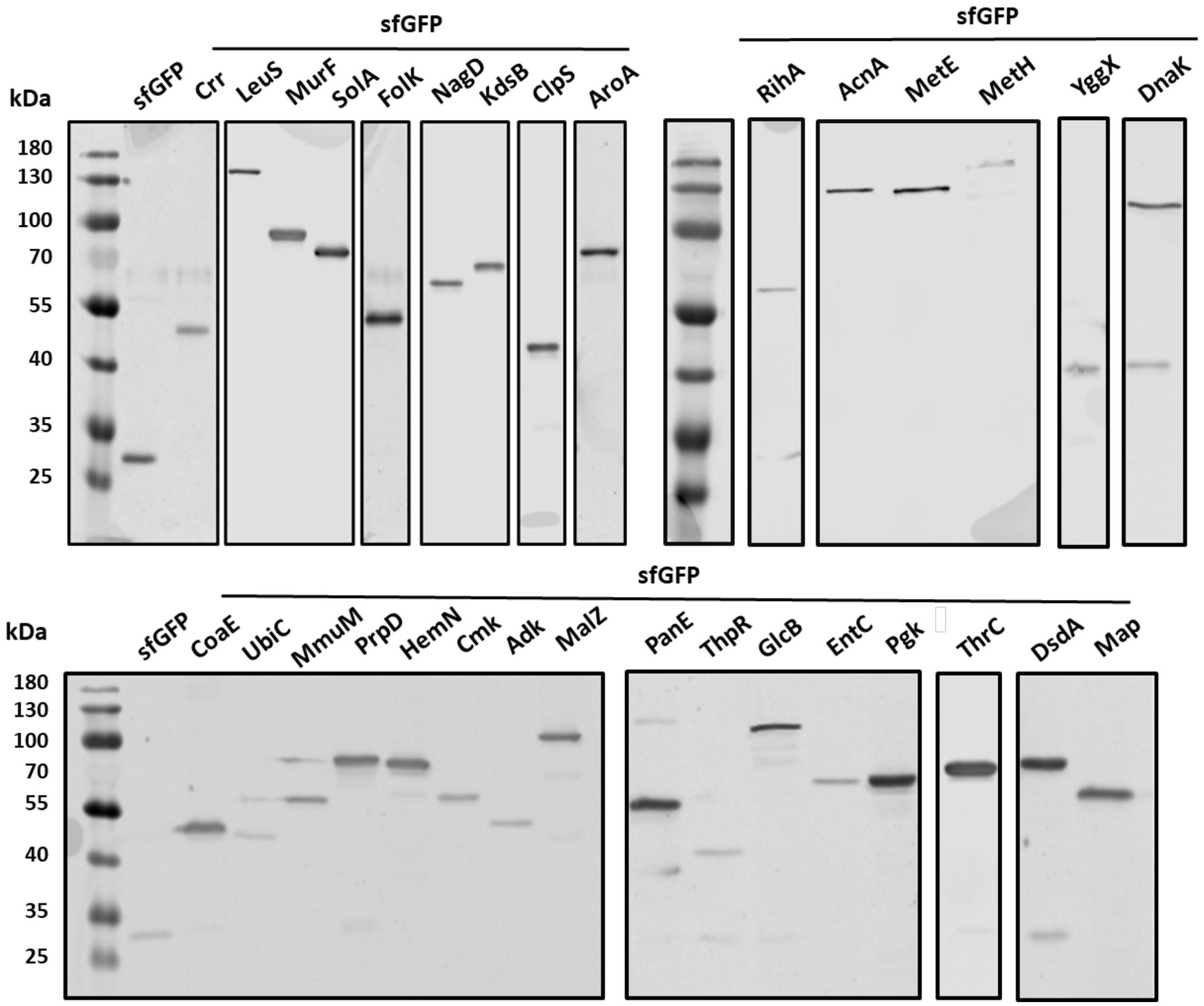


**Figure 1 – figure supplement 1. Expression analysis for all *E. coli* protein constructs made in this study.** Expression of proteins was analyzed by SDS-PAGE and immunoblotting using a primary antibody specific for GFP. All fusion proteins displayed a dominant band corresponding to the expected molecular mass of the full-length fusion (Figure 1 – supplementary table 1).

**Figure 1 – figure supplement 1 – source data**

Uncropped western blot images for Figure 1 – figure supplement 1

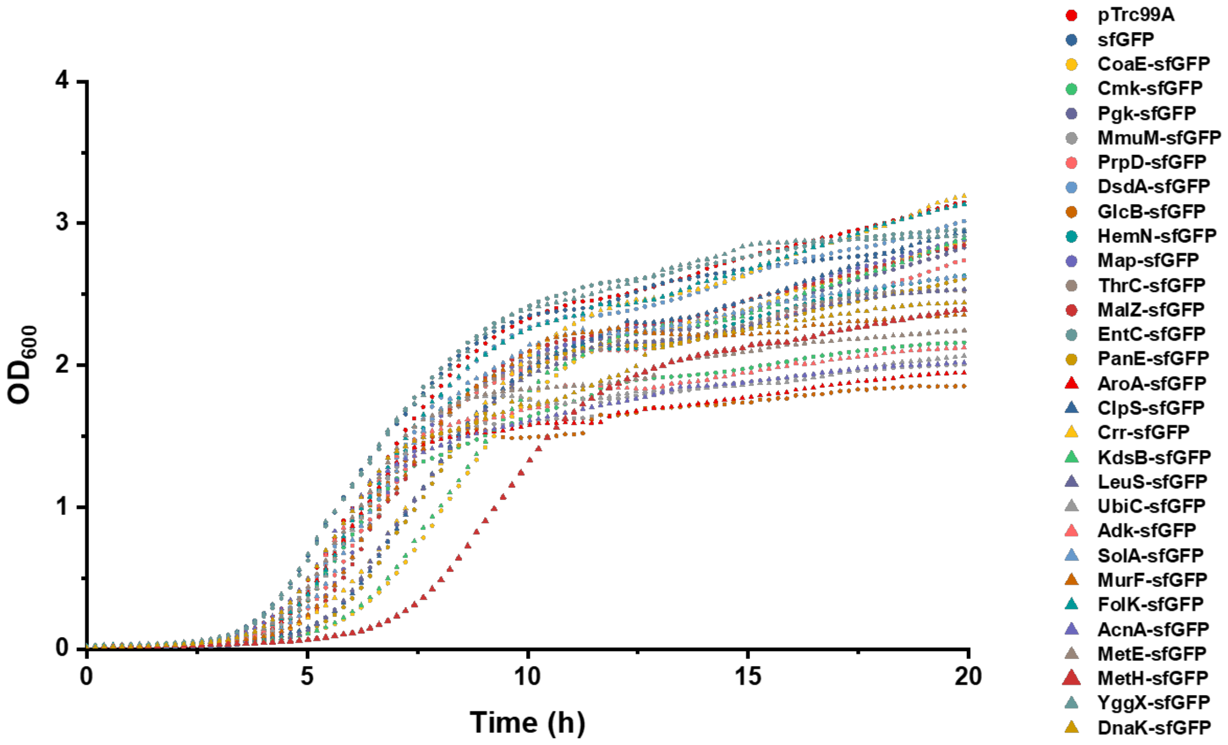


**Figure 1 – figure supplement 2. Growth curves of *E. coli* strains expressing tested sfGFP-tagged proteins.** Growth curves of strains expressing sfGFP, sfGFP-tagged proteins or carrying the control empty vector pTrc99A were measured at 37°C. The optical density of cultures was monitored at 600 nm ($OD_{600}$) and the measured values were normalized for an optical path of 1 cm.

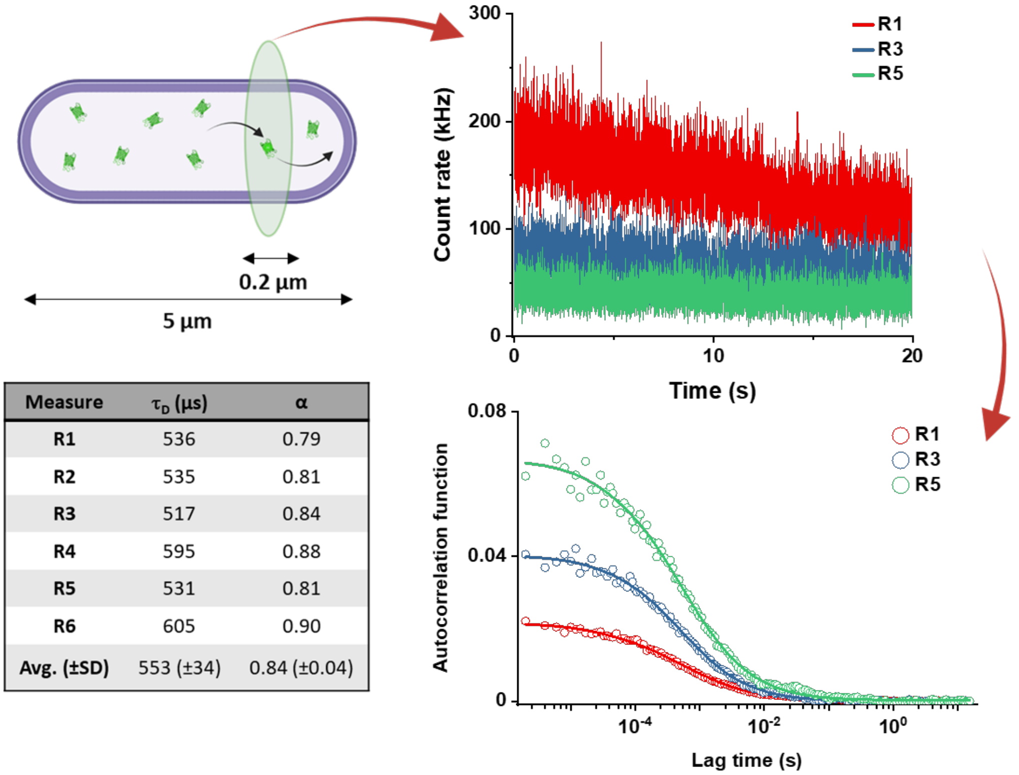


| Measure | $\tau_D$ (µs) | α |
|---|---|---|
| **R1** | 536 | 0.79 |
| **R2** | 535 | 0.81 |
| **R3** | 517 | 0.84 |
| **R4** | 595 | 0.88 |
| **R5** | 531 | 0.81 |
| **R6** | 605 | 0.90 |
| **Avg. (±SD)** | 553 (±34) | 0.84 (±0.04) |

**Figure 1 – figure supplement 3. Workflow of a typical FCS experiment.** The focus of the confocal microscope is positioned near one pole in the cell of interest, and six subsequent acquisitions of the fluorescence intensity of 20 seconds each are performed. The example for sfGFP shows only traces for R1, R3 and R5 measurements (different colors). The autocorrelation functions (ACFs) are then calculated independently for each acquisition. The values for $\tau_D$ and $\alpha$ are extracted independently from the fit to each ACF using the anomalous diffusion model (solid lines) and then averaged to obtain the $\tau_D$ and $\alpha$ for each individual cell.

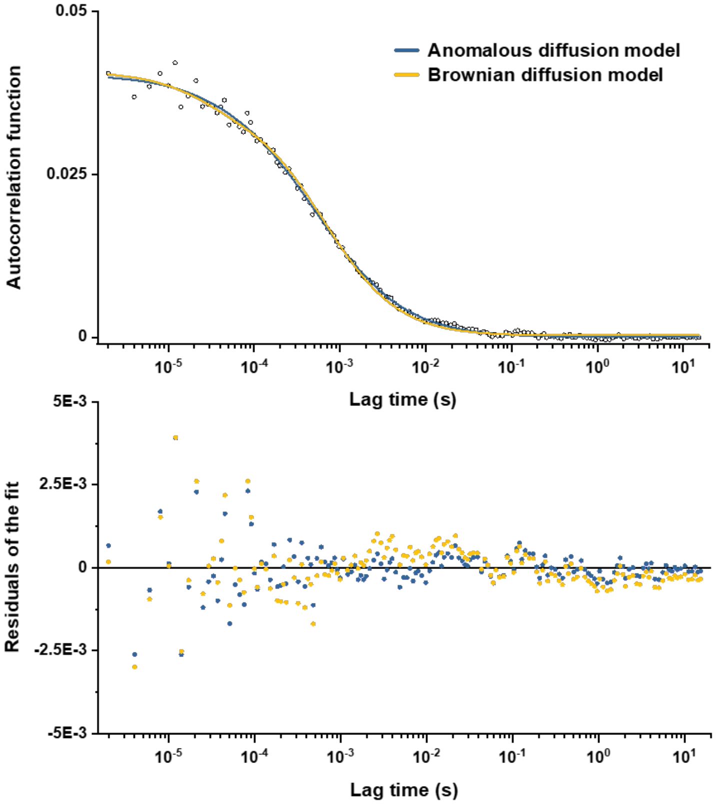


**Figure 1 – figure supplement 4. Comparison between fits of the experimental data with Brownian and anomalous diffusion models.** The experimental data (here the example for the R3 measurement from Figure 1 – figure supplement 3) were fitted by the model of free Brownian diffusion and by the anomalous diffusion model as indicated by different colors (upper panel), with the corresponding values of residuals (lower panel).

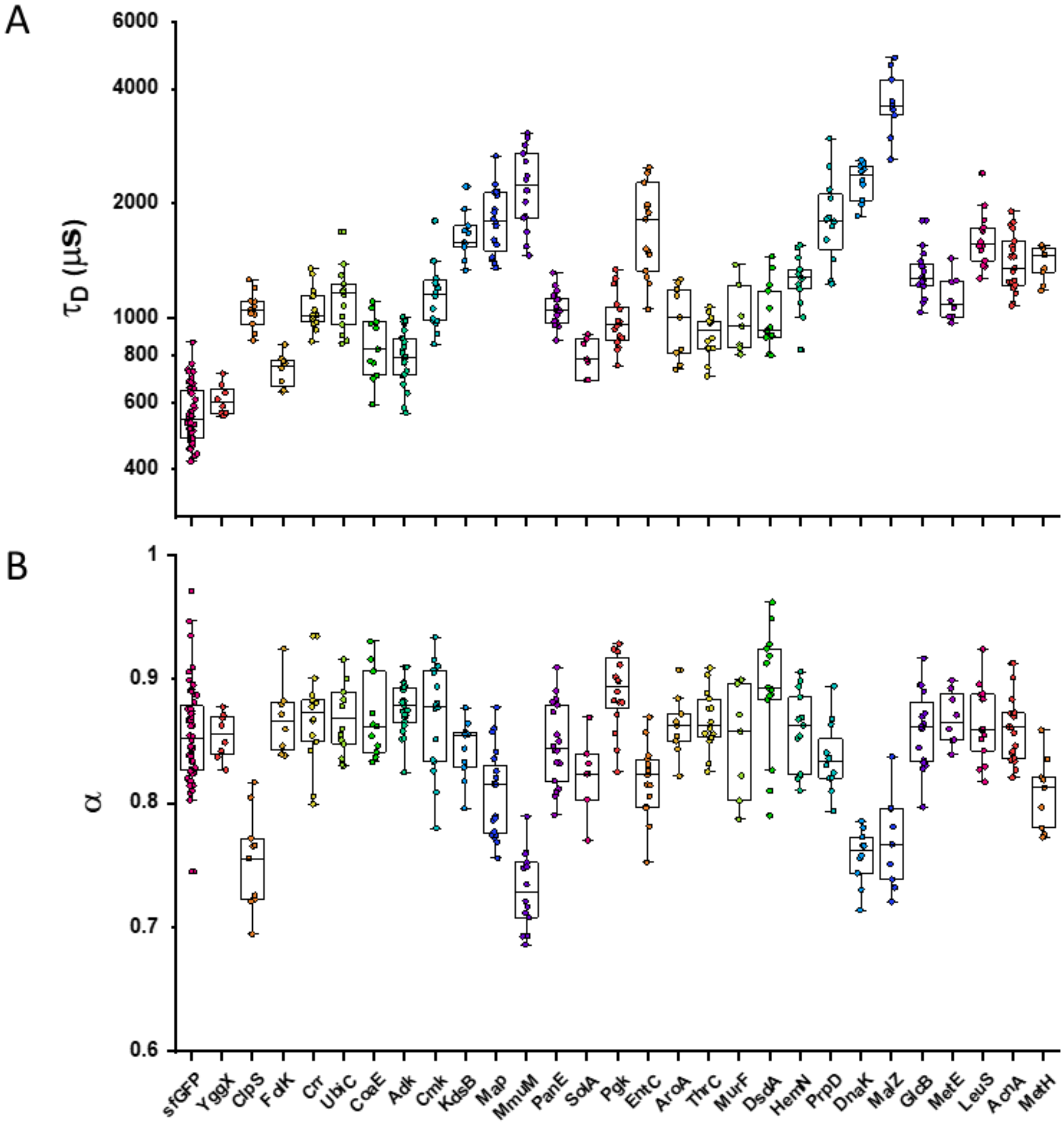


**Figure 1 – figure supplement 5. Individual measurements of $\tau_D$ and $\alpha$ for all *E. coli* protein constructs included in the analysis of mass dependence.** Each dot in the box plot represents the values of $\tau_D$ (A) and $\alpha$ (B) for one individual cell. The averages values calculated from these datasets are shown in Figure 1.

**Figure 1 – figure supplement 5 – source data**

Individual values of $\tau_D$ from Figure 1 – figure supplement 5A

Individual values of $\alpha$ from Figure 1 – Figure supplement 5B

A

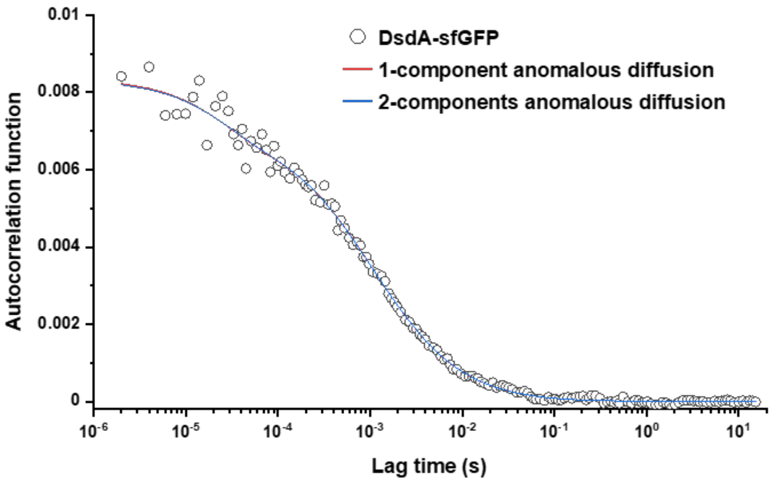


B

| Model | $\tau_D$ (µs; avg. ± SE) | α (avg. ± SE) |
|---|---|---|
| **1-component** | 1017 ± 53 | 0.89 ± 0.01 |
| **2-components** | 1092 ± 62 | 0.88 ± 0.02 |

**Figure 1 – figure supplement 6. Comparison between one-component and two-components anomalous diffusion fit for DsdA-sfGFP.** (A) Fitting the experimental data for DsdA-sfGFP with a two-component model for anomalous diffusion, where the fast component corresponding to free sfGFP was fixed to 15% (corresponding to the measured degree of fusion protein degradation) and with $\tau_D$ =561 µs and $\alpha$ = 0.86, i.e. the average values obtained for sfGFP. For comparison, standard fitting with a one-component anomalous diffusion model is also shown. (B) The fitted values of $\tau_D$ with the two-component and one-component model.

**Figure 1 – figure supplement 6 – source data**

Individual values used to calculate mean and standard error of the mean values of $\tau_D$ and α values from Figure 1 – figure supplement 6B.

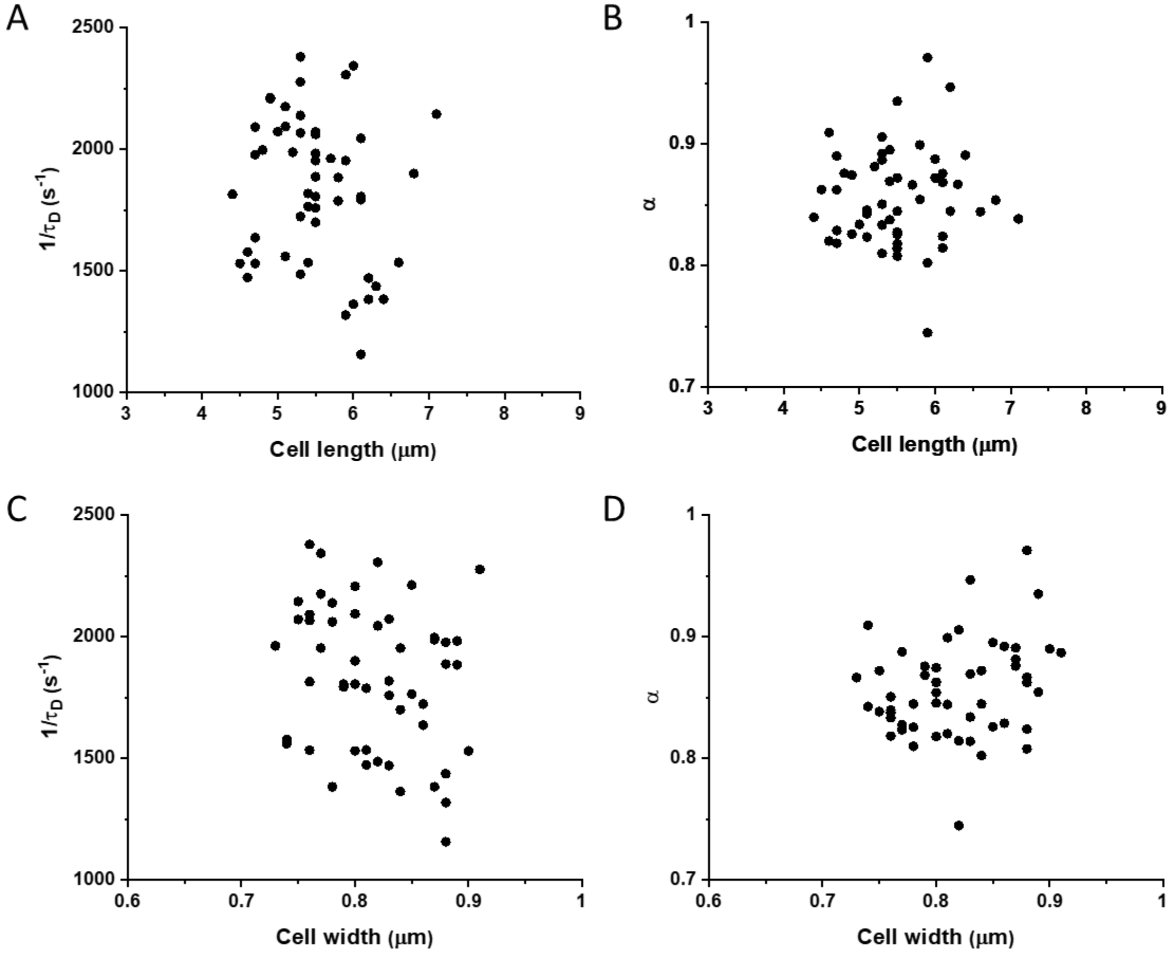


**Figure 1 – figure supplement 7. Mobility (1/$\tau_D$) and anomaly of diffusion ($\alpha$) of sfGFP in individual cells with different width and length.** Values of 1/$\tau_D$ (A, C) and α (B, D) for single cells expressing sfGFP plotted against the length (A, B) or width (C, D) of respective cell.

**Figure 1 – figure supplement 7 – source data**

Individual values of 1/$\tau_D$ and measurements of cell length from Figure 1 – figure supplement 7A.

Individual values of $\alpha$ and measurements of cell length from Figure 1 – figure supplement 7B.

Individual values of 1/$\tau_D$ and measurements of cell width from Figure 1 – figure supplement 7C.

Individual values of $\alpha$ and measurements of cell width from Figure 1 – figure supplement 7D.

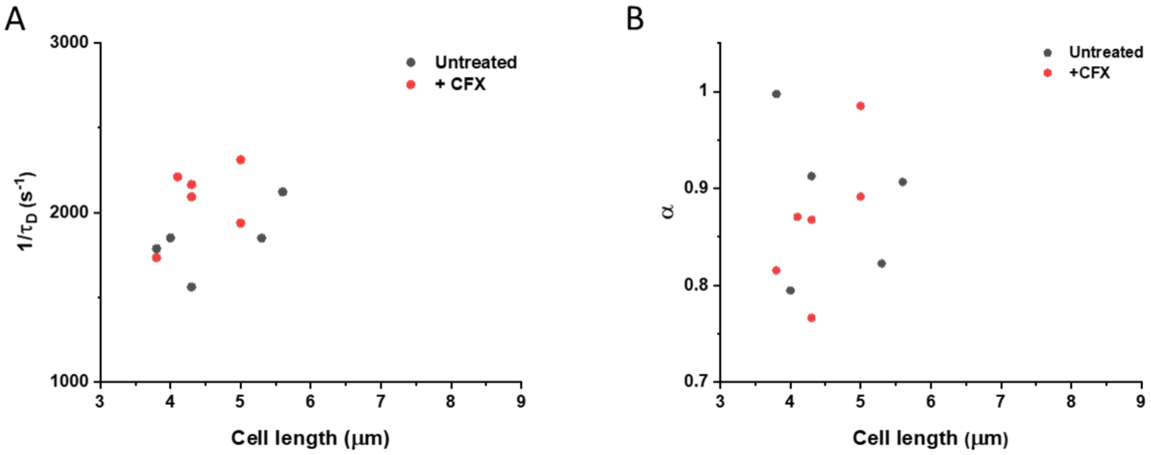


**Figure 1 – figure supplement 8. Comparison of protein mobility in cephalexin-treated and untreated cells.** Mobility of sfGFP measured in single cells treated with cephalexin (red dots) or in control untreated cells (black dots). Only cells of similar length were chosen for this comparison. No significant difference in protein mobility ($1/\tau_D$; p = 0.08) or anomaly of diffusion ($\alpha$; p = 0.67) is observed.

**Figure 1 – figure supplement 8 – source data**

Individual values of $1/\tau_D$ and measurements of cell length from Figure 1 – figure supplement 8A.

Individual values of $\alpha$ and measurements of cell length from Figure 1 – figure supplement 8B.

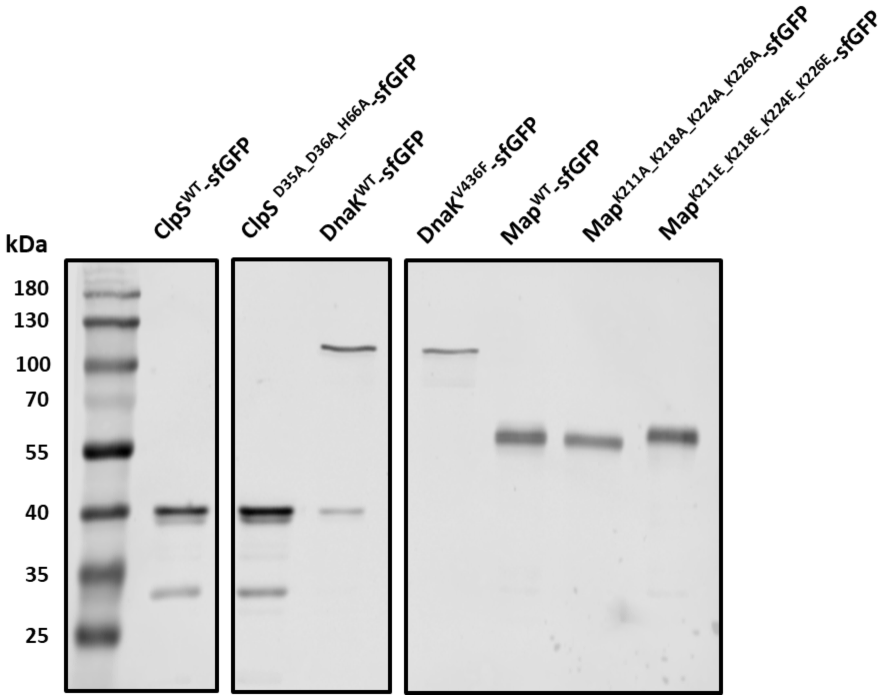


**Figure 1 – figure supplement 9. Expression analysis for the mutants with impaired interactions.** The expression of indicated point mutants of ClpS, DnaK and Map was analyzed by SDS-PAGE and immunoblotting using a primary antibody specific for GFP. All mutants displayed a dominant band corresponding to the expected molecular mass of the full-length fusion, and comparable to that of the wild-type counterpart. ClpS$^{D35A_D36A_H66A}$-sfGFP was measured in the same *ΔclpA* background as subsequently used for the FCS experiments.

**Figure 1 – figure supplement 9 – source data**

Uncropped western blot image for Figure 1 – figure supplement 9.

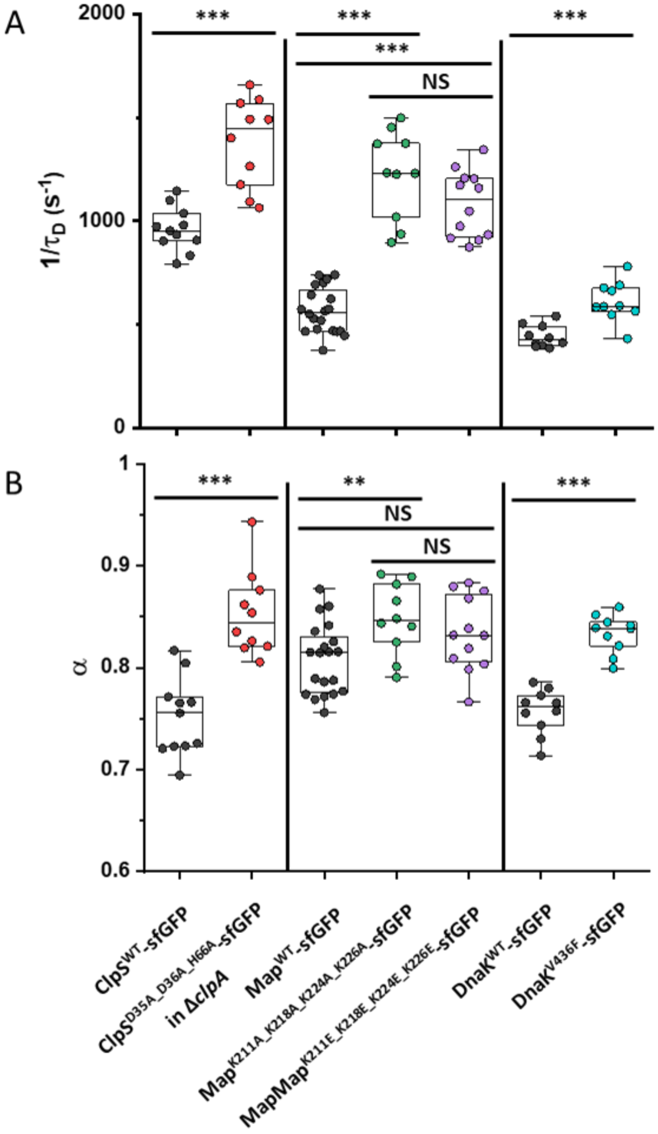


**Figure 1 – figure supplement 10. Mobility ($1/\tau_D$) and anomaly of diffusion ($\alpha$) of ClpS, Map and DnaK and of indicated mutants with disrupted protein interactions.** Each dot in the box plot represents the values of $1/\tau_D$ (A) and $\alpha$ (B) for one individual cell. ClpS mutant was measured in *ΔclpA* background. *** $p<0.0001$; * $p<0.05$; NS: no statistically significant difference in a two-tailed heteroscedastistic *t*-test.

**Figure 1 – figure supplement 10 – source data**

Individual values of $1/\tau_D$ from Figure 1 – figure supplement 10A.

Individual values of $\alpha$ from Figure 1 – figure supplement 10B.

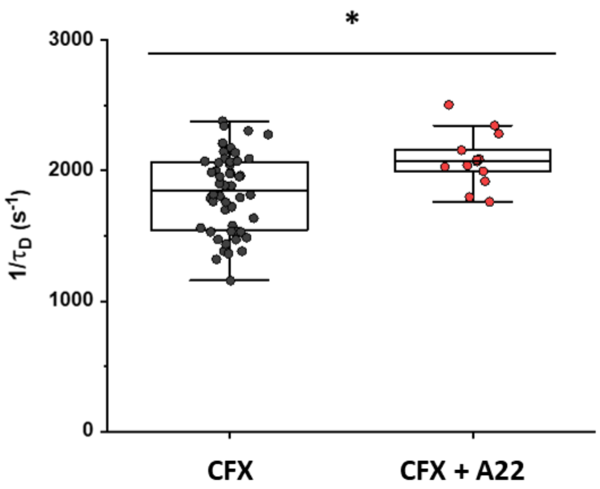


**Figure 2 – figure supplement 1. Mobility of sfGFP in cells treated with cephalexin (CFX) or the combination of cephalexin and A22.** Each dot in the box plot represents the values of $1/\tau_D$ for one individual cell. Cultures are grown for ~ 3.5 hours in absence or presence of A22 before being both treated for 45 min with cephalexin. * $p<0.05$ in a two-tailed heteroscedastistic *t*-test.

**Figure 2 – figure supplement 1 – source data**

Individual values of $1/\tau_D$ from Figure 2 – figure supplement 1.

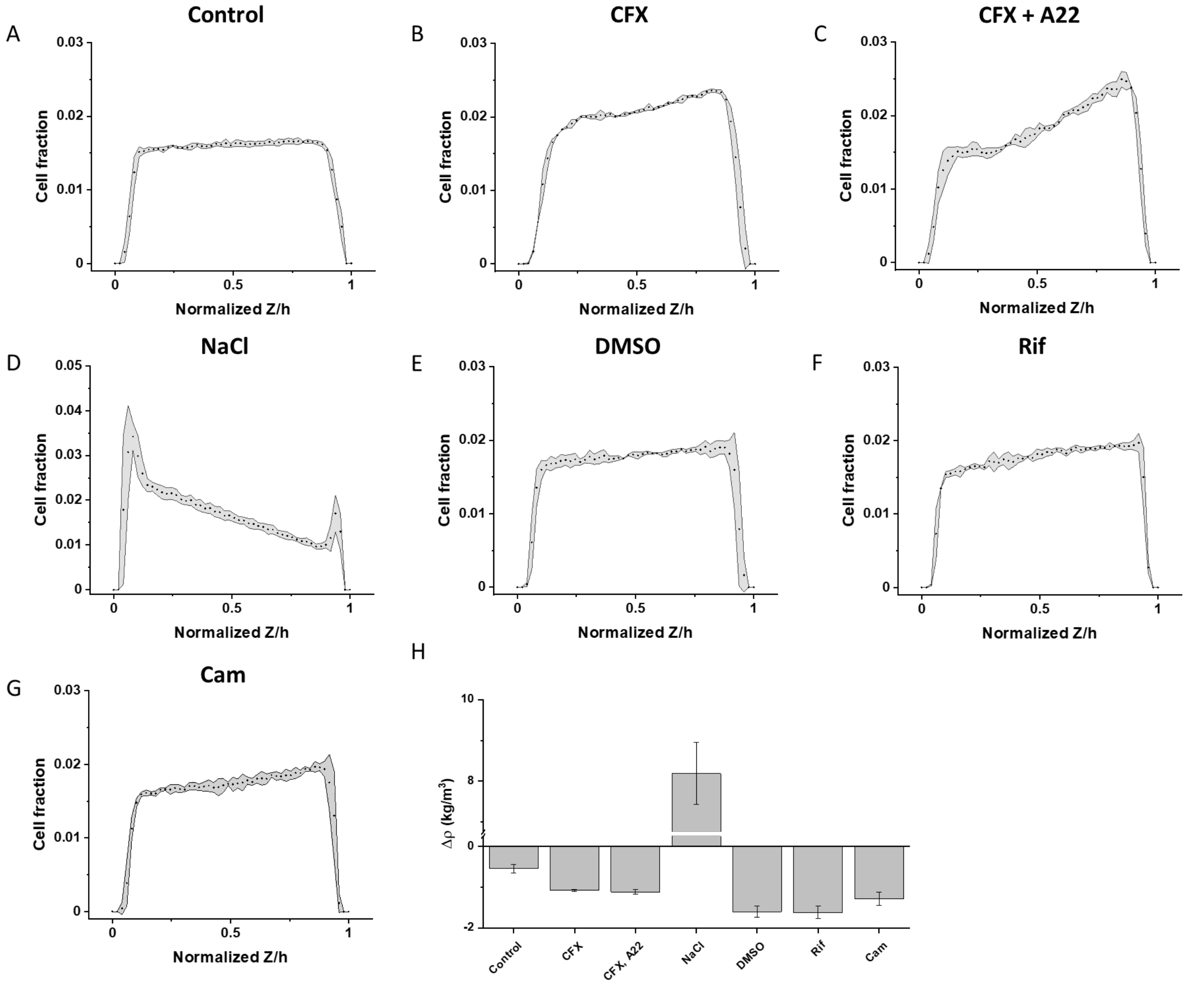


**Figure 2 – figure supplement 2. Sedimentation assay of cellular density for indicated treatments.** Sedimentation assays were performed using non-motile and non-aggregating (*ΔfliC Δflu*) variant of the same *E. coli* strain W3110 $RpoS^+$ strain as used in other experiments. Cells were grown and treated as described in the correspondent sections and assayed for their density in motility buffer (MB) containing 20% iodixanol to match the density of control untreated cells (A). Treatments with cephalexin (CFX; B), cephalexin and A22 (C), 100 mM NaCl (D), DMSO (E), rifampicin (Rif; F), and chloramphenicol (Cam; G) are shown. Dots represent the cell fraction at each given Z position normalized on the total height of the microfluidic channel (50 µm). The grey shadings indicate the standard deviation of the three technical replicates. (H) Calculated values of cellular density mismatch from MB with 20% iodixanol. Error bars represent the standard error of the mean. Calculations were performed as described in Materials and methods.

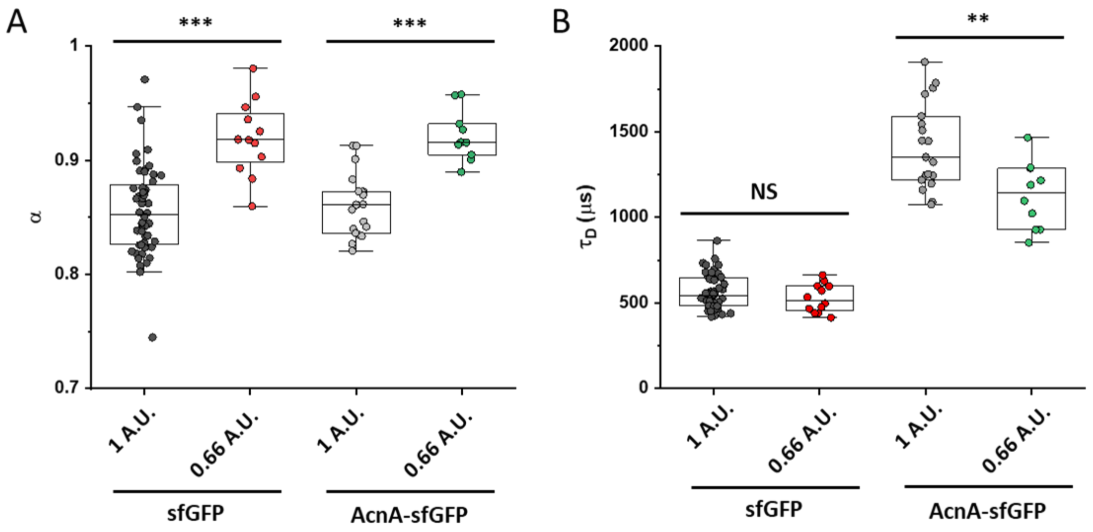


**Figure 2 – figure supplement 3. Apparent anomaly of diffusion and residence time for different pinhole sizes.** FCS measurements of sfGFP and of AcnA-sfGFP were performed at the suboptimal pinhole size of 0.66 Airy units (A.U.) and the values of the anomaly of diffusion ($\alpha$; A) and residence time ($\tau_D$; B). As expected, $\tau_D$ of AcnA-sfGFP scales according to the reduction in the beam waist. The $\tau_D$ of sfGFP shows the same trend, but less pronounced.

**Figure 2 – figure supplement 3 – source data**

Individual values of $\alpha$ from Figure 2 – figure supplement 3A.

Individual values of $\tau_D$ from Figure 2 – figure supplement 3B.

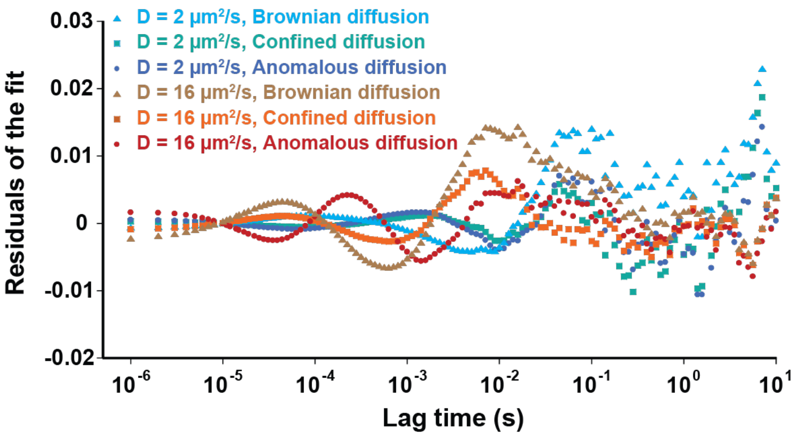


**Figure 2 – figure supplement 4. Residuals of fitting the simulated ACFs with different models.** Simulated fluorescence intensity ACF data were fitted by the models of anomalous and Brownian diffusion, as well as by the OU model of Brownian diffusion under confinement, as indicated.

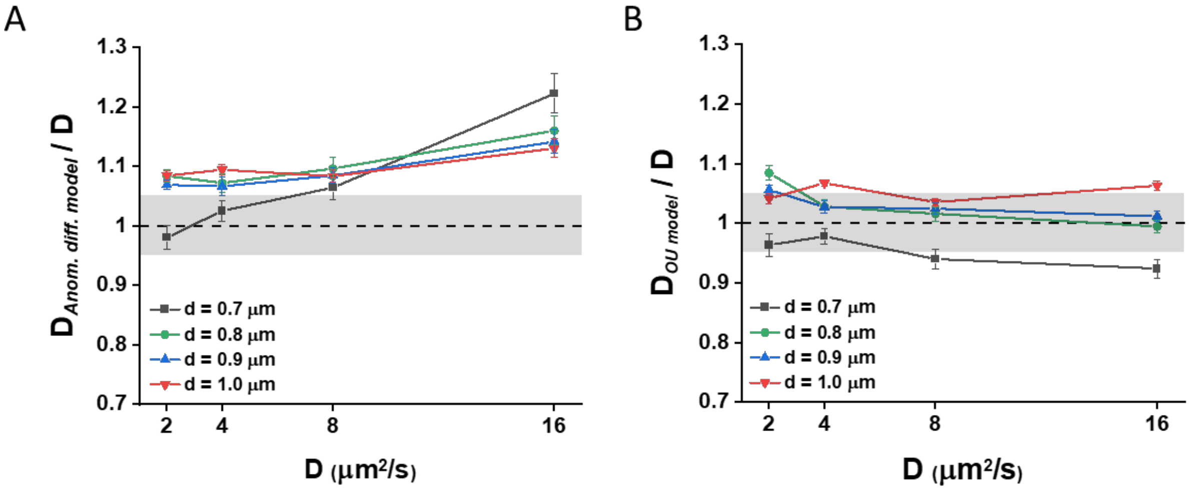


**Figure 2 – figure supplement 5. Diffusion coefficients fitted from simulation data.** Diffusion coefficients computed ($D = \omega_0^2/4\tau_D$) from the diffusion times extracted from the fit of the Brownian simulation data with (A) the anomalous diffusion model or (B) the Ornstein-Uhlenbeck (OU) model of Brownian diffusion under confinement at various value of the ansatz *D* and of the cell diameter d, normalized by the ansatz *D*. The grey areas represent ± 5% accuracy.

**Figure 2 – figure supplement 5 – source data**

Average and error from simulation in Figure 2 – figure supplement 5A.

Average and error from simulation in Figure 2 – figure supplement 5B.

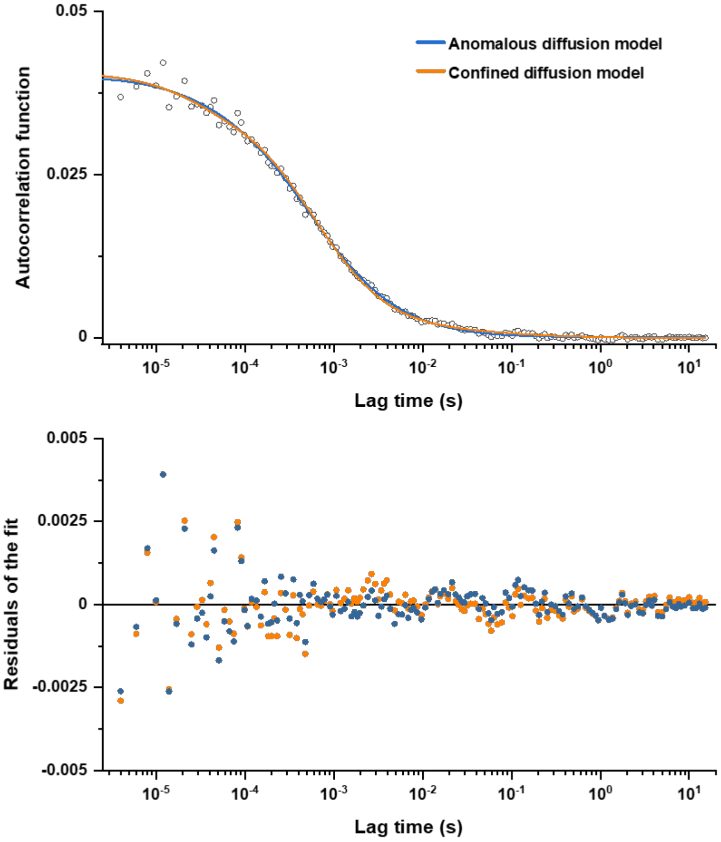


**Figure 2 – figure supplement 6. Comparison between fits of the experimental data with confined diffusion and anomalous diffusion models.** The experimental data (here the example for the R3 measurement from Figure 1 – figure supplement 2) were fitted by the model of confined diffusion and by the anomalous diffusion model as indicated by different colors (upper panel), yielding comparable values of residuals (lower panel).

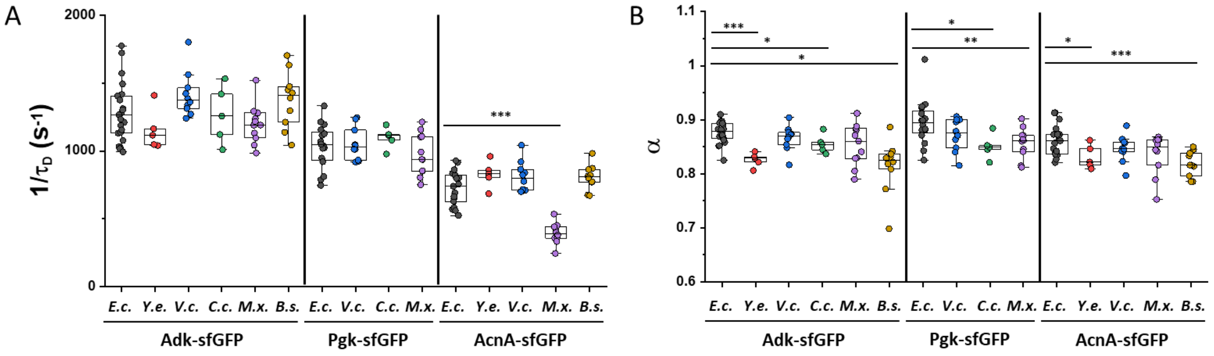


**Figure 4 – figure supplement 1. Mobility of homologous proteins from other bacterial species in *E. coli*.** Each dot represents the protein mobility ($1/\tau_D$; A) or the anomalous diffusion exponent ($\alpha$; B) from one individual cell expressing the indicated sfGFP fusions to homologues of Adk, Pgk and AcnA from indicated bacterial species (*E.c.* = *Escherichia coli; Y.e.* = *Yersinia enterocolitica; V.c.* = *Vibrio cholerae; C.c.* = *Caulobacter crescentus; M.x.* = *Myxococcus xanthus; B.s.* = *Bacillus subtilis*) compared with that of their counterpart from *E. coli*. *** $p<0.0001$; ** $p<0.001$; * $p<0.05$ in a two-tailed heteroscedastistic *t*-test. When not indicated, no statistically significant difference is observed.

**Figure 4 – figure supplement 1 – source data**

Individual values of $1/\tau_D$ from Figure 4 – figure supplement 1A.

Individual values of $\alpha$ from Figure 4 – figure supplement 1B.

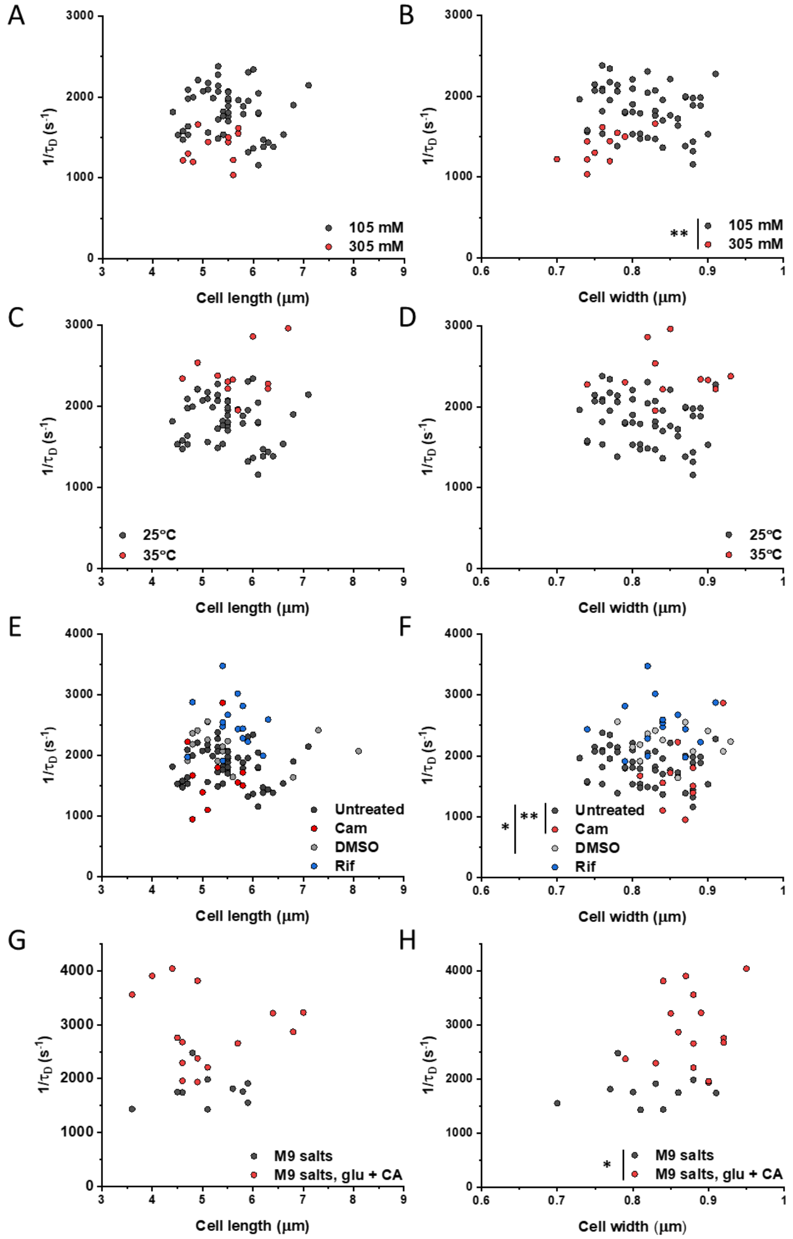


**Figure 5 – figure supplement 1. Mobility of sfGFP as a function of length and width of individual cells upon indicated perturbations.** The mobility ($1/\tau_D$) of sfGFP in single cells plotted as a function of the respective cell length (A, C, E, G) or cell width (B, D, F, H) at different ionic strengths (A, B), environmental temperatures (C, D), after treatments with different antibiotics (E, F) and cell growth (G, H). Significance analysis was performed for the respective cell dimension. When not indicated, no significant difference is observed. ** p<0.001; * p<0.05 in a two-tailed heteroscedastistic *t*-test.

**Figure 5 – figure supplement 1– source data**

Individual measurements of cell length from Figure 5 – figure supplement 1A.

Individual measurements of cell width from Figure 5 – figure supplement 1B.

Individual measurements of cell length from Figure 5 – figure supplement 1C.

Individual measurements of cell width from Figure 5 – figure supplement 1D.

Individual measurements of cell length from Figure 5 – figure supplement 1E.

Individual measurements of cell width from Figure 5 – figure supplement 1F.

Individual measurements of cell length from Figure 5 – figure supplement 1G.

Individual measurements of cell width from Figure 5 – figure supplement 1H.

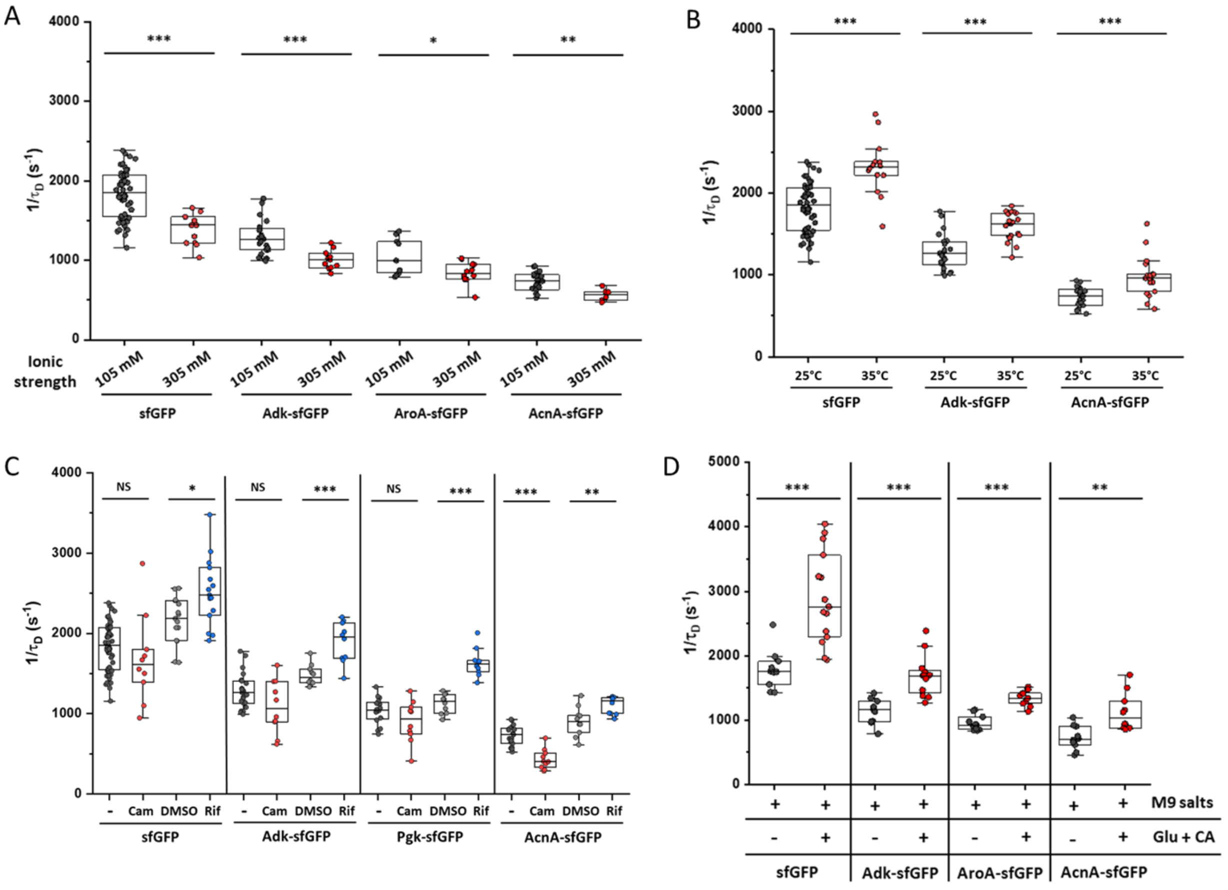


**Figure 5 – figure supplement 2. Effect of different perturbations on protein mobility (1/$\tau_D$) in individual cells.** Experiments for ionic strength (A), environmental temperature (B), antibiotics treatment (C), and growth rate (D) are shown for indicated proteins. Individual measurements and significance analysis of data from figure 5. Each dot in the box plot represents the value of 1/$\tau_D$ for one individual cell measured in the indicated condition. *** $p<0.0001$; ** $p<0.001$; * $p<0.05$; NS: no statistically significant difference in a two-tailed heteroscedastistic *t*-test.

**Figure 5 – figure supplement 2 – source data**

Individual values of 1/$\tau_D$ from Figure 5 – figure supplement 2A.

Individual values of 1/$\tau_D$ from Figure 5 – figure supplement 2B.

Individual values of 1/$\tau_D$ from Figure 5 – figure supplement 2C.

Individual values of 1/$\tau_D$ from Figure 5 – figure supplement 2D.

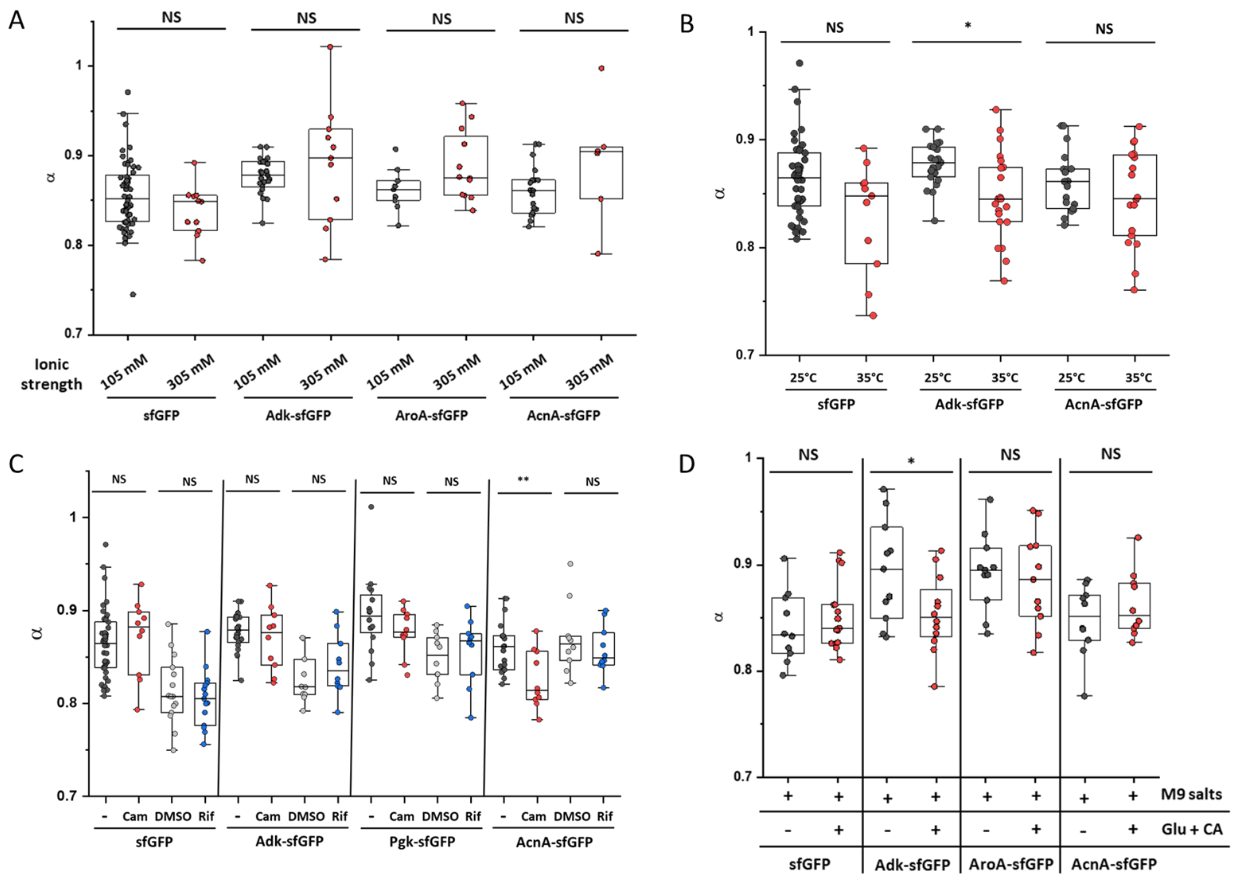


**Figure 5 – figure supplement 3. Effect of different perturbations on the anomaly of protein diffusion (*α*) in individual cells.** Experiments for ionic strength (A), environmental temperature (B), antibiotics treatment (C), and growth rate (D) are shown for indicated proteins. Each dot in the box plot represents the value of *α* for one individual cell measured in the indicated condition. ** $p<0.001$; * $p<0.05$; NS: no statistically significant difference in a two-tailed heteroscedastistic *t*-test.

**Figure 5 – figure supplement 3 – source data**

Individual values of *α* from Figure 5 – figure supplement 3A.

Individual values of *α* from Figure 5 – figure supplement 3B.

Individual values of *α* from Figure 5 – figure supplement 3C.

Individual values of *α* from Figure 5 – figure supplement 3D.

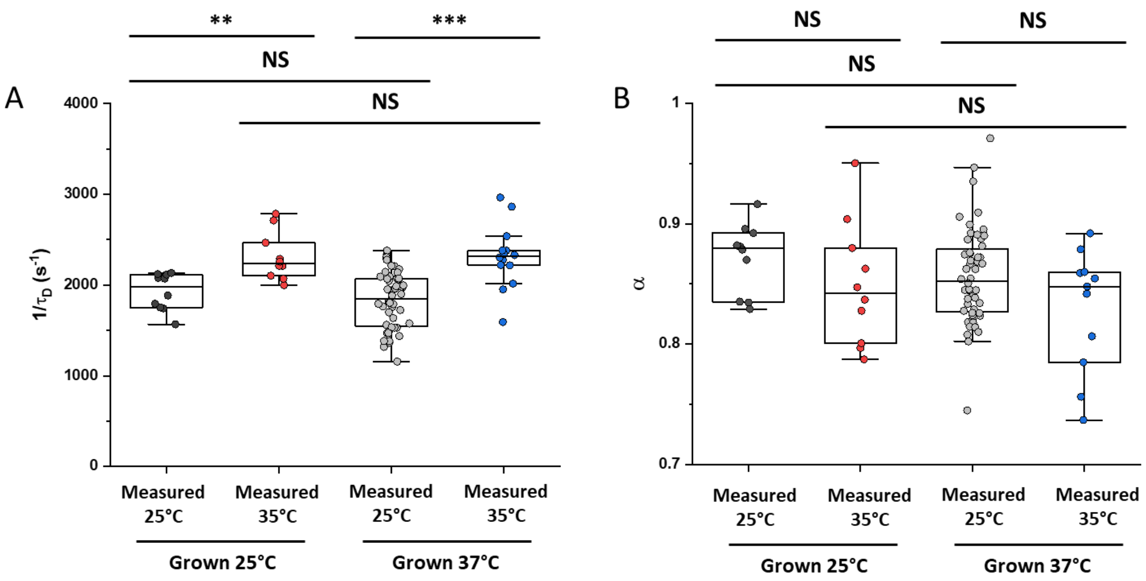


**Figure 5 – figure supplement 4. Effect of growth and measurement temperature on protein diffusion.** Mobility ($1/\tau_D$; A) and anomaly of diffusion ($\alpha$; B) for sfGFP in cells grown either at 25°C or at 37°C and measured either at 25°C or at 35°C, as in Figure 4A. *** p<0.0001; ** p<0.001; NS: no statistically significant difference in a two-tailed heteroscedastistic *t*-test.

**Figure 5 – figure supplement 4 – source data**

Individual values of $1/\tau_D$ from Figure 5 – figure supplement 4A.

Individual values of $\alpha$ from Figure 5 – figure supplement 4B.

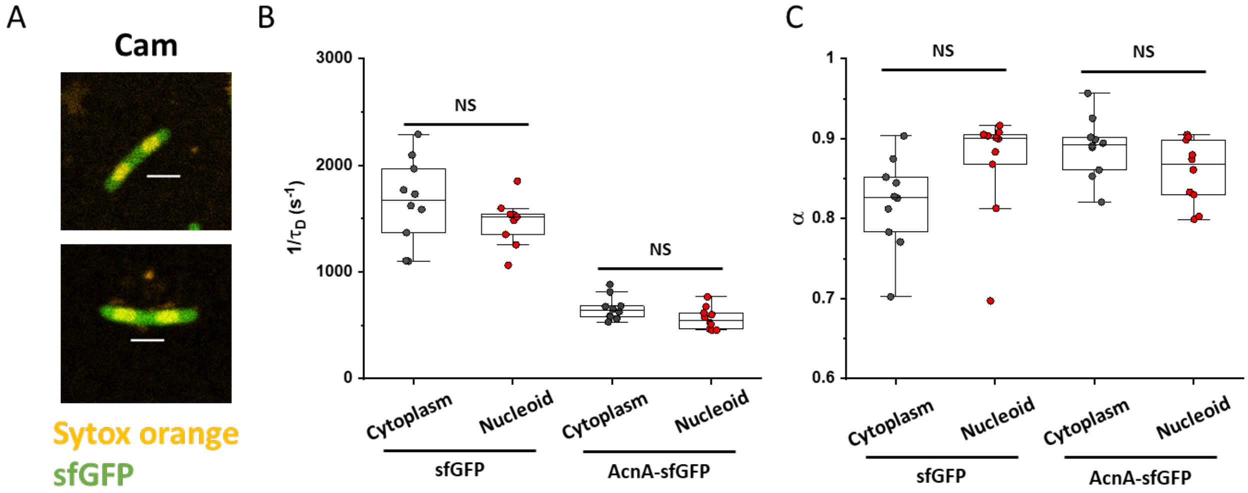


**Figure 5 – figure supplement 5. Influence of nucleoid on protein mobility.** (A) Bacterial cells were treated with chloramphenicol to achieve nucleoid compaction and stained with the DNA-binding dye SYTOX Orange. Scale bar is 2 µm. The mobility ($1/\tau_D$; B) and anomaly of diffusion (α; C) of sfGFP and of one larger construct (AcnA-sfGFP) was measured in both the cytoplasm and in the nucleoid of chloramphenicol treated cells. NS: no statistically significant difference in a two-tailed heteroscedastistic t-test.

**Figure 5 – figure supplement 5 – source data**

Individual values of $1/\tau_D$ from Figure 5 – figure supplement 5B.

Individual values of $\alpha$ from Figure 5 – figure supplement 5C.

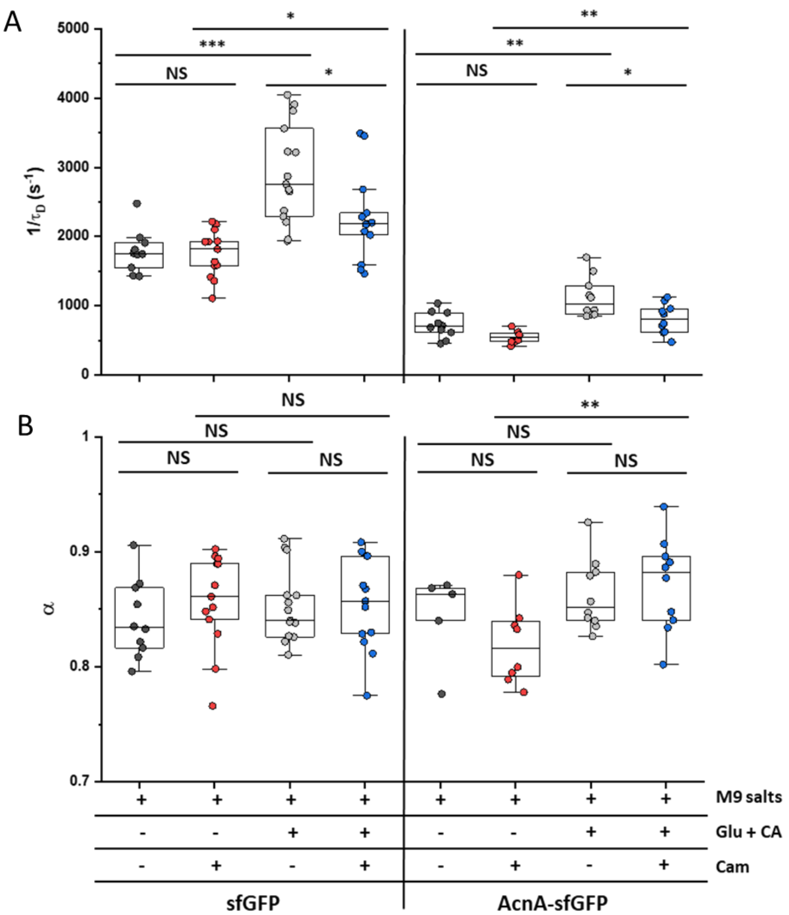


**Figure 5 – figure supplement 6. Effect of nutrient availability and growth on protein mobility.** The mobility ($1/\tau_D$; A) and anomaly of diffusion (α; B) of indicated protein constructs was measured in cells incubated at 35°C on agarose pads containing either only M9 salts or M9 salts together with 20 mM glucose and 0.2% casamino acids (Glu + CA). Where indicated, chloramphenicol (Cam) was also added to the agarose pads. *** $p<0.0001$; ** $p<0.001$; * $p<0.05$; NS: no statistically significant difference in a two-tailed heteroscedastistic *t*-test.

**Figure 5 – figure supplement 6 – source data**

Individual values of $1/\tau_D$ from Figure 5 – figure supplement 6A.

Individual values of $\alpha$ from Figure 5 – figure supplement 6B.

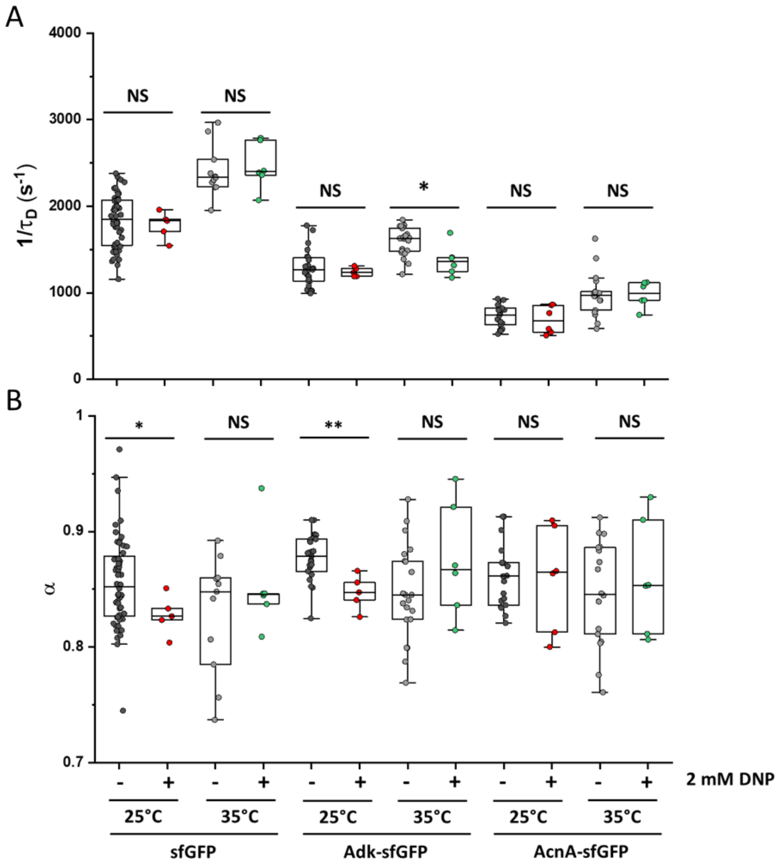


**Figure 5 – figure supplement 7. Effect of DNP on protein mobility.** The mobility ($1/\tau_D$; A) and the anomality of diffusion ($\alpha$; B) of sfGFP and of two constructs with higher molecular mass was measured in bacterial cells treated in batch for 60 minutes with 2 mM DNP and compared with the respective untreated control. Measurements were performed on 1% agarose pads prepared in tethering buffer and supplemented with 2 mM DNP at the indicated incubation temperature. ** $p<0.001$; * $p<0.05$; NS: no statistically significant difference in a two-tailed heteroscedastistic t-test.

**Figure 5 – figure supplement 7 – source data**

Individual values of $1/\tau_D$ from Figure 5 – figure supplement 7A.

Individual values of $\alpha$ from Figure 5 – figure supplement 7B.

### Appendix 1

#### Notes on the acquisition and analysis protocols for FCS measurements in bacterial cells

Due to the limited size of bacterial cells, FCS measurements require precise positioning of the confocal volume in the bacterial cytoplasm and minimization of the photobleaching-induced effects. In order to ensure that, before fitting an autocorrelation function, we verified the stability of the lateral (*xy*) positioning of the observation volume by visually analyzing for lateral drifts in confocal images acquired immediately before and after the FCS acquisition. This was done by annotating the *xy* position in the pre-aquisition image and verifying that the positioning did not change in the post-aquisition image after 120 seconds. Measurements showing *xy* drift were excluded from the analysis (Appendix 1 - figure 1). Furthermore, the focal stability of the sample was increased by thermal equilibration on the microscope stage before measurements.

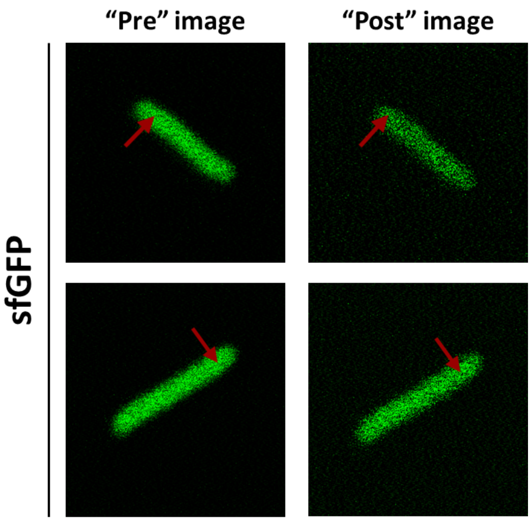


**Appendix 1 - figure 1. Typical examples of presence or absence of lateral focal drift during FCS measurements.** Substantial lateral drift could be observed for <10% of experiments (upper images), whereas most measurement showed no perceptible lateral drift (lower images).

Long-term photobleaching due to the progressive decrease of the total number of fluorescent proteins during FCS experiments (Appendix 1 - figure 2) is unavoidable due to the small volume of *E. coli* cells, and it requires correction to avoid artifacts. We observed that almost identical ACFs were obtained when correcting for the photobleaching using either multi-segment detrending (Jay Unruh, https://research.stowers.org/imagejplugins/index.html, Stowers Institute for Medical Research, USA) or a local averaging approach (Wachsmuth et al. 2015) (Appendix 1 - figure 3).

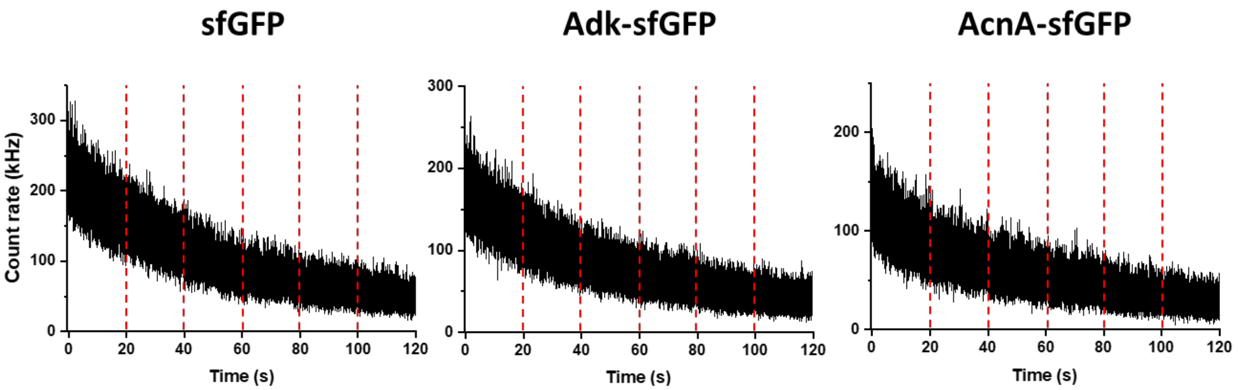


**Appendix 1 - figure 2. Typical traces of fluorescence intensity during FCS measurements.** Examples of fluorescence intensity traces for indicated protein fusions. The vertical red dashed lines separate sequential fluorescence intensity acquisitions on the same cell.

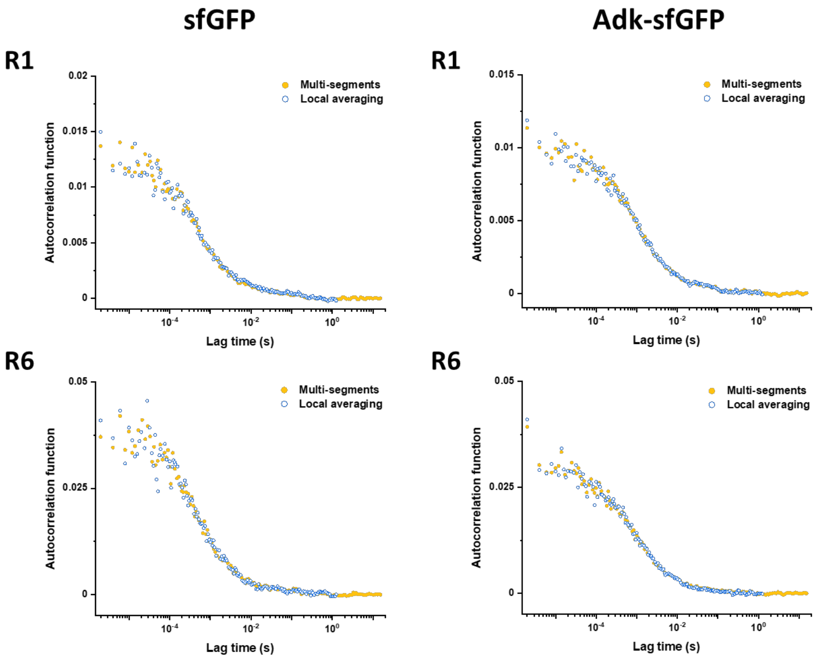


**Appendix 1 - figure 3. Results of detrending with multi-segments and local averaging approaches.** Comparison of experimental ACFs corrected using either multi-segments or local averaging approaches (as indicated) for sfGFP and Adk-sfGFP and different data acquisition segments (R1 vs R6).

We also confirmed that there was no systematic trend in the fitted values of $\tau_D$ and $\alpha$ with the time of the fluorescence trace acquisition (Appendix 1 - figure 4). An additional process that could potentially affect autocorrelation functions is short-term photobleaching of the fluorophore in the confocal volume, also known as cryptic photobleaching, which can artificially accelerate the decrease of the autocorrelation function and lead to an underestimation of the protein residence time (Macháň, Foo, and Wohland 2016). This process is different from long-term photobleaching, which is caused by the continuous illumination in the entire illumination light cone. However, the effect of cryptic photobleaching was shown to be typically <5%, even for proteins that diffuse 10 to 100 times slower and have higher bleaching rates than our constructs (Stasevich et al. 2010; Wachsmuth et al. 2015).

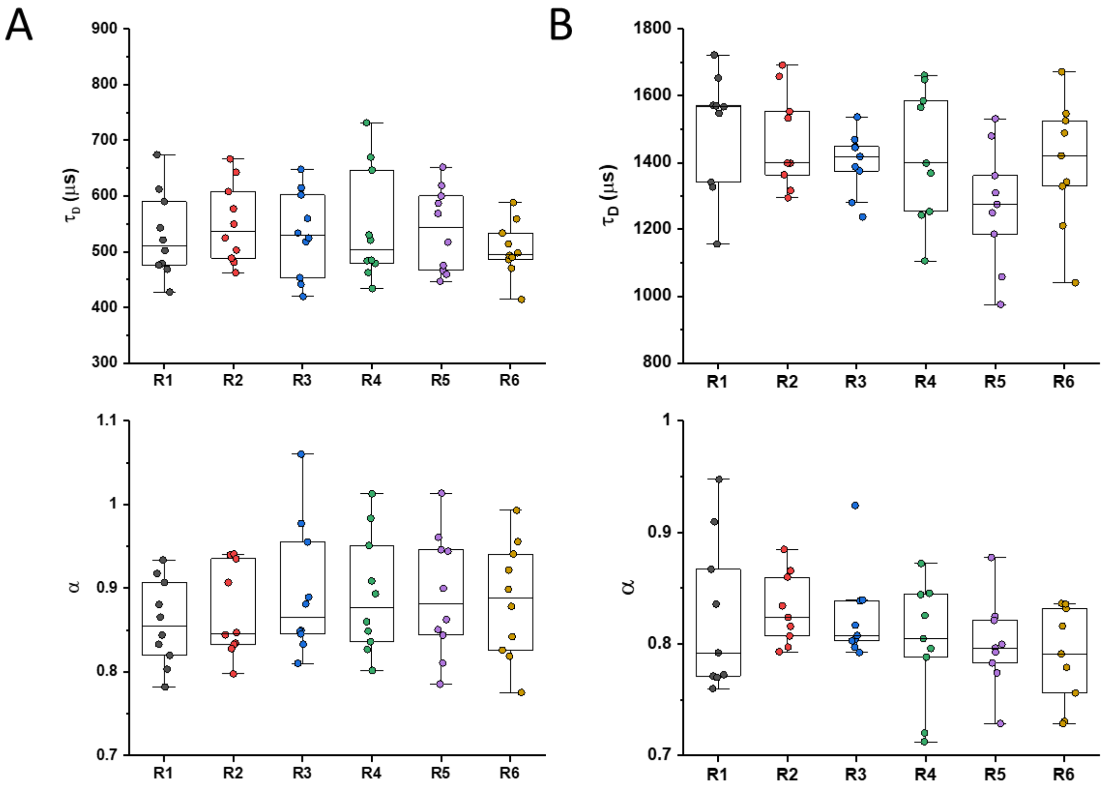


**Appendix 1 - figure 4. Values of $\tau_D$ or $\alpha$ for the six sequential ACFs.** Values were determined by fitting the anomalous diffusion model to experimental ACFs for the six sequential time segments per individual cell expressing sfGFP (A) or MetH-sfGFP (B)

### Appendix 2

#### Ornstein-Uhlenbeck model for confinement effect in FCS measurements

We aim to derive the autocorrelation function for an FCS experiment in which the fluorescent particles are confined by a (possibly anisotropic) harmonic potential centered at $(x, y, z) = (0,0,0)$, i.e.

$$V(x, y, z) = \frac{k_x x^2}{2} + \frac{k_y y^2}{2} + \frac{k_z z^2}{2}$$

where $k_i$ represents the stiffness of the potential in each dimension, and thus the extent $\sigma_i$ of confinement along that dimension given by $\sigma_i^2 \equiv k_\mathrm{B} T / k_i$.

We can treat each dimension independently, using $x$ without loss of generality in what follows. Diffusion in a harmonic potential is described by the Ornstein-Uhlenbeck process. The corresponding Green's function $P(x, t|x_0)$, representing the probability of finding a particle at position $x$ at time $t$ given that it was at position $x_0$ at time $t = 0$, is

$$P(x, t|x_0) = \frac{1}{\sqrt{2\pi\sigma_x^2 \left(1 - e^{-\frac{2Dt}{\sigma_x^2}}\right)}} \exp\left[-\frac{1}{2\sigma_x^2} \frac{\left(x - x_0 e^{-\frac{Dt}{\sigma_x^2}}\right)^2}{1 - e^{-\frac{2Dt}{\sigma_x^2}}}\right]$$

at long times, $t \to \infty$, we recover the stationary state given by the Boltzmann distribution corresponding to the harmonic trap

$$P_\mathrm{st}(x) = \frac{1}{\sqrt{2\pi\sigma_x^2}} \exp\left[-\frac{x^2}{2\sigma_x^2}\right].$$

The autocorrelation function (ACF) $G(t)$ of an FCS measurement is given as the multiple integral over the product of the probability to detect a photon from a molecule at some initial position $x_0$, the probability density that it diffuses from this position to a final position $x$ within time $t$ (given by Green's function), and the probability to detect a photon from a molecule at this final position (Enderlein et al. 2005; Enderlein 2012). Note that the probability of detection of a molecule will necessarily be proportional to the intensity of the laser beam, which we can assume Gaussian and also centered at $(x, y, z) = (0,0,0)$, with the usual form

$$I(x, y, z) = I_0 \exp\left(-\frac{2x^2}{\omega_0^2}\right) \exp\left(-\frac{2y^2}{\omega_0^2}\right) \exp\left(-\frac{2z^2}{S^2\omega_0^2}\right) \equiv I_0 I_x(x) I_y(y) I_z(z)$$

where $\omega_0$ is the width of the (circular) laser beam along the $x$ and $y$ directions, and $S$ is a dimensionless factor accounting for the anisotropy along the $z$ direction, i.e. the axial direction of the beam.

Ignoring constant normalization factors and baselines, the time dependent part of the autocorrelation function is then given by $G(t) = G_x(t)G_y(t)G_z(t)$ with

$$G_x(t) = \int \mathrm{d}x \int \mathrm{d}x_0 I_x(x) P(x, t|x_0) I_x(x_0) P_{\mathrm{st}}(x_0)$$

which can be directly integrated to give, after normalizing so that $G_x(t = 0) = 1$, the expression

$$G_x(t) = \left[1 + 2\frac{\sigma_x^2}{\omega_0^2} \frac{1 - e^{-\frac{2Dt}{\sigma_x^2}}}{1 + \frac{1}{8}\frac{\omega_0^2}{\sigma_x^2}}\right]^{-\frac{1}{2}}. \quad \text{(A2-1)}$$

Note that, in the limit of no confinement, $\sigma_x^2 \to \infty$, this equation reduces to the well-known ACF for unconfined diffusion

$$G_x(t) = \left[1 + \frac{4Dt}{\omega_0^2}\right]^{-\frac{1}{2}} = \left[1 + \frac{t}{\tau_D}\right]^{-\frac{1}{2}}$$

where we have defined the diffusion time $\tau_D \equiv \omega_0^2/(4D)$. With this definition, the ACF in Eq. (A2-1) can be rewritten as

$$G_x(t) = \left[1 + 2\frac{\sigma_x^2}{\omega_0^2}\frac{1 - e^{-\frac{1}{2}\frac{\omega_0^2}{\sigma_x^2}\frac{t}{\tau_D}}}{1 + \frac{1}{8}\frac{\omega_0^2}{\sigma_x^2}}\right]^{-\frac{1}{2}}.$$

The full three-dimensional ACF is then, in general,

$$G(t) = \left[1 + 2\frac{\sigma_x^2}{\omega_0^2}\frac{1 - e^{-\frac{1}{2}\frac{\omega_0^2}{\sigma_x^2}\frac{t}{\tau_D}}}{1 + \frac{1}{8}\frac{\omega_0^2}{\sigma_x^2}}\right]^{-\frac{1}{2}}\left[1 + 2\frac{\sigma_y^2}{\omega_0^2}\frac{1 - e^{-\frac{1}{2}\frac{\omega_0^2}{\sigma_y^2}\frac{t}{\tau_D}}}{1 + \frac{1}{8}\frac{\omega_0^2}{\sigma_y^2}}\right]^{-\frac{1}{2}}\left[1 + 2\frac{\sigma_z^2}{S^2\omega_0^2}\frac{1 - e^{-\frac{1}{2}\frac{\omega_0^2}{\sigma_z^2}\frac{t}{\tau_D}}}{1 + \frac{1}{8}\frac{S^2\omega_0^2}{\sigma_z^2}}\right]^{-\frac{1}{2}}.$$

The cylindrical geometry of a bacterium can be approximated by an infinite cylinder along the $y$ direction, so that $\sigma_x = \sigma_z = \sigma$ and $\sigma_y \to \infty$, resulting in the ACF

$$G(t) = \left[1 + 2\frac{\sigma^2}{\omega_0^2}\frac{1-e^{-\frac{1}{2}\frac{\omega_0^2}{\sigma^2}\frac{t}{\tau_D}}}{1+\frac{1}{8}\frac{\omega_0^2}{\sigma^2}}\right]^{-\frac{1}{2}}\left[1 + \frac{t}{\tau_D}\right]^{-\frac{1}{2}}\left[1 + 2\frac{\sigma^2}{S^2\omega_0^2}\frac{1-e^{-\frac{1}{2}\frac{\omega_0^2}{\sigma^2}\frac{t}{\tau_D}}}{1+\frac{1}{8}\frac{S^2\omega_0^2}{\sigma^2}}\right]^{-\frac{1}{2}}. \quad \text{(A2-2)}$$

Eq. (A2-2), with the added baseline and multiplicative correction accounting for particles in the non-fluorescent state, corresponds to Eq. (3) in the main text.

### Appendix 3

#### Effective diffusion coefficient of two linked proteins

In previous work, we studied the diffusion of two spherical objects with radii $a_1$ and $a_2$, joined together by a flexible linker (Agudo-Canalejo and Golestanian 2020). In the limit of a rigid linker of length $\ell$, the effective diffusion coefficient of the composite object goes as:

$$D \simeq \frac{k_B T}{6\pi\eta a_1} \frac{a_1}{(a_1+a_2)} \left[1 + 2\frac{a_1 a_2}{(a_1+a_2)(a_1+a_2+\ell)} - \frac{9}{8}\frac{a_1 a_2 (a_1-a_2)^2}{(a_1+a_2)^2(a_1+a_2+\ell)^2}\right] \quad \text{(A3-1)}$$

plus higher order correction terms of order $O\left(\frac{a_i^3}{(a_1+a_2+\ell)^3}\right)$.

We can then consider what is the effective diffusion coefficient of two proteins that are linked to each other. For that, we first need to connect the molecular mass to the effective radius of the protein. If we identify subunit 1 with GFP, and subunit 2 with the protein attached to it, and we call $M_{\text{GFP}}$ the molecular mass of GFP and $M_{\text{tot}}$ the total molecular mass (sum of GFP and the protein), we expect relations of the form

$$a_1 = C M_{\text{GFP}}^{\beta}$$

$$a_2 = C (M_{\text{tot}} - M_{\text{GFP}})^{\beta}$$

where $C$ is a proportionality constant assumed to be typical for all proteins (which is of order 1, with values reported in the literature of about 0.65 (Smilgies and Folta-Stogniew 2015), when the mass is in kDa and the radius is in nm), and $\beta$ is the scaling exponent introduced in the main text. For the linker which is made of 6 amino-acids, we may use the typical conversion factor 0.35 nm/amino-acid to estimate $\ell \approx 2$ nm.

Plugging these expressions for $a_1$ and $a_2$ into Eq. (A3-1) above, one obtains an expression for the diffusion coefficient $D$ as a function of $M_{\mathrm{tot}}$ that depends only on three parameters (since the molecular mass of GFP is known): (i) the diffusion coefficient of GFP $\frac{k_{\mathrm{B}}T}{6\pi\eta a_1}$, (ii) the exponent $\beta$, and (iii) the rescaled linker length $\ell/C$.

### Appendix 4

#### Exact p-values for all significance analysis

**Figure 1C**

| *Testing pair* | *P-value* |
|---|---|
| sfGFP vs Adk-sfGFP | 0.000000010 |
| Adk-sfGFP vs AcnA-sfGFP | 0.00000000044 |

**Figure 1 – figure supplement 10A**

| *Testing pair* | *P-value* |
|---|---|
| ClpS$^{WT}$-sfGFP vs ClpS$^{D35A_D36A_H66A}$-sfGFP | 0.000094 |
| Map$^{WT}$-sfGFP vs Map$^{Lys\rightarrow Ala}$-sfGFP | 0.0000012 |
| Map$^{WT}$-sfGFP vs Map$^{Lys\rightarrow Glu}$-sfGFP | 0.000000016 |
| Map$^{Lys\rightarrow Ala}$-sfGFP vs Map$^{Lys\rightarrow Glu}$-sfGFP | 0.10 |
| DnaK$^{WT}$-sfGFP vs DnaK$^{V436F}$-sfGFP | 0.00023 |

**Figure 1 – figure supplement 10B**

| *Testing pair* | *P-value* |
|---|---|
| ClpS$^{WT}$-sfGFP vs ClpS$^{D35A_D36A_H66A}$-sfGFP | 0.000014 |
| Map$^{WT}$-sfGFP vs Map$^{Lys\rightarrow Ala}$-sfGFP | 0.0092 |
| Map$^{WT}$-sfGFP vs Map$^{Lys\rightarrow Glu}$-sfGFP | 0.065 |
| Map$^{Lys\rightarrow Ala}$-sfGFP vs Map$^{Lys\rightarrow Glu}$-sfGFP | 0.37 |
| DnaK$^{WT}$-sfGFP vs DnaK$^{V436F}$-sfGFP | 0.00000019 |

**Figure 2C**

| *Testing pair* | *P-value* |
|---|---|
| Untreated vs A22 treatment | 0.0000000000007 |

**Figure 2D**

| *Testing pair* | *P-value* |
|---|---|
| Untreated vs A22 treatment | 0.000001 |

**Figure 2 – figure supplement 1**

| *Testing pair* | *P-value* |
|---|---|
| Untreated vs A22 treatment | 0.002 |

**Figure 2 – figure supplement 3A**

| *Testing pair* | *P-value* |
|---|---|
| sfGFP, 1 A.U. vs 0.66 A.U. | 0.00001 |
| AcnA-sfGFP, 1 A.U. vs 0.66 A.U. | 0.000002 |

**Figure 2 – figure supplement 3B**

| *Testing pair* | *P-value* |
|---|---|
| sfGFP, 1 A.U. vs 0.66 A.U. | 0.24 |
| AcnA-sfGFP, 1 A.U. vs 0.66 A.U. | 0.002 |

**Figure 4 – figure supplement 1A**

| *Testing pair* | *P-value* |
|---|---|
| Adk$^{E.c.}$-sfGFP vs Adk$^{Y.e.}$-sfGFP | 0.17 |
| Adk$^{E.c.}$-sfGFP vs Adk$^{V.c.}$ -sfGFP | 0.056 |
| Adk$^{E.c.}$-sfGFP vs Adk$^{C.c.}$-sfGFP | 0.93 |
| Adk$^{E.c.}$-sfGFP vs Adk$^{M.x.}$-sfGFP | 0.21 |
| Adk$^{E.c.}$-sfGFP vs Adk$^{B.s.}$-sfGFP | 0.23 |
| Pgk$^{E.c.}$-sfGFP vs Pgk$^{V.c.}$-sfGFP | 0.75 |
| Pgk$^{E.c.}$-sfGFP vs Pgk$^{C.c.}$-sfGFP | 0.26 |
| Pgk$^{E.c.}$-sfGFP vs Pgk$^{M.x.}$-sfGFP | 0.33 |

| | |
|---|---|
| AcnA$^{E.c.}$-sfGFP vs AcnA$^{Y.e.}$-sfGFP | 0.093 |
| AcnA$^{E.c.}$-sfGFP vs AcnA$^{V.c.}$-sfGFP | 0.084 |
| AcnA$^{E.c.}$-sfGFP vs AcnA$^{M.x.}$-sfGFP | 0.0000000023 |
| AcnA$^{E.c.}$-sfGFP vs AcnA$^{B.s.}$-sfGFP | 0.069 |

**Figure 4 – figure supplement 1B**

| ***Testing pair*** | ***P-value*** |
|---|---|
| Adk$^{E.c.}$-sfGFP vs Adk$^{Y.e.}$-sfGFP | 0.000060 |
| Adk$^{E.c.}$-sfGFP vs Adk$^{V.c.}$ -sfGFP | 0.18 |
| Adk$^{E.c.}$-sfGFP vs Adk$^{C.c.}$-sfGFP | 0.042 |
| Adk$^{E.c.}$-sfGFP vs Adk$^{M.x.}$-sfGFP | 0.070 |
| Adk$^{E.c.}$-sfGFP vs Adk$^{B.s.}$-sfGFP | 0.0029 |
| Pgk$^{E.c.}$-sfGFP vs Pgk$^{V.c.}$-sfGFP | 0.11 |
| Pgk$^{E.c.}$-sfGFP vs Pgk$^{C.c.}$-sfGFP | 0.0082 |
| Pgk$^{E.c.}$-sfGFP vs Pgk$^{M.x.}$-sfGFP | 0.0087 |
| AcnA$^{E.c.}$-sfGFP vs AcnA$^{Y.e.}$-sfGFP | 0.035 |
| AcnA$^{E.c.}$-sfGFP vs AcnA$^{V.c.}$-sfGFP | 0.16 |
| AcnA$^{E.c.}$-sfGFP vs AcnA$^{M.x.}$-sfGFP | 0.083 |
| AcnA$^{E.c.}$-sfGFP vs AcnA$^{B.s.}$-sfGFP | 0.00024 |

**Figure 5 – figure supplement 2A**

| ***Testing pair*** | ***P-value*** |
|---|---|
| sfGFP ionic strength 105 mM vs 305 mM | 0.0000036 |
| Adk-sfGFP ionic strength 105 mM vs 305 mM | 0.000044 |
| AroA-sfGFP ionic strength 105 mM vs 305 mM | 0.035 |
| AcnA-sfGFP ionic strength 105 mM vs 305 mM | 0.0018 |

**Figure 5 – figure supplement 2B**

| *Testing pair* | *P-value* |
|---|---|
| sfGFP 25°C vs 35°C | 0.00015 |
| Adk-sfGFP 25°C vs 35°C | 0.0000025 |
| AcnA-sfGFP 25°C vs 35°C | 0.00077 |

**Figure 5 – figure supplement 2C**

| *Testing pair* | *P-value* |
|---|---|
| sfGFP Untreated vs Chloramphenicol | 0.40 |
| sfGFP DMSO vs Rifampicin | 0.012 |
| Adk-sfGFP Untreated vs Chloramphenicol | 0.12 |
| Adk-sfGFP DMSO vs Rifampicin | 0.00048 |
| Pgk-sfGFP Untreated vs Chloramphenicol | 0.17 |
| Pgk-sfGFP DMSO vs Rifampicin | 0.0000012 |
| AcnA-sfGFP Untreated vs Chloramphenicol | 0.000011 |
| AcnA-sfGFP DMSO vs Rifampicin | 0.0085 |

**Figure 5 – figure supplement 2D**

| *Testing pair* | *P-value* |
|---|---|
| sfGFP M9 salts vs M9 salts, Glu+CA | 0.000023 |
| Adk-sfGFP M9 salts vs M9 salts, Glu+CA | 0.000070 |
| AroA-sfGFP M9 salts vs M9 salts, Glu+CA | 0.00000015 |
| AcnA-sfGFP M9 salts vs M9 salts, Glu+CA | 0.0022 |

**Figure 5 – figure supplement 3A**

| *Testing pair* | *P-value* |
|---|---|
| sfGFP ionic strength 105 mM vs 305 mM | 0.097 |
| Adk-sfGFP ionic strength 105 mM vs 305 mM | 0.54 |
| AroA-sfGFP ionic strength 105 mM vs 305 mM | 0.077 |

| | |
|---|---|
| AcnA-sfGFP ionic strength 105 mM vs 305 mM | 0.31 |

**Figure 5 – figure supplement 3B**

| *Testing pair* | *P-value* |
|---|---|
| sfGFP 25°C vs 35°C | 0.12 |
| Adk-sfGFP 25°C vs 35°C | 0.005 |
| AcnA-sfGFP 25°C vs 35°C | 0.26 |

**Figure 5 – figure supplement 3C**

| *Testing pair* | *P-value* |
|---|---|
| sfGFP Untreated vs Chloramphenicol | 0.32 |
| sfGFP DMSO vs Rifampicin | 0.50 |
| Adk-sfGFP Untreated vs Chloramphenicol | 0.53 |
| Adk-sfGFP DMSO vs Rifampicin | 0.32 |
| Pgk-sfGFP Untreated vs Chloramphenicol | 0.17 |
| Pgk-sfGFP DMSO vs Rifampicin | 0.59 |
| AcnA-sfGFP Untreated vs Chloramphenicol | 0.008 |
| AcnA-sfGFP DMSO vs Rifampicin | 0.42 |

**Figure 5 – figure supplement 3D**

| *Testing pair* | *P-value* |
|---|---|
| sfGFP M9 salts vs M9 salts, Glu+CA | 0.44 |
| Adk-sfGFP M9 salts vs M9 salts, Glu+CA | 0.035 |
| AroA-sfGFP M9 salts vs M9 salts, Glu+CA | 0.69 |
| AcnA-sfGFP M9 salts vs M9 salts, Glu+CA | 0.31 |

**Figure 5 – figure supplement 4A**

| *Testing pair* | *P-value* |
|---|---|
| sfGFP Grown 25°C, measured 25°C vs 35°C | 0.0020 |
| sfGFP Measured 25°C, grown 25°C vs 37°C | 0.060 |
| sfGFP Measured 35°C, grown 25°C vs 37°C | 0.98 |
| sfGFP Grown 37°C, measured 25°C vs 35°C | 0.000040 |

**Figure 5 – figure supplement 4B**

| *Testing pair* | *P-value* |
|---|---|
| sfGFP Grown 25°C, measured 25°C vs 35°C | 0.26 |
| sfGFP Measured 25°C, grown 25°C vs 37°C | 0.45 |
| sfGFP Measured 35°C, grown 25°C vs 37°C | 0.44 |
| sfGFP Grown 37°C, measured 25°C vs 35°C | 0.12 |

**Figure 5 – figure supplement 5A**

| *Testing pair* | *P-value* |
|---|---|
| sfGFP cytoplasm vs nucleoid | 0.20 |
| AcnA-sfGFP cytoplasm vs nucleoid | 0.062 |

**Figure 5 – figure supplement 5B**

| *Testing pair* | *P-value* |
|---|---|
| sfGFP cytoplasm vs nucleoid | 0.09 |
| AcnA-sfGFP cytoplasm vs nucleoid | 0.09 |

**Figure 5 – figure supplement 6A**

| *Testing pair* | *P-value* |
|---|---|
| sfGFP M9 salts vs M9 salts, Cam | 0.82 |
| sfGFP M9 salts vs M9 salts, Glu+CA | 0.000023 |
| sfGFP M9 salts, Cam vs M9 salts, Cam, Glu+CA | 0.019 |
| sfGFP M9 salts, Glu+CA vs M9 salts, Cam, Glu+CA | 0.019 |

| | |
|---|---|
| AcnA-sfGFP M9 salts vs M9 salts, Cam | 0.37 |
| AcnA-sfGFP M9 salts vs M9 salts, Glu+CA | 0.0022 |
| AcnA-sfGFP M9 salts, Cam vs M9 salts, Cam, Glu+CA | 0.0035 |
| AcnA-sfGFP M9 salts, Glu+CA vs M9 salts, Cam, Glu+CA | 0.014 |

**Figure 5 – figure supplement 6B**

| ***Testing pair*** | ***P-value*** |
|---|---|
| sfGFP M9 salts vs M9 salts, Cam | 0.33 |
| sfGFP M9 salts vs M9 salts, Glu+CA | 0.44 |
| sfGFP M9 salts, Cam vs M9 salts, Cam, Glu+CA | 0.91 |
| sfGFP M9 salts, Glu+CA vs M9 salts, Cam, Glu+CA | 0.80 |
| AcnA-sfGFP M9 salts vs M9 salts, Cam | 0.099 |
| AcnA-sfGFP M9 salts vs M9 salts, Glu+CA | 0.31 |
| AcnA-sfGFP M9 salts, Cam vs M9 salts, Cam, Glu+CA | 0.0085 |
| AcnA-sfGFP M9 salts, Glu+CA vs M9 salts, Cam, Glu+CA | 0.56 |

**Figure 5 – figure supplement 7A**

| ***Testing pair*** | ***P-value*** |
|---|---|
| sfGFP untreated vs DNP treatment, 25°C | 0.52 |
| sfGFP untreated vs DNP treatment, 35°C | 0.66 |
| Adk-sfGFP untreated vs DNP treatment, 25°C | 0.46 |
| Adk-sfGFP untreated vs DNP treatment, 35°C | 0.03 |
| AcnA-sfGFP untreated vs DNP treatment, 25°C | 0.59 |
| AcnA-sfGFP untreated vs DNP treatment, 35°C | 0.98 |

**Figure 5 – figure supplement 7B**

| *Testing pair* | *P-value* |
|---|---|
| sfGFP untreated vs DNP treatment, 25°C | 0.013 |
| sfGFP untreated vs DNP treatment, 35°C | 0.32 |
| Adk-sfGFP untreated vs DNP treatment, 25°C | 0.006 |
| Adk-sfGFP untreated vs DNP treatment, 35°C | 0.25 |
| AcnA-sfGFP untreated vs DNP treatment, 25°C | 0.94 |
| AcnA-sfGFP untreated vs DNP treatment, 35°C | 0.57 |

### Appendix 5

#### Numerosity of the constructs and conditions for each experiment

**Table 1. Figure 1 and Figure 1 – figure supplement 5**

| *Construct* | *Numerosity (n)* | *Construct* | *Numerosity (n)* |
|---|---|---|---|
| sfGFP | 52 | EntC-sfGFP | 15 |
| YggX-sfGFP | 8 | AroA-sfGFP | 9 |
| ClpS$^{WT}$-sfGFP | 11 | ThrC-sfGFP | 14 |
| FolK-sfGFP | 8 | MurF-sfGFP | 7 |
| Crr-sfGFP | 14 | DsdA-sfGFP | 14 |
| UbiC-sfGFP | 14 | HemN-sfGFP | 13 |
| CoaE-sfGFP | 11 | PrpD-sfGFP | 12 |
| Adk-sfGFP | 23 | DnaK$^{WT}$-sfGFP | 10 |
| Cmk-sfGFP | 16 | MalZ-sfGFP | 9 |
| KdsB-sfGFP | 22 | GlcB-sfGFP | 16 |
| Map$^{WT}$-sfGFP | 20 | MetE-sfGFP | 8 |
| MmuM-sfGFP | 14 | LeuS-sfGFP | 14 |
| PanE-sfGFP | 18 | AcnA-sfGFP | 19 |
| SolA-sfGFP | 7 | MetH-sfGFP | 9 |
| Pgk-sfGFP | 16 | | |

**Table 2. Figure 1 -figure supplement 7**

| *Construct* | *Numerosity (n)* |
|---|---|
| sfGFP | Same as table 1 |

**Table 3. Figure 1 -figure supplement 8**

| *Condition* | *Numerosity (n)* |
|---|---|
| Untreated | 5 |

| | |
|---|---|
| Cephalexin | 6 |

**Table 4. Figure 1 -figure supplement 10**

| ***Construct*** | ***Numerosity (n)*** |
|---|---|
| ClpS$^{WT}$-sfGFP | Same as table 1 |
| ClpS$^{D35A_D36A_H66A}$-sfGFP | 10 |
| Map$^{WT}$-sfGFP | Same as table 1 |
| Map$^{Lys\rightarrow Glu}$-sfGFP | 10 |
| Map$^{Lys\rightarrow Ala}$-sfGFP | 12 |
| DnaK$^{WT}$-sfGFP | Same as table 1 |
| DnaK$^{V436F}$-sfGFP | 10 |

**Table 5. Figure 2C, 2D and Figure 2 – figure supplement 1**

| ***Condition*** | ***Numerosity (n)*** |
|---|---|
| Untreated | Same as table 1 |
| A22-treated | 12 |

**Table 6. Figure 2 – figure supplement 3**

| ***Condition*** | ***Numerosity (n)*** |
|---|---|
| sfGFP 1 A.U. | Same as table 1 |
| sfGFP 0.66 A. U. | 12 |
| AcnA-sfGFP 1 A.U. | Same as table 1 |
| AcnA-sfGFP 0.66 A. U. | 10 |

**Table 7. Figure 3C**

| ***Construct*** | ***Numerosity (n)*** | ***Construct*** | ***Numerosity (n)*** |
|---|---|---|---|
| sfGFP | 11 | DsdA-sfGFP | 10 |

| | | | |
|---|---|---|---|
| YggX-sfGFP | 10 | GlcB-sfGFP | 10 |
| Adk-sfGFP | 16 | AcnA-sfGFP | 10 |
| PanE-sfGFP | 11 | MetH-sfGFP | 15 |

**Table 8. Figure 4 and Figure 4 – figure supplement 1**

| *Construct* | *Numerosity (n)* | *Construct* | *Numerosity (n)* |
|---|---|---|---|
| Adk$^{E.c.}$-sfGFP | Same as table 1 | Pgk$^{C.c.}$-sfGFP | 5 |
| Adk$^{Y.e.}$-sfGFP | 5 | Pgk$^{M.x.}$-sfGFP | 11 |
| Adk$^{V.c.}$-sfGFP | 10 | AcnA$^{E.c.}$-sfGFP | Same as table 1 |
| Adk$^{C.c.}$-sfGFP | 5 | AcnA$^{Y.e.}$-sfGFP | 5 |
| Adk$^{M.x.}$-sfGFP | 11 | AcnA$^{V.c.}$-sfGFP | 10 |
| Adk$^{B.s.}$-sfGFP | 10 | AcnA$^{M.x.}$-sfGFP | 10 |
| Pgk$^{E.c.}$-sfGFP | Same as table 1 | AcnA$^{B.s.}$-sfGFP | 10 |
| Pgk$^{V.c.}$-sfGFP | 10 | | |

**Table 9. Figure 5A, Figure 5 – figure supplement 1A, B, Figure 5 - figure supplement 2A and Figure 5 - figure supplement 3A**

| *Construct and condition* | *Numerosity (n)* | *Construct and condition* | *Numerosity (n)* |
|---|---|---|---|
| sfGFP, 105 mM | Same as table 1 | AroA-sfGFP, 105 mM | Same as table 1 |
| sfGFP, 305 mM | 11 | AroA-sfGFP, 305 mM | 12 |
| Adk-sfGFP, 105 mM | Same as table 1 | AcnA-sfGFP, 105 mM | Same as table 1 |
| Adk-sfGFP, 305 mM | 11 | AcnA-sfGFP, 305 mM | 6 |

**Table 10. Figure 5B, Figure 5 – figure supplement 1C, D, Figure 5 - figure supplement 2B and Figure 5 - figure supplement 3B**

| *Construct and condition* | *Numerosity (n)* | *Construct and condition* | *Numerosity (n)* |
|---|---|---|---|
| sfGFP, 25°C | Same as table 1 | AcnA-sfGFP, 25°C | Same as table 1 |

| | | | |
|---|---|---|---|
| sfGFP, 35°C | 14 | AcnA-sfGFP, 35°C | 18 |
| Adk-sfGFP, 25°C | Same as table 1 | | |
| Adk-sfGFP, 35°C | 21 | | |

**Table 11. Figure 5C, Figure 5 – figure supplement 1E, F, Figure 5 - figure supplement 2C and Figure 5 – figure supplement 3C**

| *Construct and condition* | *Numerosity (n)* | *Construct and condition* | *Numerosity (n)* |
|---|---|---|---|
| sfGFP, untreated | Same as table 1 | Pgk-sfGFP, untreated | Same as table 1 |
| sfGFP, chloramphenicol | 10 | Pgk-sfGFP, chloramphenicol | 10 |
| sfGFP, DMSO | 15 | Pgk-sfGFP, DMSO | 10 |
| sfGFP, rifampicin | 15 | Pgk-sfGFP, rifampicin | 10 |
| Adk-sfGFP, untreated | Same as table 1 | AcnA-sfGFP, untreated | Same as table 1 |
| Adk-sfGFP, chloramphenicol | 10 | AcnA-sfGFP, chloramphenicol | 10 |
| Adk-sfGFP, DMSO | 10 | AcnA-sfGFP, DMSO | 10 |
| Adk-sfGFP, rifampicin | 10 | AcnA-sfGFP, rifampicin | 10 |

**Table 12. Figure 5D, Figure 5 – figure supplement 1G, H, Figure 5 - figure supplement 2D and Figure 5 - figure supplement 3D**

| *Construct and condition* | *Numerosity (n)* | *Construct and condition* | *Numerosity (n)* |
|---|---|---|---|
| sfGFP, M9 salts | 10 | AroA-sfGFP, M9 salts | 11 |
| sfGFP, M9 salts, Glu+CA | 15 | AroA-sfGFP, M9 salts, Glu+CA | 11 |
| Adk-sfGFP, M9 salts | 11 | AcnA-sfGFP, M9 salts | 10 |
| Adk-sfGFP, M9 salts, Glu+CA | 12 | AcnA-sfGFP, M9 salts, Glu+CA | 10 |

**Table 13. Figure 5 – figure supplement 4**

| *Construct and condition* | *Numerosity (n)* |
|---|---|
| sfGFP, grown 25°C, measured 25°C | 10 |
| sfGFP, grown 25°C, measured 35°C | 10 |
| sfGFP, grown 37°C, measured 25°C | Same as table 1 |
| sfGFP, grown 37°C, measured 35°C | Same as table 10 |

**Table 14. Figure 5 – figure supplement 5**

| ***Construct and condition*** | ***Numerosity (n)*** |
|---|---|
| sfGFP, cytoplasm | 10 |
| sfGFP, nucleoid | 10 |
| AcnA-sfGFP, cytoplasm | 10 |
| AcnA-sfGFP, nucleoid | 10 |

**Table 15. Figure 5 – figure supplement 6**

| ***Construct and condition*** | ***Numerosity (n)*** | ***Construct and condition*** | ***Numerosity (n)*** |
|---|---|---|---|
| sfGFP, M9 salts | Same as table 12 | AcnA-sfGFP, M9 salts | Same as table 12 |
| sfGFP, M9 salts, Cam | 13 | AcnA-sfGFP, M9 salts, Cam | 8 |
| sfGFP, M9 salts, Glu+CA | Same as table 12 | AcnA-sfGFP, M9 salts, Glu+CA | Same as table 12 |
| sfGFP, M9 salts, Cam, Glu+CA | 13 | AcnA-sfGFP, M9 salts, Cam, Glu+CA | 10 |

**Table 16. Figure 5 – figure supplement 7**

| ***Construct and condition*** | ***Numerosity (n)*** |
|---|---|
| sfGFP, untreated, 25°C | Same as table 1 |
| sfGFP, 2 mM DNP, 25°C | 5 |
| sfGFP, untreated, 35°C | Same as table 10 |
| sfGFP, 2 mM DNP, 35°C | 6 |
| Adk-sfGFP, untreated, 25°C | Same as table 1 |
| Adk-sfGFP, 2 mM DNP, 25°C | 5 |
| Adk-sfGFP, untreated, 35°C | Same as table 10 |
| Adk-sfGFP, 2 mM DNP, 35°C | 6 |
| AcnA-sfGFP, untreated, 25°C | Same as table 1 |
| AcnA-sfGFP, 2 mM DNP, 25°C | 5 |
| AcnA-sfGFP, untreated, 35°C | Same as table 10 |
| AcnA-sfGFP, 2 mM DNP, 35°C | 6 |
